# The Ecology of Palm Genomes: Repeat-associated genome size expansion is constrained by aridity

**DOI:** 10.1101/2021.11.04.467295

**Authors:** Rowan J. Schley, Jaume Pellicer, Xue-Jun Ge, Craig Barrett, Sidonie Bellot, Maïté S. Guignard, Petr Novák, Jan Suda, Donald Fraser, William J. Baker, Steven Dodsworth, Jiří Macas, Andrew R. Leitch, Ilia J. Leitch

## Abstract

- Genome size varies 2,400-fold across plants, influencing their evolution through changes in cell size and cell division rates which impact plants’ environmental stress tolerance. Repetitive element expansion explains much genome size diversity, and the processes structuring repeat ‘communities’ are analogous to those structuring ecological communities. However, which environmental stressors influence repeat community dynamics has not yet been examined from an ecological perspective.
- We measured genome size and leveraged climatic data for 91% of genera within the ecologically diverse palm family (Arecaceae). We then generated genomic repeat profiles for 141 palm species, and analysed repeats using phylogenetically-informed linear models to explore relationships between repeat dynamics and environmental factors.
- We show that palm genome size and repeat ‘community’ composition are best explained by aridity. Specifically, *EnSpm CACTA* repeats were more abundant in palm species from wetter environments, which generally had larger genomes (>2.15Gbp/1C), suggesting amplification. In contrast, *Ty1-copia Angela* elements were more abundant in drier environments.
- Our results suggest water stress inhibits the expansion of repeats through selection on upper genome size limits. However, *Ty1-copia Angela* elements, which may associate with stress-response genes, have amplified in arid-adapted palm species. Overall, we provide novel evidence of climate influencing the assembly of repeat ‘communities’.

## Introduction

Repetitive elements (hereafter, ‘repeats’) constitute a large part of most eukaryotic genomes and are responsible for much of the 64,000-fold variation in genome sizes within eukaryotes (Hidalgo *et al*., 2017). Repeats have a major effect on genome size variation through expansion and deletion of elements (Novák *et al*., 2020a). Previous work suggests that genome size may impact fitness, with larger genomes being advantageous in certain environments but disadvantageous in others (e.g. Knight *et al*., 2005; Faizullah *et al*., 2021). This may arise through increased biochemical costs of maintaining larger genomes and cells (e.g. Kang *et al*., 2015; Guignard *et al*., 2017), changes to cell cycle times (Francis *et al*., 2008), and/or impacts on cell size (Doyle & Coate 2019), which can affect gas exchange (e.g. Franks & Beerling 2009), water use efficiency (e.g. Lawson & Blatt 2014; Simonin & Roddy 2018) and photosynthesis (Roddy *et al*., 2020). Repeats can also directly affect host fitness by the activation or repression of genes through insertion or deletion of elements into coding sequences or their regulatory regions (Casacuberta & González 2013; Lisch, 2013).

Several studies have investigated the link between genome size variation and repetitive element dynamics in plants. For example, in the legume tribe Fabeae (Macas *et al*., 2015) and *Hesperis* (Brassicaceae) (Hloušková *et al*., 2019), much of the diversity in genome size is derived from the expansion of certain repeat lineages. Similarly, other studies have explored relationships between genome size variation and environmental conditions. For example, in orchids, models of genome size divergence indicated different genome size optima for species adapted to contrasting habitats (e.g., terrestrial and epiphytic growth forms) and suggested associations between genome size and climatic conditions (e.g. precipitation and temperature) (Trávníček *et al*., 2019). However, few studies have linked the combined impact of repeat dynamics and the environment of the host in generating genome size diversity. In mangroves there is evidence of long terminal repeat (LTR) retrotransposon excision and associated genome downsizing across lineages which are convergently adapted to stressful intertidal environments (Lyu *et al*., 2018), although there remains the need to examine environmental factors explicitly. Whilst previous work suggests that there may be interactions between repeats, genome sizes and the environment, no study has yet integrated repeat and genome size dynamics with ecological factors across a plant family.

Genomes may be seen as ecosystems ‘populated’ by repeats, each of which interacts with other repeats, genes, regulatory sequences and the genomic machinery (e.g., nuclear components involved in recombination, replication and DNA repair (Venner *et al*., 2009; Stitzer *et al*., 2021)). As such, repeat dynamics can be considered from a community ecology perspective, where the host genome is analogous to an ecosystem, repeat lineages to species and copy numbers of a given repeat lineage to numbers of individuals. The similarities between repeat dynamics in genomes and community dynamics in ecosystems were highlighted by a review examining the ‘Ecology of the Genome’ (Brookfield, 2005), in which the author suggested that certain ecological model parameters effectively describe aspects of repeat dynamics. For example, much like carrying capacity *K* in community ecology, we can assume that organisms have a certain ‘carrying capacity’ for repeats in the genome. The carrying capacity for repeats (i.e., set by the upper limit of an organism’s genome size) will be defined by the cost of genomic maintenance and replication, as well as by any physical constraints that genome size poses on cell size, cell division rate and cell physiology (Hidalgo *et al*., 2017). We might predict that the dynamics (i.e. loss and amplification) of a given repeat will be influenced by this upper genome size limit, and by whether the amplification of any element is detrimental for the host (Novák *et al*., 2020a). In turn, the limit on a species’ genome size may itself be impacted by the environment in which the organism lives (as in Trávníček *et al*., 2019; Faizullah *et al*., 2021).

Furthermore, many quantitative aspects of repeat ‘communities’ (such as the richness of repeat lineages, their diversity and the amount of the genome they occupy) are directly analogous to such metrics which describe species composition in ecological communities. The similarities between genomes and ecosystems are summarised in Figure 1, including calculations for species (or repeat) richness (Menhinick’s index (Whittaker, 1977)) and diversity (Shannon’s index (Shannon, 1948)) for both an ecosystem and a genome. Despite the call for further exploration of genome dynamics using ecological methods (Brookfield, 2005; Mauricio, 2005; Venner *et al*., 2009) there remains little work dealing directly with this subject.

**Figure 1:**
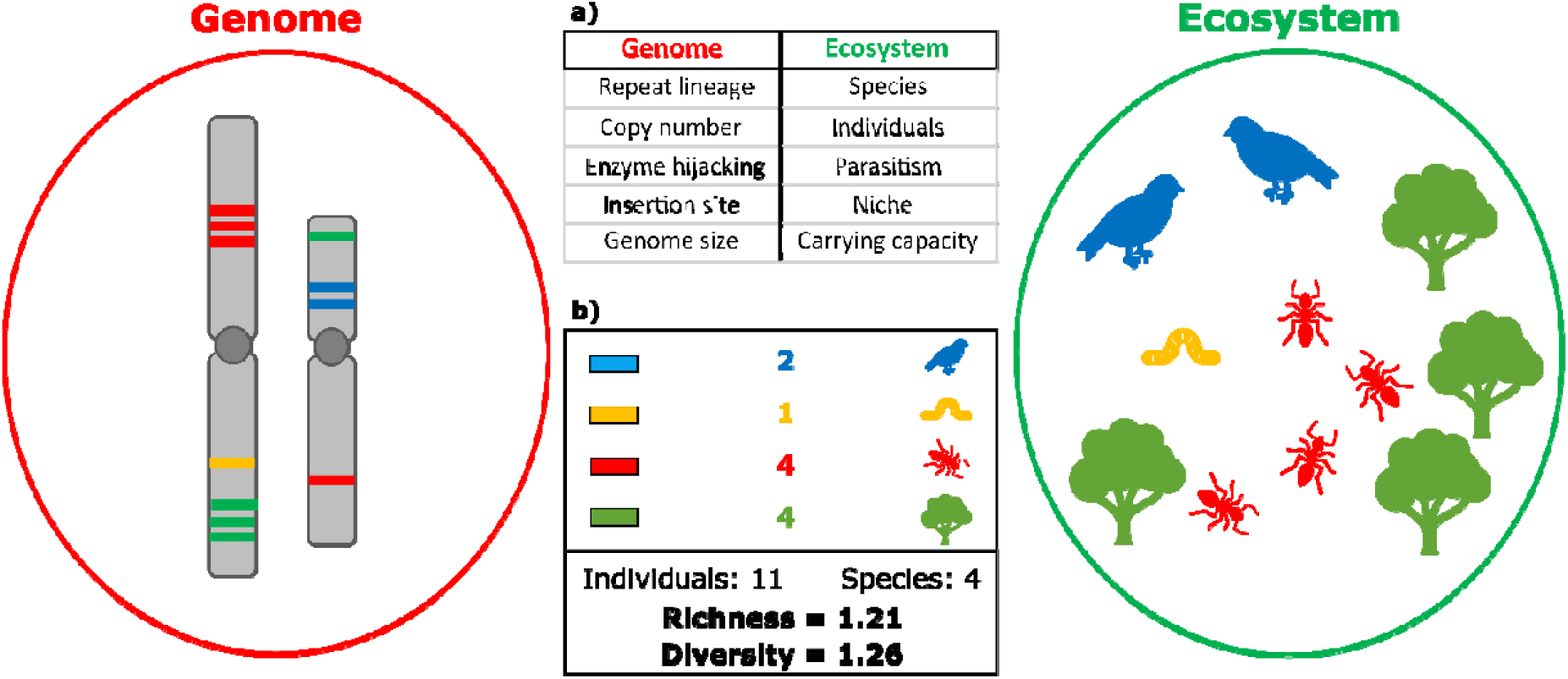
Summary of the similarities between repeats in a genome and organisms within an ecosystem. Repeat sequences are shown as bands along two chromosomes of a hypothetical species with n = 2 (left), with the colour of each band representing a specific repeat lineage. The number of bands with the same colour represent the copy number of that repeat lineage. Similarly, four species of organisms are shown in the simplified ecosystem (right), with the shape and colour of an icon representing the species, and the number of each icon representing the number of individuals of that species. **a)** Similarities between genomes and ecosystems. The copy number of repeats of a certain lineage in a genome is analogous to the number of individuals of a species in an ecosystem. Hijacking of replication machinery between repeat lineages occurs in genomes (Robillard *et al*. 2016), being analogous to parasitic interactions between species in an ecosystem. Insertion preferences in repeats (e.g., closer to centromeres) can represent an ecological ‘niche’ of that repeat (Venner *et al*. 2009), and the total carrying capacity (analogous to *K* in an ecosystem) is influenced by multiple factors, such as constraints on absolute genome size of the host species. **b)** Similarities between repeat lineages in genomes and species in ecosystems allow the use of similar descriptive metrics. In this simple example, there are 11 individuals (i.e., copies) in total belonging to four species (i.e., repeat lineages). Thus, Menhinick’s richness index and Shannon’s diversity index can be calculated for both genomes and ecosystems, giving values of 1.21 and 1.26 in the figure, respectively. See Methods S1 (Supporting Information 2) for the formulae used to calculate these indices.

One reason for the paucity of integrative studies focusing on repeat dynamics within the genome may have been the lack of suitable genomic data for non-model organisms. However, the advent of high-throughput sequencing techniques such as genome skimming (summarised in Dodsworth, 2015), which sequences DNA broadly across a genome, now permits the investigation of repeat dynamics in any eukaryote (Novák *et al*., 2020a). Thus, we are now in a position to explore the relationships between the ecology of the genome and the ecology of the species. An ideal study system for this is the palm family (Arecaceae), an iconic and economically important plant group which is a key element of tropical floras (Couvreur & Baker 2013). Palm species are adapted to a wide range of environments spanning extremes of water stress, from the aridity of the Sahara Desert to the perhumid rainforests of New Guinea (Dransfield *et al*., 2008; Kissling *et al*., 2012) and genome size varies 58-fold across palms (based on data for 121 species; Plant DNA C-values database, https://cvalues.science.kew.org/, Pellicer & Leitch (2020)). Moreover, chromosome numbers are available for many palm species and polyploidy is rare (thus far only reported in four out of the c. 2,600 species described (Röser, 1994; Röser *et al*., 1997; Dransfield *et al*., 2008)) despite evidence of an ancient whole-genome duplication at the base of the group (Barrett *et al*., 2019). This allows us to differentiate between genome size variation due to repeat dynamics from variation due to polyploidy. In addition, a wealth of other datasets exist for this important family, including trait data and distribution data (Kissling *et al*., 2019; WCVP, 2020).

Here, we harness the power of existing ecological and distribution datasets for palms, combining them with new genome size data for 437 species and high-throughput DNA sequencing data to explore whether and how environmental factors influence repeat dynamics and genome size. We analyse these data using an approach inspired by the community ecology literature, allowing us to closely examine the ‘ecology’ of repeat lineages in palm genomes to improve our understanding of how repeat dynamics and their effect on genome size may be influenced by past and present climate. Specifically, we explore (i) whether genome size variation within the palm family is explained by individual climatic factors, and (ii) whether differential expansion of specific repeats in different climates plays a role in generating this genome size diversity.

## Materials and Methods

### Plant material collection and genome size measurement

We collected 513 accessions from 437 of the ca. 2,600 palm species (19.7%), representing 165 out of 181 palm genera (91.1%) (Baker & Dransfield 2016), and all five subfamilies of the Arecaceae. Palm accessions were sampled from the living collections at the Royal Botanic Gardens, Kew (UK), Montgomery Botanical Center (US), Fairchild Tropical Botanic Garden (US), Prague Botanical Garden (CZ), Frankfurt Palmengarten (DE) and supplemented with field collections. Nuclear DNA contents were estimated by flow cytometry, following the one-step procedure (Pellicer *et al*., 2020). Where multiple genome size measurements were available for a species, we calculated a per-species mean genome size. The table of genome sizes and accession information can be found in Supporting Information 1, and details of sample collection and flow cytometry are shown in Methods S1 (Supporting Information 2). The phylogenetic spread of the genome size data we generated (n=437) as well as published data from the Plant DNA C-values database (https://cvalues.science.kew.org/, Pellicer & Leitch (2020), n=35), totalling 472 species, was visualised using the *plotTree*.*wBars()* function in *phytools* (Supporting Information 2: Fig. S1).

### Phylogenetic, environmental and genomic data collection

In order to provide a phylogenetic backbone to our study we used the phylogenetic tree for all palm species generated by Faurby *et al*. (2016). We then assembled a list of accepted species names from the World Checklist of Vascular Plants (WCVP, 2020) (as of January 2020) and updated species names across our datasets (including the phylogeny) to follow this list of accepted names.

Geographic occurrence data were collated from an existing palm distribution dataset which contained occurrence data from GBIF (www.gbif.com; dataset https://doi.org/10.15468/dd.at82kf) and from herbarium specimens (collected from K and L). To collect data from GBIF, all palm names published at that time (March 2018, from WCVP (2020)) were searched against the GBIF taxonomic backbone, and occurrences were retrieved for the 7,469 names that matched. Occurrences were then reconciled to a list of accepted palm names at the time (WCVP, 2020), and cleaned based on the GBIF coordinate issue flags and using the *R* (R Development Core Team, 2013) package *CoordinateCleaner* v1.0-7 (Zizka *et al*., 2019). We first corrected issues such as incorrect coordinate signs, and removed coordinates falling into maritime areas, city, province or country centroids. We also removed coordinates from biodiversity institutions, coordinates with zero values or those with an uncertainty > 100 km. Finally, we removed coordinates which were inconsistent with the country assignment of the record, as well as those falling outside the native distribution range of the species and those recorded before 1945. Duplicated occurrence records were also discarded.

Based on this refined occurrence dataset, we downloaded environmental data from *WorldClim* for all 472 species with genome size estimates using the *R* package *raster* (Hijmans & van Etten 2012). Data were extracted for each individual in the occurrence dataset for all palm species, comprising all nineteen bioclimatic variables from the *WorldClim* dataset, which detail biologically significant measures of temperature and precipitation (BIO1 to BIO19), as well as elevation data. From this we calculated a ‘per-species’ mean for each variable by averaging every value for all individuals of a species.

To examine repeat profiles of as many species as possible from across the palm family, we used genome skimming data from 141 accessions, representing 141 species from 88 palm genera and all subfamilies excluding the monospecific Nypoideae. Total DNA was extracted from silica-dried plant material using the CTAB method (Doyle & Doyle 1987), followed by library preparation using the NEBNext Ultra II library kit (New England BioLabs Ltd, Hitchin, UK). The final library pools were generated and sequenced on the Illumina X ten platform (Illumina, San Diego, USA) by the Beijing Genomics Institute (BGI, Shenzhen, China), generating 2×150 bp paired-end sequencing reads. Species-specific genome size data were not available for 63 of the 141 palm species sequenced, so for these we calculated a mean genome size estimate based on data for congeneric species given that genome size showed phylogenetic signal (see Results). A table of accessions and their voucher information is provided in Supporting Information 2: Table S1, and the phylogenetic spread of these data are shown in Supporting Information 2: Fig. S2, plotted using the *R* package *phytools* (Revell, 2012).

### Modelling relationships between genome size and environmental variables

To assess whether genome size variation within the palm family is correlated with environmental factors, we used a phylogenetically-informed linear modelling approach known as phylogenetic generalised least squares (PGLS) (Grafen, 1989) in the *R* package *caper* (Orme *et al*., 2013). We included all 472 palm species from our new genome size estimates (437 species) and the Plant DNA C-values database (35 species), comprising 165 genera across all five palm subfamilies.

First, the distribution of genome sizes was visualised using the *hist()* function in *R*, followed by superimposing the genome size data onto the palm phylogenetic tree using the *plotTree*.*wBars()* function in *phytools*. We then assessed the degree of phylogenetic signal in the genome size dataset using the λ value with the *phylosig*() function in *phytools*, and tested between the following models: a neutral model of trait evolution (Brownian motion), a model describing rapid diversification in trait values near the root of the tree (Early Burst) and a model describing evolution towards an optimal genome size values (Ornstein-Uhlenbeck) in *phytools*.

To assess how environmental variables may influence genome size variation, our PGLS analysis comprised six *WorldClim* bioclimatic variables and elevation as predictors, with a log-transformed response variable (genome size, 1C-values, measured in gigabase pairs (Gbp)) to improve normalcy. Our initial PGLS model was *log(Genome size) ∼ Isothermality + Precipitation of the Driest Month + Minimum Temperature of the coldest month + Precipitation of the Wettest Month + Precipitation Seasonality + Precipitation of the Coldest Quarter + Elevation*. Before beginning PGLS analysis, we explored autocorrelations between all 19 bioclimatic variables from WorldClim using the functions *corr*(), *heatmap*() and *cophylo()* in *R*. Predictors were chosen from all WorldClim variables to represent the highest level of temporal resolution (e.g., precipitation of the driest month rather than precipitation of the driest quarter). Perfectly autocorrelated predictors were identified and removed using the *alias()* function in *R*. Multicollinearity in the PGLS was evaluated with variance inflation factors (VIF), all of which were below 10, using the *vif*() function in the *car* package (Fox & Weisberg 2018).

For our initial PGLS model we used the lambda branch transformation in *caper* as this provided superior model fit when compared to the delta branch transformation, based on the corrected Akaike Information Criterion (AICc) of each model (Barton and Barton 2015), and phylogenetic covariance was estimated based on the Faurby *et al*. (2016) palm phylogenetic tree. The initial model was then reduced to the minimum adequate model in a stepwise fashion using *update()*, by removing explanatory variables with *P-*values >0.05 in the model summary. We also ran a model identical to that described above but excluding the four polyploid palm species (*Voanioala gerardii, Jubaeopsis caffra, Rhapis humilis* and *Arenga caudata*) to test whether model output was consistent without these polyploid taxa.

Given the unequal distribution of genome size across species, we performed further analyses following PGLS to establish how relationships between environmental variables and genome size vary across the range of values. Since the minimum adequate PGLS model indicated that ‘Precipitation of the Driest Month’ (i.e., aridity preference) was the most significant term explaining genome size variation, we split genome size and aridity preference into four quartile bins for each variable, assigned each palm species to one of these bins for each variable, and performed a chi-squared test in *R* (shown in Supporting Information 2: Table S2). These associations were then visualized using the R package *corrplot* (Wei *et al*., 2017).

### DNA repeat profiling

We quantified the amounts of different repeat lineages in 141 palm genomes, thereby generating a repeat profile for each species in the genome skimming dataset, using the *RepeatExplorer2* pipeline (Novák *et al*., 2013) and its published protocol (Novák *et al*., 2020b) on the James Hutton Institute’s Crop Diversity HPC. We prepared the genome skimming data for all available palm species by first qualityl□checking reads using *FASTQC* v0.11.3 (Andrews, 2010), following which *SOAPnuke* v1.5.6 (https://github.com/BGI-flexlab/SOAPnuke) was used to remove adapters, to remove reads with a Phred quality score <15 and to remove reads that contained >10% of unidentified (N) bases. Reads were subsequently trimmed to 100 bp using *Trimmomatic* v.0.3.6 (Bolger *et al*., 2014) as required by *RepeatExplorer2*. Following this we interleaved paired-end reads with *seqtk* v1.3-r106 (https://github.com/lh3/seqtk) (*-mergepe* flag), sampled reads relative to each species’ genome size to attain a 0.1x genome proportion across species (-*sample* flag) and transformed read files into FASTA format (-*seq* flag) for input to *RepeatExplorer2*. Genome proportion was calculated as *((Number of reads*Read length)/Genome size in base pairs)*, where a genome proportion of 0.1x is equal to 10% of the sampled genome. This proportion was used to include as many species as possible in the *RepeatExplorer2* analysis while having a sufficient genome proportion to ensure the repeat analyses were representative of each genome, based on previous studies (e.g. Macas *et al*., 2015).

We ran *RepeatExplorer2* ensuring that only clusters making up at least 0.05% of analysed reads were reported (*-m* 0.05) with a minimum overlap of 55 bp for reads to be assigned to clusters (*-o* 55) according to the developers (Novák *et al*., 2020b). This means *RepeatExplorer2* will detect both active repeats and inactive, degenerate repeats according to these thresholds. We used the VIRIDIPLANTAE3.0 database from REXdb (Neumann *et al*., 2019) as a reference for assigning clusters to different repeat lineages (*-tax* VIRIDIPLANTAE3.0). *RepeatExporer2* output cluster tables were then collated and processed using custom *BASH* and *R* scripts (Supporting Information 3), followed by manual correction of repeat annotations as described in the *RepeatExplorer2* protocol (Novák *et al*., 2020b).

### Assessing repeat dynamics in palm genomes

We used three different metrics from community ecology to describe repeat compositions in an analogous fashion to species compositions of ecological communities. We defined repeat groups (hereafter ‘lineages’) based on the lowest hierarchical classification for each lineage in the REXdb plant repeat database, in which the classifications are based mainly on similarities in conserved polyprotein regions, along with structural and sequence variation (Neumann *et al*., 2019). The classification of these repeat lineages, along with how we defined them, is given in Supporting Information 2: Table S3.

We first calculated the richness (Menhinick’s Index), total occupancy and diversity (Shannon’s Index) of repeats to provide three ecological summary metrics for each palm genome’s repeat ‘community’, where a repeat lineage in a genome is analogous to a species in an ecosystem. We then tested whether there were significant relationships between each of these ecological metrics and aridity preferences, using genome size as an interaction term in PGLS. Using with the Faurby *et al*. palm dendrogram as a covariate, our initial model for repeat richness was *log(Repeat richness) ∼ Precipitation of the Driest Month*Genome size*Total repeat occupancy*. For total repeat occupancy and repeat diversity the same model was used, except the responses were not logged because these two metrics were normally distributed. Non-significant terms were removed and models were compared using AICc.

We performed PGLS regression in *caper* to test whether the amount of the genome (in gigabase pairs, Gbp/1C) occupied by certain repeat lineages was correlated with genome size, and hence whether differential expansion or reduction of specific repeat lineages was responsible for genome size diversity in palms. The initial model used for this was *log(Genome size) ∼ (All repeat lineages)*, where ‘*(All repeat lineages)’* indicates the amount of each species’ genome occupied by each repeat lineage. The amounts of the genome occupied by each repeat lineage were thus used as separate predictor variables. Following this, non-significant terms were removed using *update()*, leaving the minimum adequate model.

Finally, to infer whether certain repeat lineages expand or contract preferentially under different precipitation regimes, we used PGLS to assess if differences in aridity preference (precipitation of the driest month) among species were explained by differences in the amount of the genome occupied by different repeat lineages (in Gbp/1C). The initial model used for this was *sqrt(Precipitation of the driest month) ∼ Genome size class:(All repeat lineages)*, using ‘large’ (> 2.15 Gbp/1C median value for the species in the genome skimming dataset) and ‘small’ (≤ 2.15 Gbp/1C median value) genome size bins as interaction terms since genome size and repeat amount often have an asymptotic relationship (Novák *et al*., 2020a). Again, model reduction was carried out to retain the minimum adequate model.

## Results

### Palm genome size variation

Combining the new genome size data estimated here (437 species) with previously published data taken from the Plant DNA C-values database (35 species) did not extend the previously reported 58-fold range of palm genome sizes, but greatly expanded the taxonomic breadth of sampling. Genome size ranged from 0.53 Gbp/1C in *Licuala orbicularis* and *Licuala sarawakensis* to 15.40 Gbp/1C in the presumed diploid *Pinanga sessilifolia* (based on chromosome counts of 2n = 32 in related species) and 30.63 Gbp/1C in the polyploid *Voanioala gerardii* (2n = 596 (Johnson *et al*., 1989; see also Röser, 1994)). The mean genome size across palms was 3.70 Gbp/1C (*s*.*d*. = 3.175), with a median value of 2.67 Gbp/1C. Genome sizes are displayed on the phylogenetic tree of the palm family in Figure 2.

**Figure 2:**
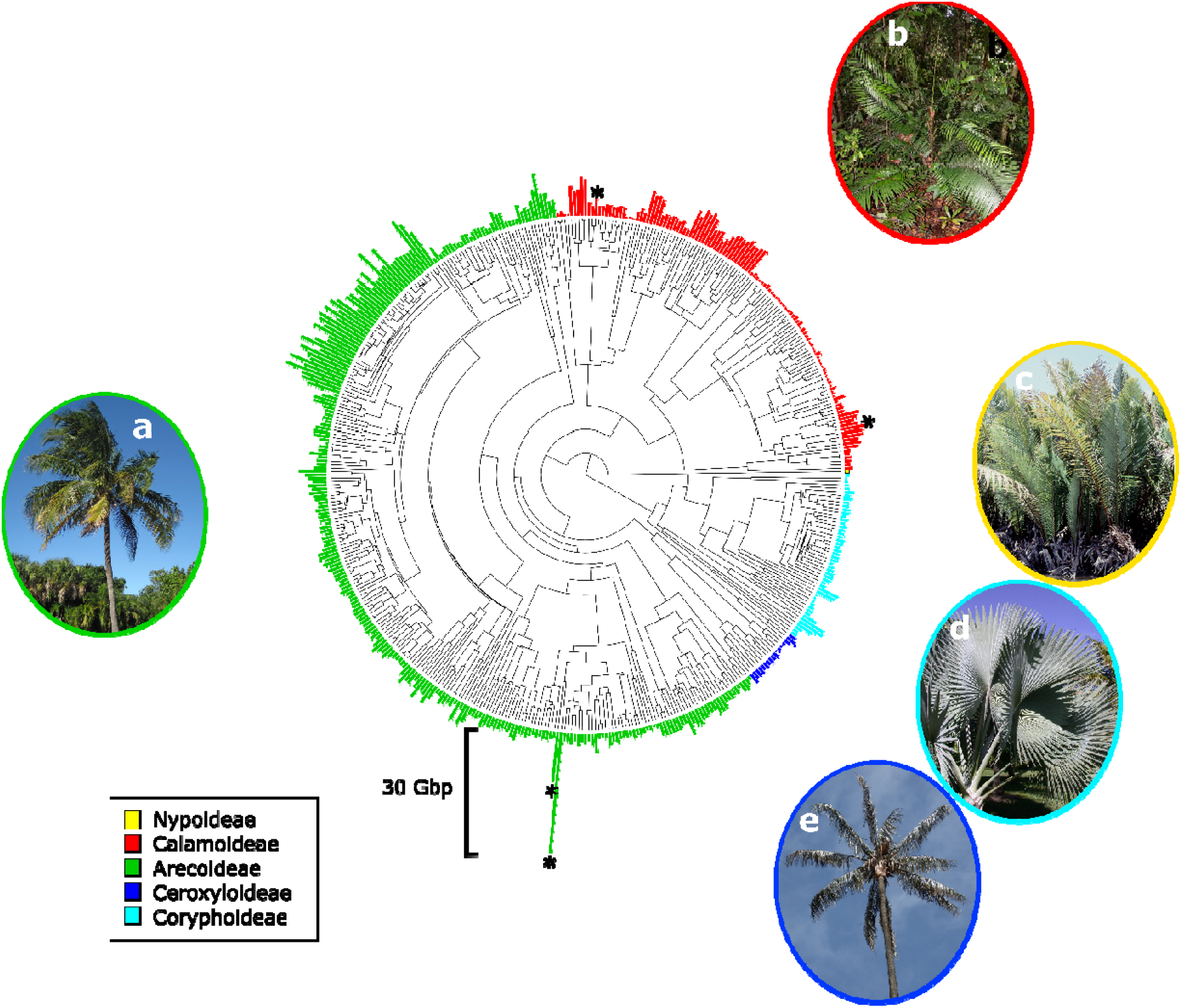
Phylogenetic tree of the Arecaceae (Faurby *et al*. 2016), with genome size data (1C values in gigabase pairs (Gbp)) for 472 species displayed as bars. Bars are coloured according to the palm subfamily to which each taxon belongs, and a 30 Gbp genome size bar is shown for scale. The four known polyploid palm species are indicated with asterisks (**\***). Photographs show palm species from each subfamily: a) *Cocos nucifera* (Arecoideae) *©* James St. John; b) *Calamus hirsutus* subsp. *korthalsii* (Calamoideae) *©* William J. Baker; c) *Nypa fruticans* (Nypoideae) *©* William J. Baker; d) *Bismarkia nobilis* (Coryphoideae) *©* William J. Baker; e) *Ceroxylon quindiuense* (Ceroxyloideae) *©* Alejandro Bayer.

We found significant evidence of phylogenetic signal in our genome size data (λ = 0.933, *P* = 1.749 × 10^−76^) and comparison of trait evolution models using AICc suggested that the Ornstein-Uhlenbeck model (i.e., evolution towards trait optima across the tree) was the best supported model (Δ*AICc* = 991.794 vs Brownian motion). A histogram showing the frequency distribution of genome sizes within the palm family is shown in Supporting Information 2: Fig. S3a.

### Aridity preferences of palm species help explain genome size variation

Modelling of the interaction between genome size and six *WorldClim* environmental variables using PGLS showed that a model containing ‘precipitation of the driest month’ and ‘minimum temperature of the coldest month’ with λ branch transformations best explained the observed variation in genome size (Δ*AICc* = 1.614). This model had a multiple *R*^2^ of 0.029 (*P=* 0.002, *d*.*f*.= 393), and is summarised in Table 1. The PGLS analysis excluding the four polyploid palm species recovered a similar minimum adequate model as above, with a multiple *R*^2^ of 0.030 (*P=* 0.002, *d*.*f*.= 389), and is summarised in Supporting Information 2: Table S4. Since precipitation of the driest month best explained genome size variation (Table 1) while minimum temperature of the coldest month was only marginally significant (*P*= 0.05), only precipitation of the driest month was used in further analyses. Precipitation of the driest month and genome size visualised onto the palm phylogenetic tree are shown in Supporting Information 2: Fig. S3b.

**Table 1:** Model summary for the minimum adequate PGLS model explaining variation in log(genome size) as a function of precipitation of the driest month and minimum temperature of the coldest month across the Arecaceae. Predictor variables (in this case, bioclimatic variables) which significantly explained variation in genome size (*P*<0.05) are indicated in bold.

|  | Estimate | Standard Error | *t* value | *Pr*(>\|*t*\|) |
| --- | --- | --- | --- | --- |
| ***Intercept*** | **0.806** | **0.345** | **2.339** | **0.020** |
| ***Precipitation of the driest month*** | **0.001** | **0.0003** | **3.385** | **7.849 x10<sup>-4</sup>** |
| *Min. temperature of the coldest month* | -0.001 | 0.0005 | -1.937 | 0.05 |

To see whether this relationship varied across the range of palm genome sizes, the genome size and precipitation data were divided into four quartiles (Table S2) and analysed using a Chi-squared test. This showed that genome size and aridity preference (precipitation of the driest month) were significantly more associated than would be expected by chance (χ2 = 37.889, P= 1.825×10-5, d.f. = 9; Fig. 3). Figure 3 also shows that species in the upper quartile of the genome size distribution were most likely to be found in very wet environments (Pearson’s residual = 3.97), and that those within the second quartile of the genome size distribution were far less likely to be found in very wet environments (Pearson’s residual = -2.71). In addition, species in this quartile were more likely to be found in semi-dry environments (Pearson residual = 2.01).

**Figure 3:**
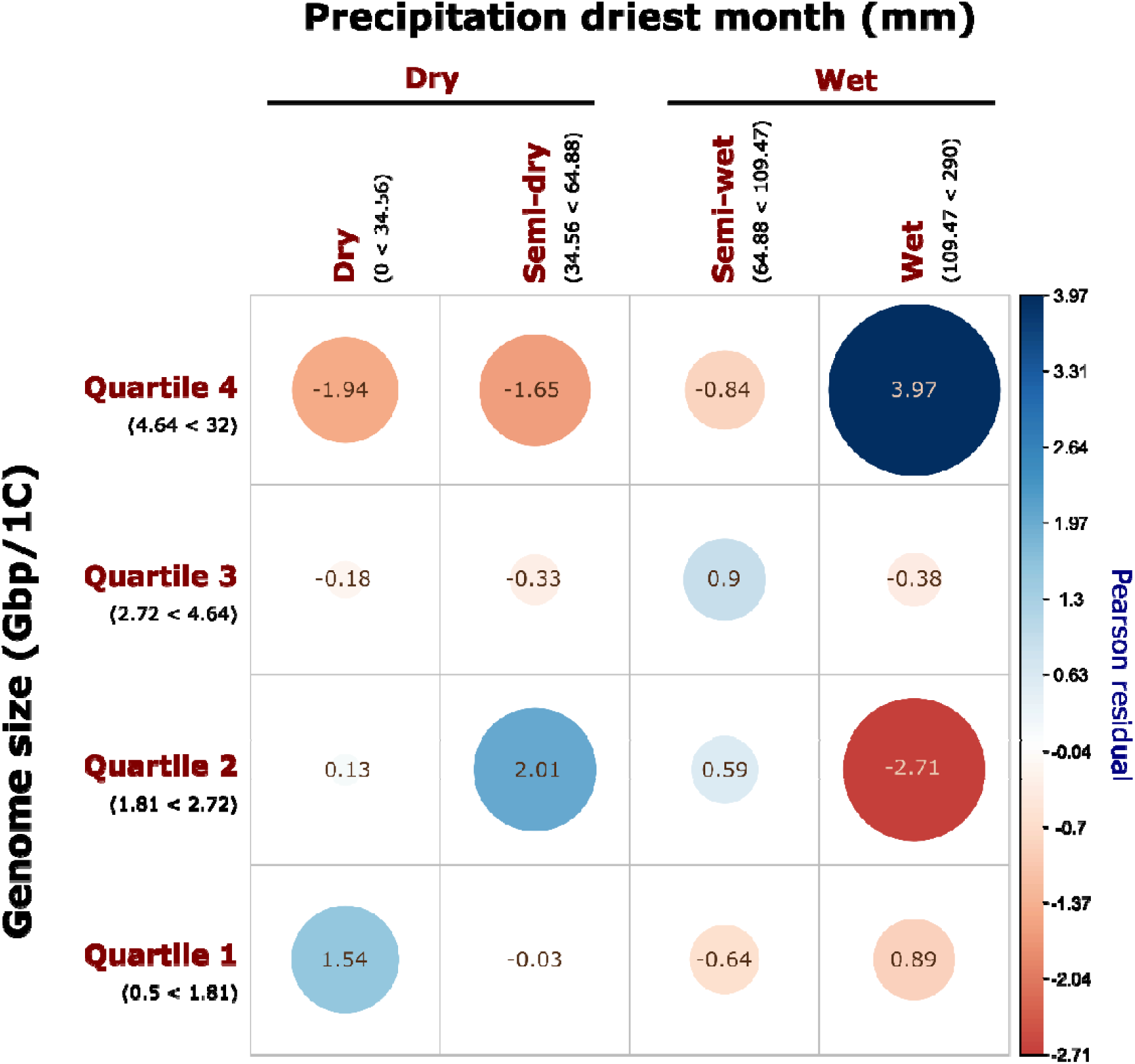
Association plot of genome size and precipitation of the driest month, binned into quartiles. Size of circles shows the size of the correlation coefficient (Pearson residuals) between variables for each relationship. The direction and magnitude of the correlation is indicated by the colour and colour intensity of each circle, with dark red denoting strongly negative associations and dark blue denoting strongly positive associations. Pearson residuals for each comparison are shown in the centre of each circle.

### Ecological metrics of palm repeat ‘communities’ vary with genome size

Calculations of the three ecological metrics (i.e. repeat richness (Menhinick’s index), total repeat occupancy and repeat diversity (Shannon index)) to characterise the repeat profiles of 141 palm species revealed considerable diversity across the palm phylogenetic tree (see Fig. S4 in Supporting Information 2). By exploring the relationships between these metrics with genome size, we found that repeat richness (Menhinick’s index), which reflects the richness of repeat lineages in a species’ genome, was explained best by total repeat occupancy (sometimes referred to as genome proportion, Novák *et al*. (2020a)) and genome size, as well as the interaction between them (Δ*AICc*=128.257, *R*^*2*^ = 0.699, *P*< 2.2×10^−16^ on 133 *d*.*f*.). Nevertheless, we observed that the relationship between repeat richness and genome size was not uniform across the diversity of genome sizes, as repeat richness decreased strongly as genome size increased, up to a threshold of around 5 Gbp/1C (Figure 4a) where it plateaued and thereafter increased with genome size. The summary for this model is given in Supporting Information 2, Table S5).

**Figure 4:**
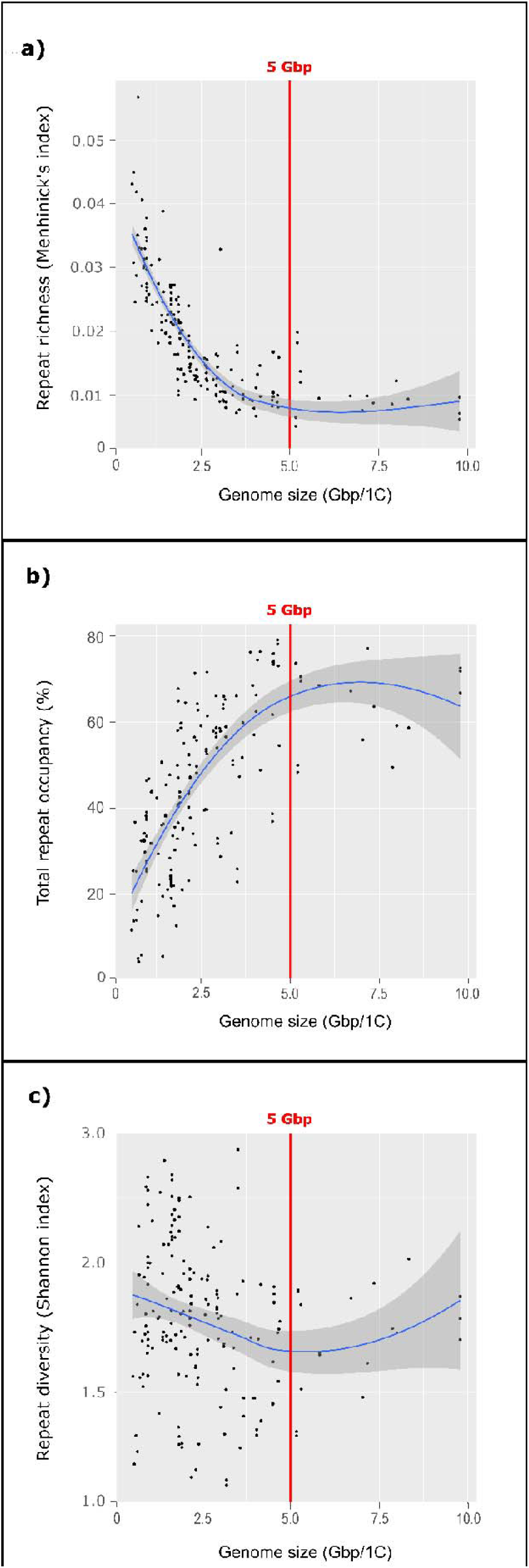
Scatterplots showing relationships between different ecological indices used to assess repeat profiles for each of the 141 palm species whose repeat compositions were analysed with *RepeatExplorer2*. Conditional means are shown by the blue line, calculated using Loess smoothing in *ggplot2*, and 95% confidence intervals are shown by the grey shading around the lines. The genome size at which repeat dynamics begin changing (5 gigabase pairs (Gbp)) are indicated by a red line on the *x-*axis. **a:** The relationship between genome size and repeat type richness (Menhinick’s Index). **b:** The relationship between genome size and total repeat occupancy of the genome. **c:** The relationship between genome size and repeat diversity (Shannon index).

Total repeat occupancy, i.e. the percentage of the genome occupied by repeats, was itself explained by genome size (Δ*AICc*=1.382, R^2^ = 0.167, *P*= 4.261×10-7 on 135 d.f.) but as for repeat richness, the direction of the relationships also changed depending on genome size (Fig. 4b). Thus, while total repeat occupancy increased with genome size up to a threshold of around 5 Gbp (Figure 4b), thereafter the relationship slightly decreased with further increases in genome size, thus repeats did not make up more than 80% of any genome, regardless of genome size. Neither repeat richness nor total repeat occupancy were explained by aridity preference (precipitation of the driest month).

In contrast to the above two ecological metrics, repeat diversity (Shannon index), which reflects the evenness in abundance of different repeat types within a genome, was not significantly explained by genome size, aridity preference or their interaction (Figure 4c). Thus while a weak negative correlation with genome size in smaller genomes (<5 Gbp) and potentially increasing diversity at larger genome sizes was observed in Figure 4c, linear modelling showed that neither were significant (data not shown).

### Repeat abundances correlate with genome size

Our phylogenetically corrected modelling of repeat profiles uncovered a significant signal of repeat expansion explaining genome size variation. PGLS modelling of genome size among species as a function of repeat lineage abundances in the genome showed that the amount of the genome occupied by the *Ty1-copia, Ty3-gypsy* and *TIR* superfamilies explained 57% of the genome size variation within palms using PGLS (Δ*AICc* = 0.965, multiple R^2^ = 0.57, *P*= <0.0001 on 126 *d*.*f*.). The *Ty1-copia* elements *Angela* and *TAR*, the *Ty3-gypsy* elements *CRM, Tekay* and *Retand*, and the *TIR* elements *EnSpm CACTA* and *MuDR Mutator* were all shown to be positively correlated with genome size, while pararetrovirus sequences were negatively correlated. The best fit model is summarised in Table 2.

**Table 2:** Model summary for the minimum adequate PGLS model explaining variation in log(Genome size) across palms according to the amounts of genome occupied by different repeat lineages. Predictor variables (I.e. the amount of the genome (Gbp) occupied by a repeat lineage) which significantly explained variation in genome size (*P*<0.05) are indicated in bold. Repeat superfamilies are indicated in the leftmost column, while the repeat lineages contained within them are shown in the column to their right.

|  |  | Estimate | Std. Error | t-value | Pr(>\|t\|) |
| --- | --- | --- | --- | --- | --- |
| *Intercept* |  | 0.323 | 0.214 | 1.510 | 0.134 |
| *Ty1-copia* | *Alesia* | -840.445 | 445.143 | -1.888 | 0.061 |
|  | *Angela* | **3.724** | **0.861** | **4.323** | **3.096 x10<sup>-5</sup>** |
|  | *TAR* | **122.195** | **43.005** | **2.841** | **0.005** |
| *Ty3-gypsy* | *CRM* | **6.710** | **2.498** | **2.687** | **0.008** |
|  | *Tekay* | **4.370** | **1.579** | **2.767** | **0.007** |
|  | *Retand* | **2.096** | **0.976** | **2.148** | **0.034** |
| *Pararetrovirus* |  | **-276.299** | **110.384** | **-2.503** | **0.014** |
| *TIR* | *EnSpm CACTA* | **20.413** | **8.323** | **2.453** | **0.016** |
|  | *hAT* | -126.147 | 72.394 | -1.743 | 0.084 |
|  | *MuDR Mutator* | **120.793** | **43.418** | **2.782** | **0.006** |

### Aridity preferences of palm species explain abundances of certain repeat lineages

PGLS modelling revealed that the absolute amounts (in Gbp) of *Ty3-gypsy, Ty1-copia* and *TIR* superfamilies, rDNA and satellite elements in the genome explained 18% of the variation in aridity preference among palm species (Δ*AICc* = 1.245, multiple R^2^ = 0.182, *P*= 0.002 on 127 d.f.). The abundance of the *TIR EnSpm CACTA* elements was positively correlated with precipitation of the driest month, suggesting that these elements are more abundant in palm species from wetter environments. In contrast, we observed a negative interaction between the amount of *Ty1-copia Angela* elements and aridity preference among palm species with small genomes (< 2.15 Gbp/1C), suggesting that *Angela* elements are more common small genomed species from drier environments. The best fit model is summarised in Table 3, which identifies both repeat lineages that are significantly correlated with aridity preference and other elements with non-significant slopes, but which still explained some variation in aridity preference. The amount of each species’ genome occupied by all repeat lineages is shown in Supporting Information 2: Fig. S5.

**Table 3:** Model summary for the minimum adequate PGLS model explaining variation in sqrt(Precipitation of the driest month) across palms according to the amounts of the genome occupied by different repeat lineages in Gbp, using genome size as a covariate. Predictor variables (in this case, amount of the genome occupied by a repeat lineage) which significantly explained variation in precipitation of the driest month (*P*<0.05) are indicated in bold. Repeat superfamilies are indicated in the leftmost column, and the repeat lineages contained within them are shown in the column to their right.

|  |  | Estimate | Std. Error | t-value | Pr(>\|t\|) |
| --- | --- | --- | --- | --- | --- |
| **Intercept** |  | **6.347** | **1.137** | **5.585** | **1.36x10<sup>-7</sup>** |
| *Ty1-copia* | *Ivana* | -994.575 | 573.787 | -1.733 | 0.085 |
|  | *Tork* | -55.490 | 29.249 | -1.897 | 0.060 |
| *Ty3-gypsy* | *Athila* | 51.177 | 35.181 | 1.455 | 0.148 |
| ***TIR*** | ***EnSpm CACTA*** | **129.360** | **64.418** | **2.008** | **0.047** |
| *rDNA* | *25S rDNA* | -982.724 | 504.525 | -1.948 | 0.054 |
|  | *5S rDNA* | 374.314 | 204.436 | 1.831 | 0.069 |
| *Satellite* |  | -80.825 | 55.162 | -1.465 | 0.145 |
| *Ty1-copia:Genome size class* | *Angela:Large genome* | -1.311 | 6.949 | -0.189 | 0.851 |
|  | ***Angela:Small genome*** | **-53.373** | **21.134** | **-2.526** | **0.013** |

## Discussion

Our study uncovered a distinct signal of repeat ‘communities’ influencing genome size and being structured by aridity. We found evidence of preferential expansion of different repeat lineages driving genome size variation, as well as associations between the abundance of two repeat lineages (*Ty1-copia Angela* and *TIR EnSpm CACTA*) and aridity preferences of palm species. Our work greatly expands existing genome size datasets for palms and is among the most extensive studies examining the ecological dynamics of repeats.

### Palm genome size variation

We found a 58-fold range of variation in genome size across the palm family, which agrees with the previously reported range from fewer species (Plant DNA C-values database, https://cvalues.science.kew.org/ (Pellicer & Leitch 2020)). The upper limit of genome size was found in the monotypic Madagascan endemic *Voanioala gerardii* (Figure 2, 30.63 Gbp/1C, 2n = c, 596 (Johnson *et al*., 1989; see also Röser, 1994)), which is 38-ploid and has the highest known chromosome number of all monocots. Excluding polyploid species, the largest genome belonged to *Pinanga sessilifolia* (15.40 Gbp/1C), and the genome sizes of the 46 *Pinanga* species analysed were among the most variable of all palm genera in this study, with the smallest being 6.55 Gbp/1C in *P. celebica*. This variability occurs despite the consistent chromosome numbers of 2n=32 reported for *Pinanga* species. Röser (1994; 1999) noted that rainforest understorey palm genera (e.g. *Chamaedorea, Geonoma, Licuala* and *Pinanga*) exhibited extreme variation in karyological traits, including genome size, and that species with very small and very large genomes were both able to exist in these wet environments. The smallest genomes we analysed belonged to another of these understorey genera, with genome sizes of 0.53 Gbp/1C in the diploids *Licuala orbicularis* and *Licuala sarawakensis*. Dransfield *et al*. (2008) stated that this variation in genome size is likely to be caused by the activity of repetitive DNA, suggesting that polyploidy plays a minimal role in palm genome size evolution (Barrett *et al*., 2019). This is supported by our study, as excluding the four polyploid species from our PGLS analyses did not materially impact minimum adequate models explaining genome size variation across Arecaceae (Supporting Information 2: Table S4).

### Aridity thresholds best explain palm genome size diversity

We found that genome size variation across palms showed significant phylogenetic signal (λ = 0.933, P = 1.749 × 10^−76^) and that genome size change across the tree better fit a model describing evolution towards genome size optima rather than random drift (Δ*AICc* = 991.794 between Ornstein-Uhlenbeck model vs. Brownian motion). This contrasts with previous work, which was based on more limited sampling (Barrett *et al*., 2019). Building on this, we found that, amongst bioclimatic variables, genome size variation was mostly explained by the aridity preferences of palm species, specifically by precipitation of the driest month (*P=*0.001, R^2^=0.03; Table 1). In addition, the relationship between aridity preference and genome size was not linear (Figure 3): while palm species from wet environments had large and small genomes, species from dry environments mostly had small genomes.

These results indicate that there is non-random evolution of genome size across the palms, and that yearly extremes of aridity may exert selective pressures on the upper limits of genome size. Genome size may have impacted adaptive evolution of plants through selection on minimum cell size, likely through simple scaling relationships between the two (i.e. species with larger genomes have larger minimum cell sizes (Faizullah *et al*., 2021)). Changes in cell size may then influence cell area/volume relationships, water mobility and biochemical reactions (e.g. Cavalier-Smith, 2005; Beaulieu *et al*., 2008). These in turn can influence photosynthesis (e.g. Roddy *et al*., 2020), gas exchange (e.g. Franks & Beerling 2009) and water use efficiency (e.g. Lawson & Blatt 2014) through their impacts on stomatal guard cell size and density (Veselý *et al*., 2012; Trávníček *et al*., 2019), all of which could exert ecological selection on a species’ genome size. Whilst an arrangement of large stomata at low density can prevent water loss, it also increases diffusion paths for CO2 and can reduce growth rates (Faizullah *et al*., 2021). This strategy has mainly been adapted by geophytes in arid areas (Veselý *et al*., 2012). In contrast, the evolution of small, high-density stomata may be favoured to enable faster growth (Franks & Beerling 2009) for non-geophytic taxa, such as palms. With many small stomata, costs in water loss can be ameliorated by the faster response rates of smaller stomata to rapid fluctuations in environmental conditions (Drake *et al*., 2013; Roddy *et al*., 2020). This may explain why we found that most arid-zone palm species had smaller genomes. Furthermore, selection on stomatal size in species with high stomatal density may become more relaxed with increasing water availability, potentially explaining why palms from wetter environments include species with the largest genomes.

### The ‘community ecology’ of repeats correlates with genome size

We found that the amount of the genome occupied by certain repeat lineages correlated significantly with genome size variation in palms. This suggests that the preferential expansion of particular repeat lineages drives genome size diversity in palms, as shown in other plant groups (e.g. Macas *et al*., 2015; Pellicer *et al*., 2021). Moreover, our results suggest that repeat richness and total repeat occupancy change asymptotically with genome size, indicating shifts in repeat ‘community’ composition and turnover across the range of palm genome sizes.

The abundance of repeats in a genome is maintained by the balance between expansion, epigenetic silencing and excision of elements through (retro)transposition, recombination and DNA repair (Schubert & Vu 2016; Wang *et al*., 2021). As such, any genome size gains made by transposition are eventually eroded by excision or mutational erosion (Petrov, 2002; Bennetzen & Wang 2014; Kelly *et al*., 2015). Accordingly, since repeats rarely provide an immediate selective advantage to their hosts, they often reach fixation within a genome largely through drift (Lynch & Walsh 2007). Therefore, those repeats present in higher proportions are more likely to be replicated and continue their expansion, as dictated by neutral models of community assemblage (Hubbell, 2001). This could explain the asymptotic relationships that we recovered between genome size and repeat richness and total occupancy (Figure 4; Supporting Information 2: Table S5), each of which exhibited changes in their rate of increase or decrease at a genome size of around 5 Gbp/1C. Amongst genomes <5Gbp/1C, the lower repeat richness associated with increasing genome size is likely due to the stochastic expansion of some repeat lineages but not others, as shown in other organisms (e.g. Serra *et al*., 2013). This results in the domination of repeat communities by a few repeat lineages (Venner *et al*., 2009), as the most abundant repeats are themselves more likely to amplify. This dynamic changes at around 5 Gbp/1C, where the tendency for domination of the genome by a few repeat lineages lessens as more repeats rise in copy number. Novák *et al*. (2020a) argued that this change in repeat dynamics is best explained by the gradual mutation and ‘fossilisation’ of older repeats driven by lower turnover, resulting in gradual accumulation of sequences which inactivate and mutate to the point that they cease to resemble their repeat progenitors. It is perhaps notable that the threshold of *c*. 5 Gbp/1C differs from that reported by Novák *et al*. (2020a) of *c*. 10 Gbp/1C, whose analysis was based on 101 diploid species across the diversity of angiosperms and gymnosperms, encompassing a 1,475-fold range in genome size. Further analyses focused at the family level are clearly needed to uncover to what extent dynamics in repeat turnover differ between families.

PGLS modelling indicated that the amount of the genome occupied by several repeat lineages from the *Ty1-copia, Ty3-gypsy* and *TIR* superfamilies explained 57% of the genome size variation within palms (Table 2). Specifically, species with larger genomes had higher amounts of *Angela* and *TAR* elements (*Ty1-copia* superfamily), *CRM, Tekay* and *Retand* elements (*Ty3-gypsy* superfamily), and *EnSpm CACTA* and *MuDR Mutator* elements (*TIR* superfamily) (Table 2, Supporting Information 2: Fig. S5). As such, it appears that stochastic expansion of these repeat lineages occurs in some palm species but not in others, driving genome size change (as shown in Serra *et al*., 2013). However, our analyses of genome size (Table 1; Figure 3) also indicate that there may be an advantage for species with compact genomes when under drought stress (e.g. Ibarra-Laclette *et al*., 2013; Kelley *et al*., 2014). This suggests that extrinsic processes may govern repeat community composition and give rise to genome size diversity in palms.

### Repeat dynamics may be modulated by aridity

While stochastic expansion of certain repeats may explain much genome size diversity in palms, it cannot fully explain the expansion of *EnSpm CACTA* repeats in species from wetter environments or the decrease in *Ty1-copia Angela* elements in small-genomed, xerophilous palm species that we observed (Table 3). Neutral processes responsible for the structure of repeat ‘communities’ are subject to extrinsic modulators, just as there are extrinsic modulators of community structure in ecosystems (reviewed in Dunson & Travis 1991). As such, our analyses suggest that arid environments select against larger genomes, such that repeat amplification is greatly reduced above an environmentally-constrained genome size optimum (i.e., the ‘carrying capacity’ (*K*) of the genome (Brookfield, 2005)). This indicates that the expansion of most repeat families in palms is selected against in species from harsher, drier environments, as suggested as a strategy for salinity tolerance in mangrove species (Lyu *et al*., 2018). In palm species from wetter environments, it is therefore possible that selection against genome size enlargement is relaxed, and so repeats such as *EnSpm CACTA* elements are free to amplify.

However, in direct contrast to this broader pattern, we found that *Ty1-copia Angela* elements were more abundant in xerophilous palm species with smaller genomes (Table 3). It is known that the expansion of repeats may be stress-induced (Casacuberta & González 2013; Galindo-González *et al*., 2017) and that this expansion is associated with epigenetic changes, sometimes leading to a reshuffling of the genome (reviewed in Bennetzen & Wang 2014). Indeed, long terminal repeat (LTR) retrotransposons, which includes *Ty1-copia Angela* elements, are particularly prone to expansion under stressful conditions (Galindo-González *et al*., 2017). When under abiotic stress, these LTR elements may bypass the regulatory machinery of the cell because they carry cis-regulatory regions in the 5’ LTR sequence controlling transcription. These regions tend to be shared with stress-response genes in the host, and can allow LTR elements to avoid epigenetic silencing when stress-response genes are activated (as in Cavrak *et al*., 2014; Galindo-González *et al*., 2017). This may explain the expansion of *Angela* elements in small-genomed, xerophilous palm species that we observed. Similar patterns of repeat expansion mediated by water stress have been shown for *BARE-1*, another LTR element which is associated with water stress-induced genes, in wild barley (*Hordeum spontaneum*) (Kalendar *et al*., 2000). Such associations may help to explain why LTR elements are the most abundant group of repeats in plant genomes (Bennetzen & Wang 2014), and why members of the *Ty1-copia* superfamily have evolved a tendency to insert near genes, given the adaptive benefit of evading cellular surveillance and excision (White *et al*., 1994; Lockton & Gaut 2009; Galindo-González & Deyholos 2012). Indeed, remnants of LTRs within many stress-response genes are necessary for their functioning, possibly alluding to the past adaptive co-option of LTRs by plant genomes (Jangam *et al*., 2017).

### Conclusions

Overall, we show that genome size within the palm family is influenced by the expansion of repeats, and that the dynamics of these repeat ‘communities’ are moderated by aridity through the selective pressure aridity exerts on repeat amplification and genome size. Our results show that while repeat ‘communities’ may be assembled largely by stochastic processes governing expansion at the level of the individual element, repeat expansion is constrained under arid climatic regimes. In contrast, we also show that certain retrotransposon lineages (e.g., *Ty1-copia Angela* elements) have amplified in arid environments, possibly through their association with stress-response genes. This suggests that interactions between repeat communities, the abiotic environment and genome size influence the ecology of palm genomes.

## Supporting Information

Additional supporting information may be found in the online version of this article.

Supporting Information 1: Table of newly generated genome size data for 437 palm species.

Supporting Information 2:

**Fig. S1:** Phylogenetic spread of genome size data for 472 palm species collected during this study and used for phylogenetic generalised least squares (P.G.L.S) modelling.

**Fig. S2:** Phylogenetic spread of genome skimming data for 141 palm species used to estimate repeat profiles with *RepeatExplorer2*.

**Fig. S3a:** Histogram showing the distribution of genome size for 472 species across the palm family.

**Fig. S3b:** *CoPhylo* plot produced using *phytools* showing genome size and aridity preference (precipitation of the driest month) variation across the palm family.

**Fig. S4:** Genome size, repeat richness (Menhinick’s Index), percentage of the genome occupied by repeats and repeat diversity (Shannon Index) for 141 palm species superimposed on the Faurby et al. (2016) phylogenetic tree.

**Fig. S5:** The amount of the genome occupied for all repeat lineages analysed, shown for the subset of palm species for which genome skimming data were available.

**Table S1:** Accessions and voucher information for palms sampled in the genome skimming dataset.

**Table S2:** Genome size (a) and precipitation of the driest month (b) bins used for chi-squared analysis

**Table S3:** Hierarchical groupings of repeat lineages as defined by the RExDB database.

**Table S4:** Model summary for minimum adequate PGLS model explaining variation in log(Genome size) across the Arecaceae, excluding the four polyploid palm species.

**Table S5:** Model summary for minimum adequate PGLS model explaining variation in repeat type richness (log(Mehinick’s Index)) as a function of genome size and total genome occupancy of repeats across the Arecaceae.

**Methods S1:** Details of genome size measurement and calculation of repeat type richness, occupancy and diversity

Supporting information 3: Scripts necessary for conducting and analysing the output of

RepeatExplorer2 analyses.

## Acknowledgements

The authors acknowledge the Research/Scientific Computing teams at The James Hutton Institute and NIAB for providing computational resources and technical support for the “UK’s Crop Diversity Bioinformatics HPC” (BBSRC grant BB/S019669/1), use of which has contributed to the results reported within this paper. Computational resources were also in part provided by the project “e-Infrastruktura CZ” (e-INFRA CZ ID:90140) supported by the Ministry of Education, Youth and Sports of the Czech Republic. The authors would like to thank the Royal Botanic Gardens, Kew (UK), Montgomery Botanical Center (US), Fairchild Tropical Botanic Garden (US), Prague Botanical Garden (CZ) and the Frankfurt Palmengarten (DE) for help with collecting samples. This study was partly financially supported by the Strategic Priority Research Program of the Chinese Academy of Sciences (No. XDB31030100) and the Biological Resources Program of Chinese Academy of Sciences (ZSSD-009). Xue-Jun Ge would like to thank the Biological Resources Program of the Chinese Academy of Sciences (ZSSD-009) and the International Partnership Program of the Chinese Academy of Science (grant number:151853KYSB20190027). JP was supported by a Ramón y Cajal Fellowship (RYC-2017-2274) funded by the Ministerio de Ciencia y Tecnología (Gobierno de España). SB was funded by a Garfield Weston Foundation postdoctoral fellowship. PN and JM were supported by the ELIXIR CZ Research Infrastructure Project (Czech Ministry of Education, Youth and Sports; grant no. LM2018131). We would also like to thank Joel Brown & Matt Lloyd Jones for ecological analysis advice, Yijing Lu for help compiling and cleaning palm occurrence data, and Robyn Powell for help with flow cytometry.

## Author Contribution

This study was conceived by R.J.S. Analyses were performed by R.J.S. and P.N., with guidance from J. P., S.B., M.S.G., S.D., J.M., A.L. and I.L. Genome size data were generated by J.P., J.S., D.F. and I.L. Genome skimming data were produced by X-J.G. and C.B. The manuscript was written by R.J.S. with contributions from J.P., X-J.G., C.B., S.B., M.S.G., P.N., D.F., W.J.B., S.D., J.M., A.R., and I.L.

## Data Availability

The data that support the findings of this study are openly available from online repositories. All raw reads generated using genome skimming which were used to assess palm repeat profiles are available on the NCBI Sequence Read Archive with the Accession nos SAMN21016546-SAMN21016686, under the BioProject number PRJNA758225. All GBIF distribution data and *WorldClim* climate data for each palm species are available on Dryad (DOI TBA). Data are under embargo until publication, and any further data required are available from the corresponding author upon reasonable request.

## Notes

### Competing Interest Statement

The authors have declared no competing interest.

