## Supporting Information 2 for "The Ecology of Palm Genomes: Repeat-associated genome size expansion is constrained by aridity"

**Supplementary figures**

| **Figure** | **Page** |
| --- | --- |
| Figure S1 | 2 |
| Figure S2 | 3 |
| Figure S3a | 4 |
| Figure S3b | 5 |
| Figure S4 | 7 |
| Figure S5 | 9 |

**Supplementary tables**

| **Table** | **Page** |
| --- | --- |
| Table S1 | 10 |
| Table S2a | 16 |
| Table S2b | 16 |
| Table S3 | 17 |
| Table S4 | 18 |
| Table S5 | 18 |

**Methods S1**

| **Table** | **Page** |
| --- | --- |
| Genome size measurement | 19 |
| Calculating repeat type richness, occupancy and diversity | 19 |

***Supplementary figures***

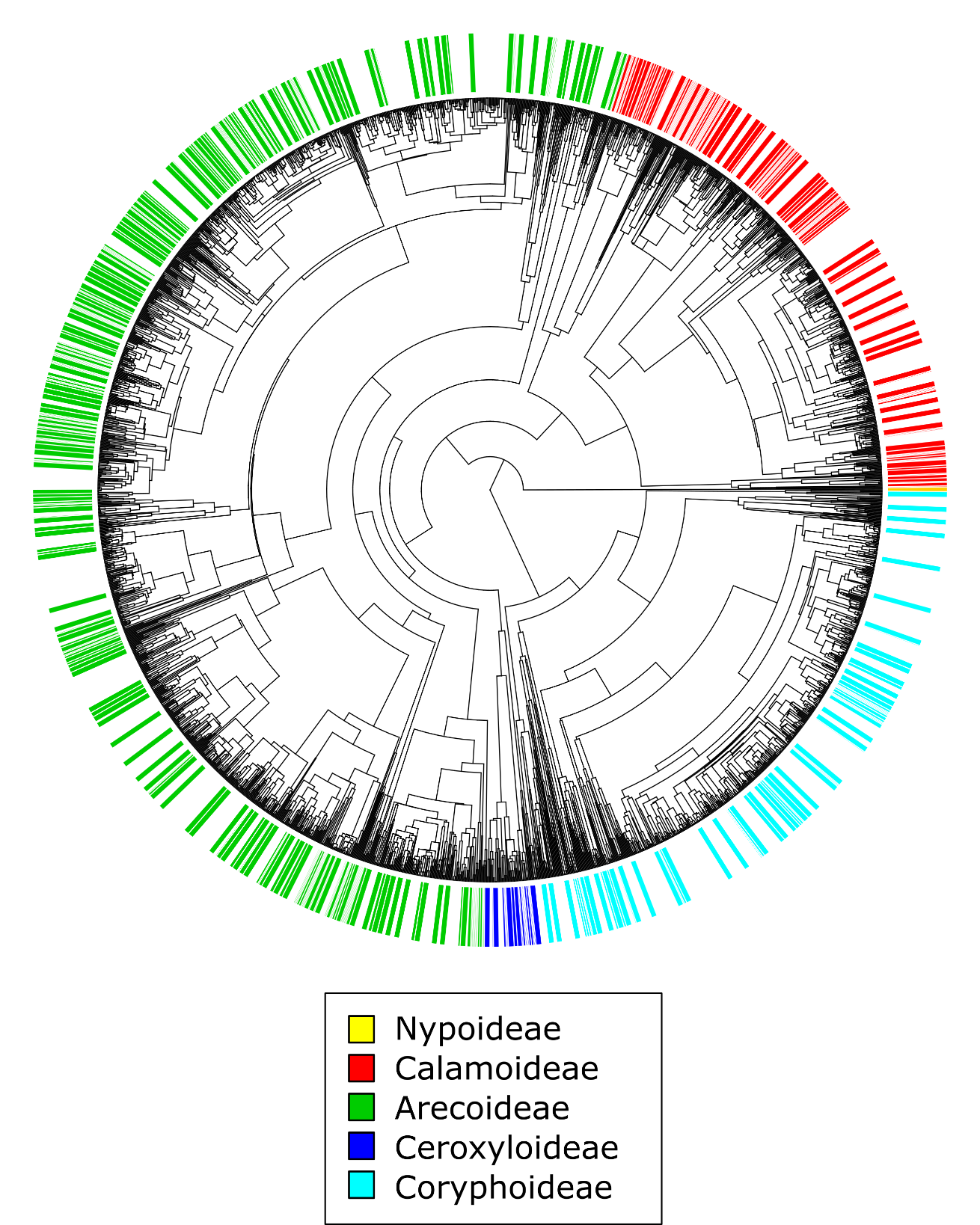

**Fig. S1:** Phylogenetic spread of genome size data for 472 palm species collected during this study and used for phylogenetic generalised least squares (P.G.L.S) modelling. This shows all five palm subfamilies were represented in our sampling.

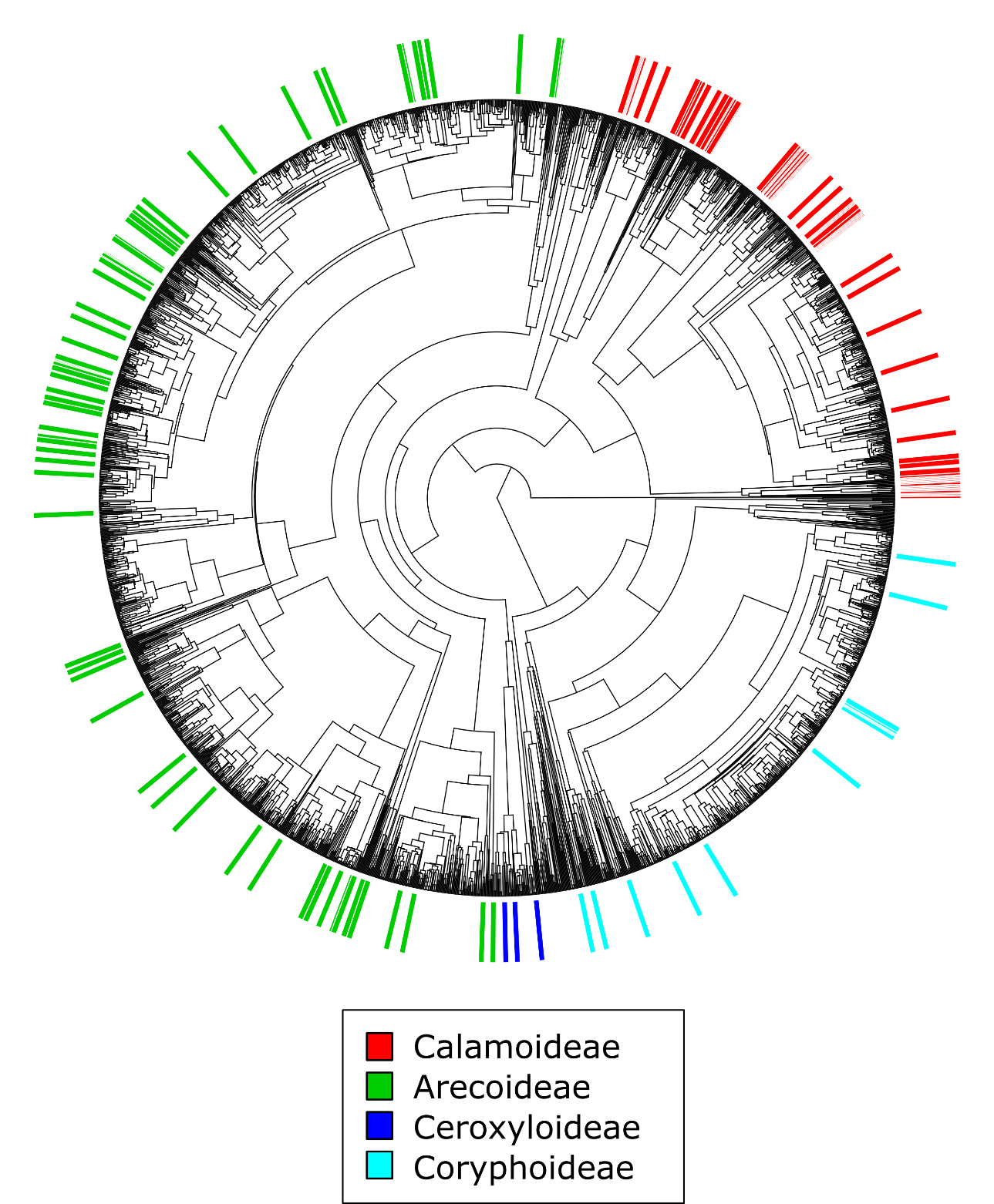

**Fig. S2:** Phylogenetic spread of genome skimming data for 141 palm species used to estimate repeat profiles with *RepeatExplorer2*, showing representational sampling of all palm subfamilies except the monospecific Nypoideae.

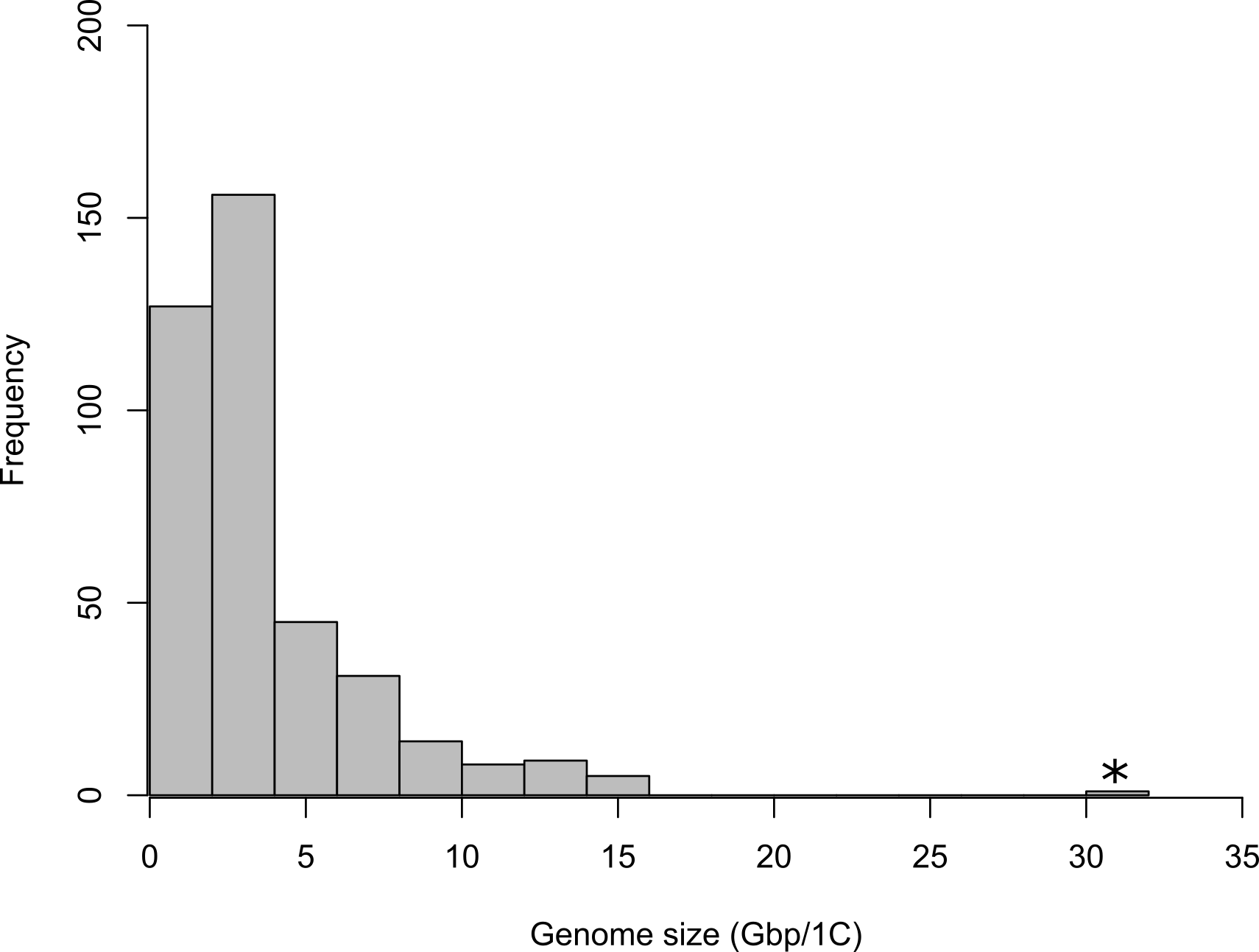

**Fig. S3a:** Histogram showing the distribution of genome size for 472 species across the palm family. The outlying value in the rightmost bin, representing the polyploid *Voanioala gerardii* (30.631 Gbp/1C), is shown with an asterisk (*).

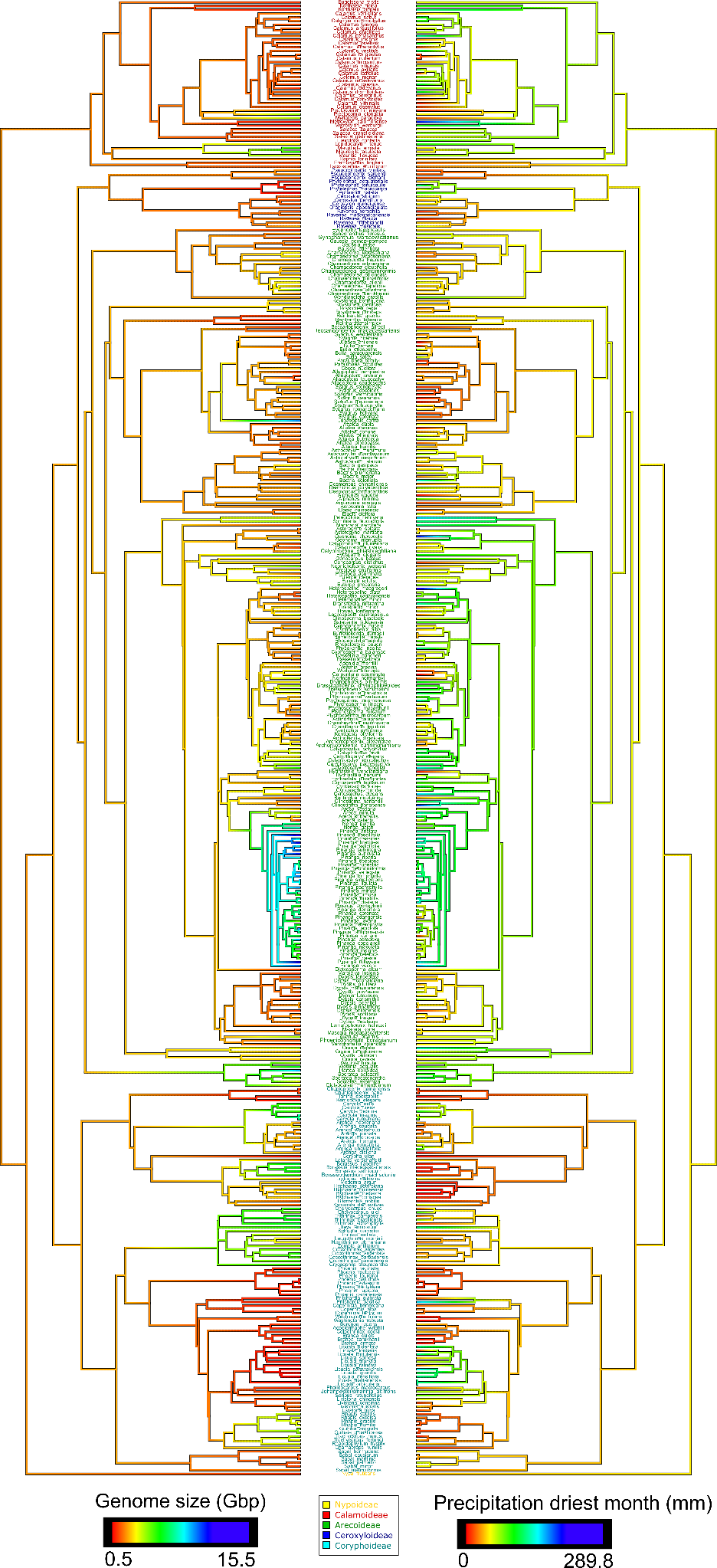

**Fig. S3b:** *CoPhylo* plot produced using *phytools* showing genome size and aridity preference (precipitation of the driest month) variation across the palm family, excluding the polyploid *Voanioala gerardii.* Species names are shown coloured by palm subfamily.

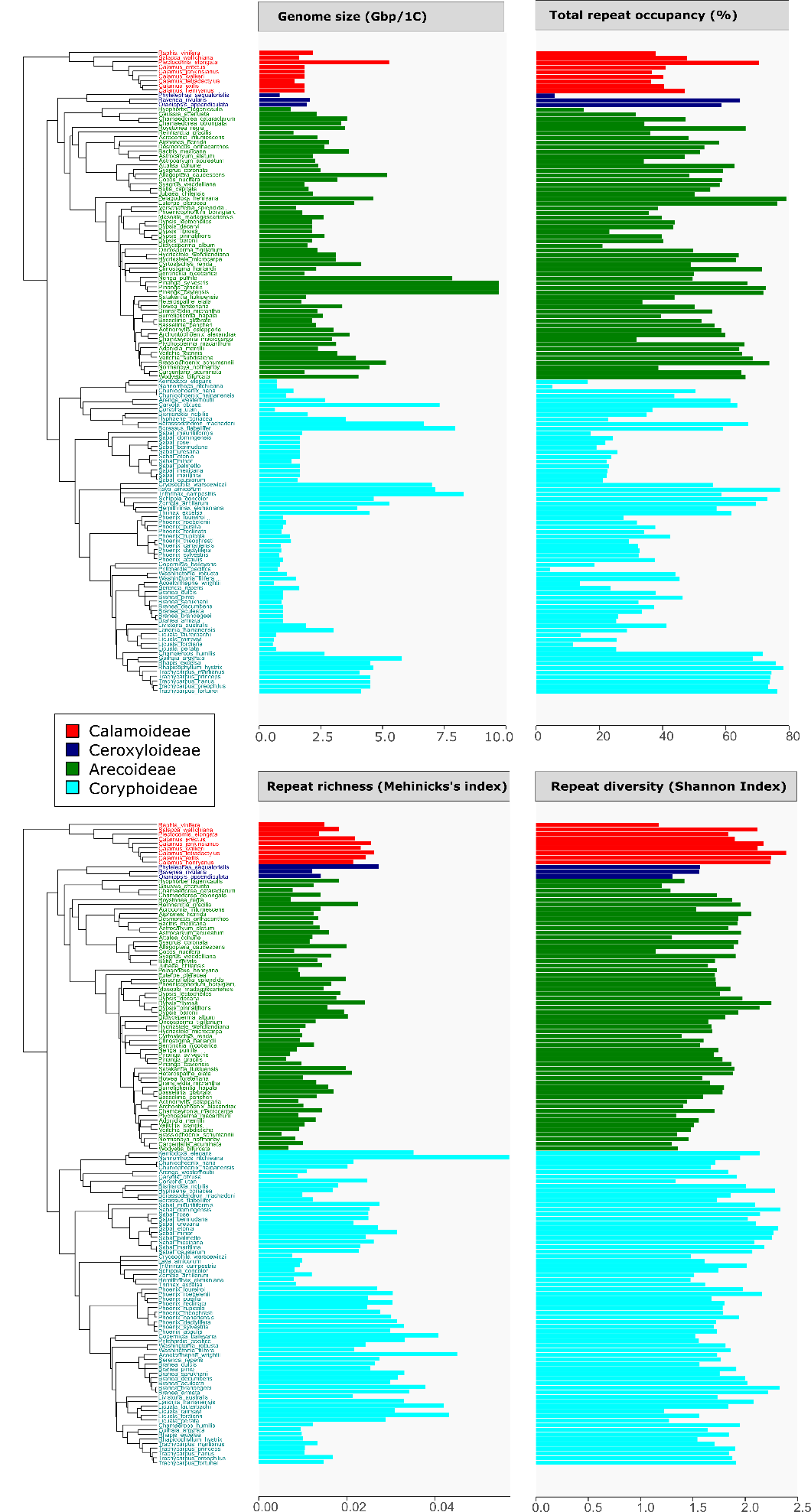

**Fig. S4:** Genome size, repeat richness (Menhinick’s Index), percentage of the genome occupied by repeats and repeat diversity (Shannon Index) for 141 palm species superimposed on the Faurby et al. (2016) phylogenetic tree. Species names are shown, coloured by palm subfamily using the same colours as in Fig. S1.

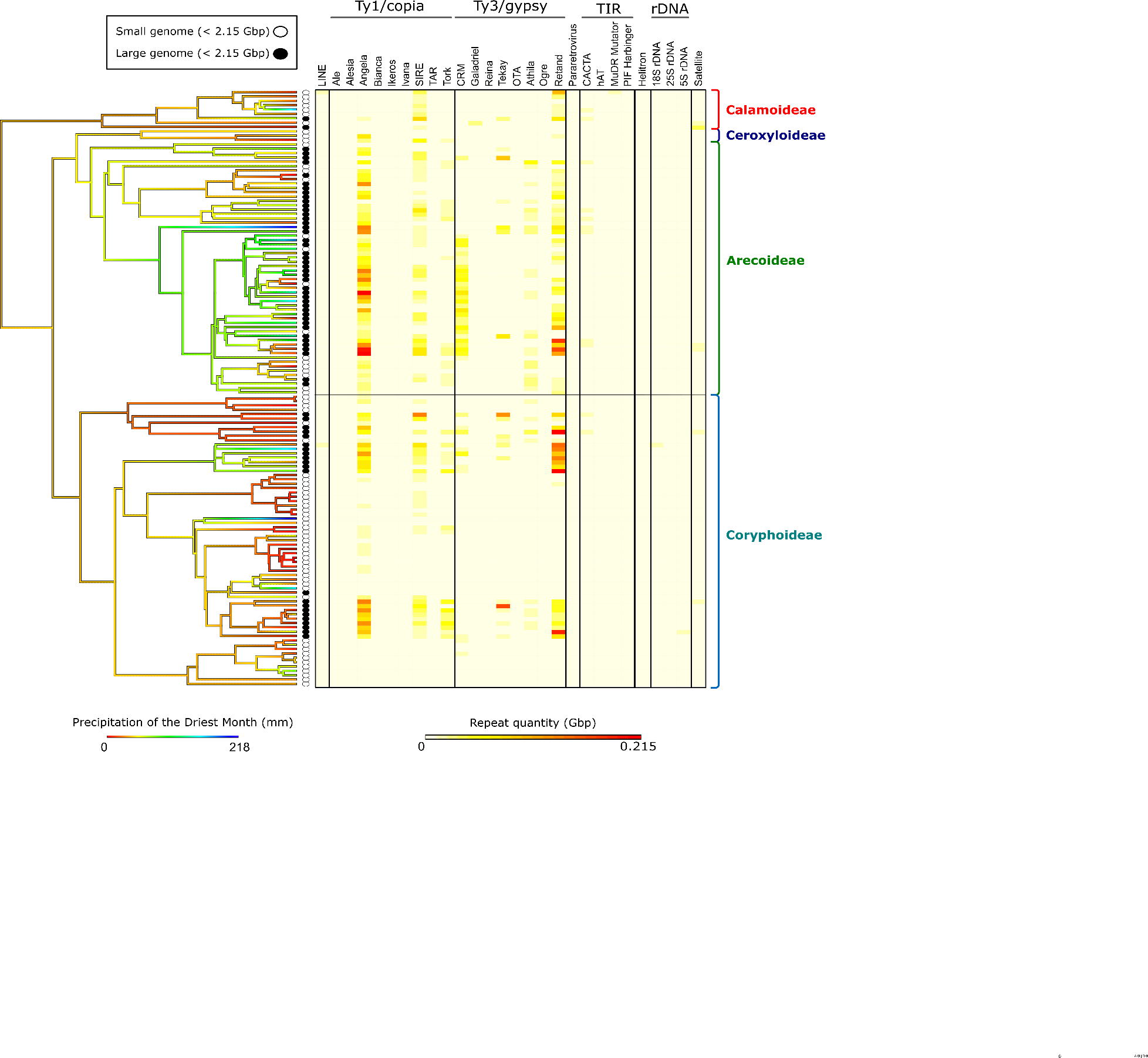

**Fig. S5:** The amount of the genome occupied for all repeat lineages analysed, shown for the subset of palm species for which genome skimming data were available. Repeat lineages are grouped by repeat superfamily above the heatmap, with corresponding columns. Precipitation of the driest month is shown reconstructed onto the Faurby *et al*. (2016) phylogenetic tree of the palm family. Genome sizes, binned into the same two groups based on the median value as in the PGLS analysis, are also shown on the tree, with black circles indicating species with large genomes (>2.15 Gbp) and with white circles showing species with small genome sizes (≤2.15 Gbp). Palm subfamilies are also labelled on the phylogeny, shown to the right of the heatmap.

***Supplementary tables***

| **Species** | **Collector n^o^** | **Collector** | **Location** |
| --- | --- | --- | --- |
| *Acoelorrhaphe wrightii* | SCBG014 | Duc Thanh Le, Yu-qu Zhang | South China Botanical Garden |
| *Acrocomia intumescens* | XMBG202 | Duc Thanh Le, Yu-qu Zhang | Xiamen Botanical Garden |
| *Actinorhytis calapparia* | MDBG01 | Craig Barrett | Fairchild Tropical Botanic Garden |
| *Adonidia merrillii* | XTBG042 | Duc Thanh Le, Yu-qu Zhang | Xishuangbanna Tropical Botanical Garden |
| *Aiphanes horrida* | XTBG235 | Duc Thanh Le, Yu-qu Zhang | Xishuangbanna Tropical Botanical Garden |
| *Allagoptera caudescens* | XTBG125 | Duc Thanh Le, Yu-qu Zhang | Xishuangbanna Tropical Botanical Garden |
| *Archontophoenix alexandrae* | SCBG887 | Duc Thanh Le, Yu-qu Zhang | South China Botanical Garden |
| *Arenga westerhoutii* | SCBG897 | Duc Thanh Le, Yu-qu Zhang | South China Botanical Garden |
| *Astrocaryum aculeatum* | XMBG043 | Duc Thanh Le, Yu-qu Zhang | Xiamen Botanical Garden |
| *Astrocaryum alatum* | SCBG021 | Duc Thanh Le, Yu-qu Zhang | South China Botanical Garden |
| *Attalea cohune* | SCBG946 | Duc Thanh Le, Yu-qu Zhang | South China Botanical Garden |
| *Bactris mexicana* | SCBG013 | Duc Thanh Le, Yu-qu Zhang | South China Botanical Garden |
| *Basselinia glabrata* | XMBG171 | Duc Thanh Le, Yu-qu Zhang | Xiamen Botanical Garden |
| *Basselinia pancheri* | XMBG142 | Duc Thanh Le, Yu-qu Zhang | Xiamen Botanical Garden |
| *Bentinckia nicobarica* | XTBG227 | Duc Thanh Le, Yu-qu Zhang | Xishuangbanna Tropical Botanical Garden |
| *Bismarckia nobilis* | SCBG894 | Duc Thanh Le, Yu-qu Zhang | South China Botanical Garden |
| *Borassodendron machadonis* | XTBG018 | Duc Thanh Le, Yu-qu Zhang | Xishuangbanna Tropical Botanical Garden |
| *Borassus flabellifer* | SCBG015 | Duc Thanh Le, Yu-qu Zhang | South China Botanical Garden |
| *Brahea aculeata* | XMBG210 | Duc Thanh Le, Yu-qu Zhang | Xiamen Botanical Garden |
| *Brahea brandegeei* | XMBG411 | Duc Thanh Le, Yu-qu Zhang | Xiamen Botanical Garden |
| *Brahea clara* | XMBG252 | Duc Thanh Le, Yu-qu Zhang | Xiamen Botanical Garden |
| *Brahea decumbens* | XMBG145 | Duc Thanh Le, Yu-qu Zhang | Xiamen Botanical Garden |
| *Brahea dulcis* | XMBG213 | Duc Thanh Le, Yu-qu Zhang | Xiamen Botanical Garden |
| *Brahea pimo* | XMBG146 | Duc Thanh Le, Yu-qu Zhang | Xiamen Botanical Garden |
| *Brahea sarukhanii* | XMBG197 | Duc Thanh Le, Yu-qu Zhang | Xiamen Botanical Garden |
| *Brassiophoenix schumannii* | XTBG118 | Duc Thanh Le, Yu-qu Zhang | Xishuangbanna Tropical Botanical Garden |
| *Burretiokentia hapala* | SCBG089 | Duc Thanh Le, Yu-qu Zhang | South China Botanical Garden |
| *Butia capitata* | SCBG875 | Duc Thanh Le, Yu-qu Zhang | South China Botanical Garden |
| *Calamus erectus* | XTBG070 | Duc Thanh Le, Yu-qu Zhang | Xishuangbanna Tropical Botanical Garden |
| *Calamus exilis* | XTBG132 | Duc Thanh Le, Yu-qu Zhang | Xishuangbanna Tropical Botanical Garden |
| *Calamus henryanus* | XTBG068 | Duc Thanh Le, Yu-qu Zhang | Xishuangbanna Tropical Botanical Garden |
| *Calamus jenkinsianus* | XTBG071 | Duc Thanh Le, Yu-qu Zhang | Xishuangbanna Tropical Botanical Garden |
| *Calamus tetradactylus* | XMBG017 | Duc Thanh Le, Yu-qu Zhang | Xiamen Botanical Garden |
| *Calamus walkeri* | XTBG134 | Duc Thanh Le, Yu-qu Zhang | Xishuangbanna Tropical Botanical Garden |
| *Carpentaria acuminata* | XMBG016 | Duc Thanh Le, Yu-qu Zhang | Xiamen Botanical Garden |
| *Caryota obtusa* | SCBG872 | Duc Thanh Le, Yu-qu Zhang | South China Botanical Garden |
| *Chamaedorea cataractarum* | SCBG019 | Duc Thanh Le, Yu-qu Zhang | South China Botanical Garden |
| *Chamaedorea oblongata* | XTBG117 | Duc Thanh Le, Yu-qu Zhang | Xishuangbanna Tropical Botanical Garden |
| *Chamaerops humilis* | XMBG060 | Duc Thanh Le, Yu-qu Zhang | Xiamen Botanical Garden |
| *Chambeyronia macrocarpa* | SCBG074 | Duc Thanh Le, Yu-qu Zhang | South China Botanical Garden |
| *Chuniophoenix hainanensis* | SCBG987 | Duc Thanh Le, Yu-qu Zhang | South China Botanical Garden |
| *Chuniophoenix nana* | SCBG963 | Duc Thanh Le, Yu-qu Zhang | South China Botanical Garden |
| *Clinostigma harlandii* | XMBG044 | Duc Thanh Le, Yu-qu Zhang | Xiamen Botanical Garden |
| *Cocos nucifera* | XMBG116 | Duc Thanh Le, Yu-qu Zhang | Xiamen Botanical Garden |
| *Copernicia baileyana* | XTBG122 | Duc Thanh Le, Yu-qu Zhang | Xishuangbanna Tropical Botanical Garden |
| *Corypha utan* | XMBG023 | Duc Thanh Le, Yu-qu Zhang | Xiamen Botanical Garden |
| *Cryosophila warscewiczii* | SCBG138 | Duc Thanh Le, Yu-qu Zhang | South China Botanical Garden |
| *Cyrtostachys renda* | XTBG100 | Duc Thanh Le, Yu-qu Zhang | Xishuangbanna Tropical Botanical Garden |
| *Desmoncus orthacanthos* | XTBG022 | Duc Thanh Le, Yu-qu Zhang | Xishuangbanna Tropical Botanical Garden |
| *Dictyosperma album* var. *aureum* | SCBG999 | Duc Thanh Le, Yu-qu Zhang | South China Botanical Garden |
| *Dransfieldia micrantha* | MDBG06 | Craig Barrett | Fairchild Tropical Botanic Garden |
| *Dypsis baronii* | SCBG139 | Duc Thanh Le, Yu-qu Zhang | South China Botanical Garden |
| *Dypsis decaryi* | SCBG888 | Duc Thanh Le, Yu-qu Zhang | South China Botanical Garden |
| *Dypsis fibrosa* | XTBG094 | Duc Thanh Le, Yu-qu Zhang | Xishuangbanna Tropical Botanical Garden |
| *Dypsis leptocheilos* | SCBG880 | Duc Thanh Le, Yu-qu Zhang | South China Botanical Garden |
| *Dypsis pinnatifrons* | XTBG225 | Duc Thanh Le, Yu-qu Zhang | Xishuangbanna Tropical Botanical Garden |
| *Euterpe oleracea* | XTBG230 | Duc Thanh Le, Yu-qu Zhang | Xishuangbanna Tropical Botanical Garden |
| *Gaussia attenuata* | XTBG226 | Duc Thanh Le, Yu-qu Zhang | Xishuangbanna Tropical Botanical Garden |
| *Guihaia argyrata* | SCBG105 | Duc Thanh Le, Yu-qu Zhang | South China Botanical Garden |
| *Hemithrinax ekmaniana* | MDBG08 | Craig Barrett | Fairchild Tropical Botanic Garden |
| *Heterospathe elata* | XTBG218 | Duc Thanh Le, Yu-qu Zhang | Xishuangbanna Tropical Botanical Garden |
| *Howea forsteriana* | SCBG040 | Duc Thanh Le, Yu-qu Zhang | South China Botanical Garden |
| *Hydriastele microcarpa* | XTBG047 | Duc Thanh Le, Yu-qu Zhang | Xishuangbanna Tropical Botanical Garden |
| *Hydriastele wendlandiana* | XTBG054 | Duc Thanh Le, Yu-qu Zhang | Xishuangbanna Tropical Botanical Garden |
| *Hyophorbe lagenicaulis* | XMBG224 | Duc Thanh Le, Yu-qu Zhang | Xiamen Botanical Garden |
| *Hyphaene coriacea* | XTBG120 | Duc Thanh Le, Yu-qu Zhang | Xishuangbanna Tropical Botanical Garden |
| *Itaya amicorum* | MDBG12 | Craig Barrett | Fairchild Tropical Botanic Garden |
| *Jubaea chilensis* | XMBG203 | Duc Thanh Le, Yu-qu Zhang | Xiamen Botanical Garden |
| *Kerriodoxa elegans* | SCBG081 | Duc Thanh Le, Yu-qu Zhang | South China Botanical Garden |
| *Lanonia hainanensis* | SCBG877 | Duc Thanh Le, Yu-qu Zhang | South China Botanical Garden |
| *Licuala fordiana* | XMBG216 | Duc Thanh Le, Yu-qu Zhang | Xiamen Botanical Garden |
| *Licuala lauterbachii* | XTBG104 | Duc Thanh Le, Yu-qu Zhang | Xishuangbanna Tropical Botanical Garden |
| *Licuala peltata* | SCBG912 | Duc Thanh Le, Yu-qu Zhang | South China Botanical Garden |
| *Licuala ramsayi* | SCBG008 | Duc Thanh Le, Yu-qu Zhang | South China Botanical Garden |
| *Livistona australis* | SCBGBG942 | Duc Thanh Le, Yu-qu Zhang | South China Botanical GardenBG |
| *Masoala madagascariensis* | XMBG217 | Duc Thanh Le, Yu-qu Zhang | Xiamen Botanical Garden |
| *Nannorrhops ritchieana* | XMBG254 | Duc Thanh Le, Yu-qu Zhang | Xiamen Botanical Garden |
| *Nenga pumila* var. *pachystachya* | SCBG962 | Duc Thanh Le, Yu-qu Zhang | South China Botanical Garden |
| *Nephrosperma vanhoutteanum* | XTBG103 | Duc Thanh Le, Yu-qu Zhang | Xishuangbanna Tropical Botanical Garden |
| *Normanbya normanbyi* | SCBG046 | Duc Thanh Le, Yu-qu Zhang | South China Botanical Garden |
| *Oncosperma tigillarium* | MDBG13 | Craig Barrett | Fairchild Tropical Botanic Garden |
| *Oraniopsis appendiculata* | SCBG133 | Duc Thanh Le, Yu-qu Zhang | South China Botanical Garden |
| *Pelagodoxa henryana* | XTBG232 | Duc Thanh Le, Yu-qu Zhang | Xishuangbanna Tropical Botanical Garden |
| *Phoenicophorium borsigianum* | XTBG103 | Duc Thanh Le, Yu-qu Zhang | Xishuangbanna Tropical Botanical Garden |
| *Phoenix acaulis* | XMBG084 | Duc Thanh Le, Yu-qu Zhang | Xiamen Botanical Garden |
| *Phoenix canariensis* | XTBG115 | Duc Thanh Le, Yu-qu Zhang | Xishuangbanna Tropical Botanical Garden |
| *Phoenix dactylifera* | SCBG056 | Duc Thanh Le, Yu-qu Zhang | South China Botanical Garden |
| *Phoenix loureiroi* | XTBG007 | Duc Thanh Le, Yu-qu Zhang | Xishuangbanna Tropical Botanical Garden |
| *Phoenix pusilla* | SCBG065 | Duc Thanh Le, Yu-qu Zhang | South China Botanical Garden |
| *Phoenix reclinata* | SCBG057 | Duc Thanh Le, Yu-qu Zhang | South China Botanical Garden |
| *Phoenix roebelenii* | XTBG142 | Duc Thanh Le, Yu-qu Zhang | Xishuangbanna Tropical Botanical Garden |
| *Phoenix rupicola* | XTBG113 | Duc Thanh Le, Yu-qu Zhang | Xishuangbanna Tropical Botanical Garden |
| *Phoenix sylvestris* | XTBG116 | Duc Thanh Le, Yu-qu Zhang | Xishuangbanna Tropical Botanical Garden |
| *Phoenix theophrasti* | XTBG112 | Duc Thanh Le, Yu-qu Zhang | Xishuangbanna Tropical Botanical Garden |
| *Phytelephas aequatorialis* | XTBG172 | Duc Thanh Le, Yu-qu Zhang | Xishuangbanna Tropical Botanical Garden |
| *Pinanga baviensis* | SCBG142 | Duc Thanh Le, Yu-qu Zhang | South China Botanical Garden |
| *Pinanga gracilis* | XTBG036 | Duc Thanh Le, Yu-qu Zhang | Xishuangbanna Tropical Botanical Garden |
| *Pinanga sylvestris* | XTBG026 | Duc Thanh Le, Yu-qu Zhang | Xishuangbanna Tropical Botanical Garden |
| *Plectocomia elongata* | XTBG077 | Duc Thanh Le, Yu-qu Zhang | Xishuangbanna Tropical Botanical Garden |
| *Pritchardia pacifica* | XTBG015 | Duc Thanh Le, Yu-qu Zhang | Xishuangbanna Tropical Botanical Garden |
| *Ptychosperma macarthurii* | SCBG889 | Duc Thanh Le, Yu-qu Zhang | South China Botanical Garden |
| *Raphia vinifera* | SCBG949 | Duc Thanh Le, Yu-qu Zhang | South China Botanical Garden |
| *Ravenea rivularis* | SCBG029 | Duc Thanh Le, Yu-qu Zhang | South China Botanical Garden |
| *Reinhardtia gracilis* | SCBG060 | Duc Thanh Le, Yu-qu Zhang | South China Botanical Garden |
| *Rhapidophyllum hystrix* | XTBG163 | Duc Thanh Le, Yu-qu Zhang | Xishuangbanna Tropical Botanical Garden |
| *Rhapis excelsa* | SCBG988 | Duc Thanh Le, Yu-qu Zhang | South China Botanical Garden |
| *Roystonea regia* | SCBG136 | Duc Thanh Le, Yu-qu Zhang | South China Botanical Garden |
| *Sabal causiarum* | SCBG035 | Duc Thanh Le, Yu-qu Zhang | South China Botanical Garden |
| *Sabal domingensis* | XTBG168 | Duc Thanh Le, Yu-qu Zhang | Xishuangbanna Tropical Botanical Garden |
| *Sabal etonia* | XMBG407 | Duc Thanh Le, Yu-qu Zhang | Xiamen Botanical Garden |
| *Sabal maritima* | SCBG027 | Duc Thanh Le, Yu-qu Zhang | South China Botanical Garden |
| *Sabal mauritiiformis* | XTBG111 | Duc Thanh Le, Yu-qu Zhang | Xishuangbanna Tropical Botanical Garden |
| *Sabal mexicana* | SCBG053 | Duc Thanh Le, Yu-qu Zhang | South China Botanical Garden |
| *Sabal minor* | SCBG096 | Duc Thanh Le, Yu-qu Zhang | South China Botanical Garden |
| *Sabal palmetto* | SCBG076 | Duc Thanh Le, Yu-qu Zhang | South China Botanical Garden |
| *Sabal princeps* | SCBG091 | Duc Thanh Le, Yu-qu Zhang | South China Botanical Garden |
| *Sabal rosei* | SCBG137 | Duc Thanh Le, Yu-qu Zhang | South China Botanical Garden |
| *Sabal uresana* | XMBG157 | Duc Thanh Le, Yu-qu Zhang | Xiamen Botanical Garden |
| *Salacca wallichiana* | XTBG148 | Duc Thanh Le, Yu-qu Zhang | Xishuangbanna Tropical Botanical Garden |
| *Satakentia liukiuensis* | XTBG236 | Duc Thanh Le, Yu-qu Zhang | Xishuangbanna Tropical Botanical Garden |
| *Schippia concolor* | SCBG945 | Duc Thanh Le, Yu-qu Zhang | South China Botanical Garden |
| *Serenoa repens* | XMBG182 | Duc Thanh Le, Yu-qu Zhang | Xiamen Botanical Garden |
| *Syagrus coronata* | SCBG022 | Duc Thanh Le, Yu-qu Zhang | South China Botanical Garden |
| *Syagrus weddelliana* | SCBG965 | Duc Thanh Le, Yu-qu Zhang | South China Botanical Garden |
| *Thrinax excelsa* | SCBG063 | Duc Thanh Le, Yu-qu Zhang | South China Botanical Garden |
| *Trachycarpus fortunei* | XTBG169 | Duc Thanh Le, Yu-qu Zhang | Xishuangbanna Tropical Botanical Garden |
| *Trachycarpus martianus* | SCBG947 | Duc Thanh Le, Yu-qu Zhang | South China Botanical Garden |
| *Trachycarpus nanus* | XTBG167 | Duc Thanh Le, Yu-qu Zhang | Xishuangbanna Tropical Botanical Garden |
| *Trachycarpus oreophilus* | XMBG214 | Duc Thanh Le, Yu-qu Zhang | Xiamen Botanical Garden |
| *Trachycarpus princeps* | XMBG226 | Duc Thanh Le, Yu-qu Zhang | Xiamen Botanical Garden |
| *Trithrinax campestris* | SCBG080 | Duc Thanh Le, Yu-qu Zhang | South China Botanical Garden |
| *Veitchia joannis* | SCBG088 | Duc Thanh Le, Yu-qu Zhang | South China Botanical Garden |
| *Veitchia subdisticha* | XTBG233 | Duc Thanh Le, Yu-qu Zhang | Xishuangbanna Tropical Botanical Garden |
| *Verschaffeltia splendida* | XTBG102 | Duc Thanh Le, Yu-qu Zhang | Xishuangbanna Tropical Botanical Garden |
| *Wallichia caryotoides* | XTBG060 | Duc Thanh Le, Yu-qu Zhang | Xishuangbanna Tropical Botanical Garden |
| *Wallichia disticha* | XMBG022 | Duc Thanh Le, Yu-qu Zhang | Xiamen Botanical Garden |
| *Wallichia gracilis* | SCBG016 | Duc Thanh Le, Yu-qu Zhang | South China Botanical Garden |
| *Wallichia oblongifolia* | SCBG007 | Duc Thanh Le, Yu-qu Zhang | South China Botanical Garden |
| *Washingtonia filifera* | SCBG881 | Duc Thanh Le, Yu-qu Zhang | South China Botanical Garden |
| *Wodyetia bifurcata* | SCBG034 | Duc Thanh Le, Yu-qu Zhang | South China Botanical Garden |
| *Zombia antillarum* | XTBG216 | Duc Thanh Le, Yu-qu Zhang | Xishuangbanna Tropical Botanical Garden |

**Table S1:** Accessions and voucher information for palms sampled in the genome skimming dataset.

**Table S2:** Genome size (a) and precipitation of the driest month (b) bins used for chi-squared analysis

(a)

| **Quartile** | **Genome size bin (Gbp/1C)** |
| --- | --- |
| *1* | 0.5 < 1.81 |
| *2* | 1.81 < 2.72 |
| *3* | 2.72 < 4.64 |
| *4* | 4.64 < 32 |

(b)

| **Quartile** | **Precipitation of the Driest Month bin (mm)** | **Terminology** | |
| --- | --- | --- | --- |
| *1* | 0 <  34.56 | Very dry | Dry |
| *2* | 34.56 < 64.88 | Semi dry |  |
| *3* | 64.88 < 109.47 | Semi wet | Wet |
| *4* | 109.47 < 290 | Wet |  |

| ***Satellite**** | | | | | | | |
| --- | --- | --- | --- | --- | --- | --- | --- |
| **rDNA** | *5S rDNA** | | | | | | |
|  | 45S rDNA | *18S rDNA** | | | | | |
|  |  | *25S rDNA** | | | | | |
| **Mobile element** | Class I | *Pararetrovirus** | | | | | |
|  |  | *LINE** | | | | | |
|  |  | Long tandem repeat (LTR) | *Ty1/copia* | *Ale** | | | |
|  |  |  |  | *Alesia** | | | |
|  |  |  |  | *Angela** | | | |
|  |  |  |  | *Bianca** | | | |
|  |  |  |  | *Ikeros** | | | |
|  |  |  |  | *Ivana** | | | |
|  |  |  |  | *SIRE** | | | |
|  |  |  |  | *TAR** | | | |
|  |  |  |  | *Tork** | | | |
|  |  |  | *Ty3/gypsy* | Chromovirus | *CRM** | | |
|  |  |  |  |  | *Galadriel** | | |
|  |  |  |  |  | *Reina** | | |
|  |  |  |  |  | *Tekay** | | |
|  |  |  |  | Non-chromovirus | OTA | *Athila** | |
|  |  |  |  |  |  | Tat | *Ogre** |
|  |  |  |  |  |  |  | *Retand** |
|  | Class II | Subclass 2 | *Helitron** | | | | |
|  |  | Subclass 1 | *TIR* | *EnSpm CACTA** | | | |
|  |  |  |  | *hAT** | | | |
|  |  |  |  | *MuDR Mutator** | | | |
|  |  |  |  | *PIF Harbinger** | | | |

**Table S3:** Hierarchical groupings of repeat lineages. The highest groups in the hierarchy are on the left side of the table, decreasing towards the right. Names marked with an asterisk (*) are those which were defined to the lowest hierarchical level in the REXdb database (Neumann *et al.,* 2019) and which we defined as our repeat ‘families’ in this study.

|  | **Estimate** | **Standard Error** | ***t* value** | ***P*r(>\|t\|)** |
| --- | --- | --- | --- | --- |
| ***(Intercept)*** | **0.795** | **0.325** | **2.447** | **0.015** |
| ***Precipitation of the driest month*** | **0.001** | **0.0004** | **3.435** | **0.001** |
| *Min. temperature of the coldest month* | -0.001 | 0.0006 | -1.924 | 0.055 |

**Table S4:** Model summary for minimum adequate PGLS. model explaining variation in log(Genome size) across the Arecaceae, excluding the four polyploid palm species. Significant terms (*P*<0.05) are indicated in bold.

|  | **Estimate** | **Standard error** | ***t* value** | ***P*r(>\|t\|)** |
| --- | --- | --- | --- | --- |
| *Intercept* | -3.006 | 0.133 | -22.608 | <2.2x10^-16^ |
| ***Total repeat occupancy*** | **-0.019** | **0.001** | **-10.215** | **<2.2x10^-16^** |
| ***Genome size*** | **-0.202** | **0.041** | **-4.863** | **3.21x10^-6^** |
| ***Total repeat occupancy:Genome size*** | **0.002** | **0.0006** | **3.419** | **0.0008** |

**Table S5:** Model summary for minimum adequate PGLS model explaining variation in repeat type richness (log(Mehinick's Index)) as a function of genome size and total genome occupancy of repeats across the Arecaceae. Significant terms (*P*<0.05) are indicated in bold.

**Methods S1**

Genome size measurement

For flow cytometry, a fresh palm leaf sample (c. 1cm^2^) was chopped together with the internal standard selected (*Solanum lycopersicum* ‘Stupiké polní rané’ (2C=2.02 pg; (Doležel *et al.,* 1992)), *Petroselinum crispum* 'Champion Moss Curled' (2C=4.50 pg; (Obermayer *et al.,* 2002)), and *Pisum sativum* ‘Ctirad’ (2C=9.09 pg; Doležel *et al.* (1992))) using a razor blade in a petri dish containing 2 mL of ‘general purpose isolation buffer’ (GPB; Loureiro et al. ( 2007))), supplemented with 3% PVP-40 and 15mM β-mercaptoethanol. The homogenate was filtered through a 30 μm nylon mesh to discard debris, stained with 100 μl of PI (1mg/mL, Sigma) and incubated for 20 min on ice. For each accession analysed, three independent samples were prepared and run on the flow cytometer. The nuclear DNA content of each sample run was estimated by recording at least 5,000 particles (c.1,000 nuclei per fluorescence peak) using a Cyflow SL3 flow cytometer (Sysmex-Partec GmbH, Munster, Germany) fitted with a 100-mW green solid-state laser (Cobolt Samba). Resulting output histograms were analysed using the FlowMax software (v. 2.9, Sysmex-Partec GmbH) for statistical calculations. We used only estimates for samples where the coefficients of variation (CV%) of the sample and standard peaks in the flow histogram were less than 5%.

Calculating repeat type richness, occupancy and diversity

We generated community ecology metrics for each palm species from the *RepeatExplorer2* repeat profile dataset in the *R* package *vegan* (Oksanen *et al.,* 2019). We first calculated the richness of different repeat families in each genome using Menhinick’s index (*MI*), which controls for sampling effort by dividing the number of species (*S*) by the square root of the number of individuals sampled (*N*). In our case, we divided the number of repeat families in a genome by the total number of reads assigned to all repeat families by *RepeatExplorer2*. The formula for Menhinick’s Index is as follows:

$$MI=\frac{S}{\surd N}$$

We then calculated the total occupancy of all repeat families in each genome (i.e., % of the genome taken up by all repeats) by dividing the total number of reads assigned to all repeat families by the total number of reads analysed by *RepeatExplorer2* for each species.

Finally, we calculated the diversity of repeat families within each genome using Shannon’s index *(H)*, which accounts both for species richness and species evenness. In a similar fashion to Menhinick’s index, we used the number of reads assigned to repeat families as ‘individuals’, and the repeat families themselves as ‘species’. The formula for Shannon’s Index (*H*) is as follows:

$$H= -\sum_{i=1}^{s} pi\ln pi$$

Where *S* is the total number of species in the community, *pi* is the proportion of S made up of the *i*th species, and *ln* is the natural log. In addition, patterns of genome size, repeat richness (Menhinick’s index), total genome occupancy and repeat diversity (Shannon’s index) were visualised for the palm species under study using the *R* package *ggplot2* (Wickham, 2016).
