## Supporting Information 3 for "The Ecology of Palm Genomes: Repeat-associated genome size expansion is constrained by aridity": 1_RepeatExplorer2_output_analysis_wrapper.sh

```
#Analysing repeat profiles from RepeatExplorer2 - wrapper around output preparation scripts and R script to analyse output from RepeatExplorer2
#Rowan Schley 03/12/2020


#### REMEMBER - CHANGE THIS SCRIPT DEPENDING ON WHICH SPP. YOURE DOING (E.G., all palms, just Phoenix, etc)


### Prepare RepeatExplorer2 output files for analysis ###

#Rename inputs:
#Inputs need to have the following format(note the double underscore): Genus_species__ReadNumber_GenomeSize_CLUSTER_TABLE.csv

cd /mnt/shared/projects/rbgk/projects/rschley_repeatexplorer/Results_RE2_10pct_Craig_Barrett/Type1
for f in *RE2;
do (mv ${f}/CLUSTER_TABLE.csv ${f}/${f}_CLUSTER_TABLE.csv);
done


cd /mnt/shared/projects/rbgk/projects/rschley_repeatexplorer/Results_RE2_10pct_XueJunGe/Type1
for f in *RE2;
do (mv ${f}/CLUSTER_TABLE.csv ${f}/${f}_CLUSTER_TABLE.csv);
done


#Now let's add read numbers and genome sizes to each file
#Get N reads analysed for each file
cd /mnt/shared/projects/rbgk/projects/rschley_repeatexplorer/Results_RE2_10pct_Craig_Barrett/Type1
for f in *RE2;
do (echo $f >> N_reads_analyzed_CraigBarrett_Type1.txt | grep "Number_of_analyzed_reads" $f/${f}_CLUSTER_TABLE.csv | awk '{print $2}' >> N_reads_analyzed_CraigBarrett_Type1.txt);
done


cd /mnt/shared/projects/rbgk/projects/rschley_repeatexplorer/Results_RE2_10pct_XueJunGe/Type1
for f in *RE2;
do (echo $f >> N_reads_analyzed_XueJunGe_Type1.txt | grep "Number_of_analyzed_reads" $f/${f}_CLUSTER_TABLE.csv | awk '{print $2}' >> N_reads_analyzed_XueJunGe_Type1.txt);
done


##### Note: Here we need to make a table of genome sizes and number of reads for each sample (just copy the folder names within each type1/type2/type3/type4 folder).
#E.g.:
#Phoenix-acaulis-PL27_S10__RE2	1	911186
#Phoenix-caespitosa-PL22_S5__RE2	1	978000
#Phoenix-loureiroi-var-pedunculata-PL26_S9__RE2	0.98	958440
#Phoenix-pusilla-PL25_S8__RE2	1	824118
#Phoenix-roebelenii-PL19_S3__RE2	0.72	704160
#Phoenix-sylvestris-PL23_S6__RE2	0.84	747280

cd /mnt/shared/projects/rbgk/projects/rschley_repeatexplorer/Results_RE2_10pct_Craig_Barrett/Type1
awk '{printf "%s%s",$0,NR%2?"\t":RS}' N_reads_analyzed_CraigBarrett_Type1.txt > CraigBarrett_genome_sizes_reads_sampled.txt
##### now add genome sizes by hand to middle column

cd /mnt/shared/projects/rbgk/projects/rschley_repeatexplorer/Results_RE2_10pct_XueJunGe/Type1
awk '{printf "%s%s",$0,NR%2?"\t":RS}' N_reads_analyzed_XueJunGe_Type1.txt > XueJunGe_genome_sizes_reads_sampled.txt
#####now add genome sizes by hand to middle column


#Now let's copy files into directory above and change the file names to contain "Genus_species__ReadNumber_GenomeSize_CLUSTER_TABLE.csv"
cd /mnt/shared/projects/rbgk/projects/rschley_repeatexplorer/Results_RE2_10pct_Craig_Barrett/Type1
for f in *_RE2
do(cp $f/${f}_CLUSTER_TABLE.csv ./);
done 
parallel --eta --colsep '\t' "mv {1}_CLUSTER_TABLE.csv {1}_CraigBarrett__{3}_{2}_CLUSTER_TABLE.csv" :::: ./CraigBarrett_genome_sizes_reads_sampled.txt


cd /mnt/shared/projects/rbgk/projects/rschley_repeatexplorer/Results_RE2_10pct_XueJunGe/Type1
for f in *_RE2
do(cp $f/${f}_CLUSTER_TABLE.csv ./);
done 
parallel --eta --colsep '\t' "mv {1}_CLUSTER_TABLE.csv {1}_XueJunGe__{3}_{2}_CLUSTER_TABLE.csv" :::: ./XueJunGe_genome_sizes_reads_sampled.txt


#Move all CLUSTER_TABLEs into the same directory:
cd /mnt/shared/projects/rbgk/projects/rschley_repeatexplorer/Results_RE2_10pct_Craig_Barrett/All_results_prepped_CraigBarrett/
mv ../Type1/*CLUSTER_TABLE.csv ./

cd ../../Results_RE2_10pct_XueJunGe/All_results_prepped_XueJunGe/
mv ../Type1/*CLUSTER_TABLE.csv ./


#Now we can prepare our input for R.
#First, let's remove the first block of text from each file
cd /mnt/shared/projects/rbgk/projects/rschley_repeatexplorer/Results_RE2_10pct_Craig_Barrett/All_results_prepped_CraigBarrett/
for f in *_CLUSTER_TABLE.csv;
do (tail -n +7 $f > ${f/_CLUSTER_TABLE.csv}_CTonly.csv);
done


cd ../../Results_RE2_10pct_XueJunGe/All_results_prepped_XueJunGe/
for f in *_CLUSTER_TABLE.csv;
do (tail -n +7 $f > ${f/_CLUSTER_TABLE.csv}_CTonly.csv);
done


#Now let's add a species column for each species/filename
cd /mnt/shared/projects/rbgk/projects/rschley_repeatexplorer/Results_RE2_10pct_Craig_Barrett/All_results_prepped_CraigBarrett/
for f in *_CTonly.csv;
do (sed -i "s/$/\t${f/_CTonly.csv}/" $f);
done


cd ../../Results_RE2_10pct_XueJunGe/All_results_prepped_XueJunGe/
for f in *_CTonly.csv;
do (sed -i "s/$/\t${f/_CTonly.csv}/" $f);
done


#Now, let's combine all "_CTonly.csv" files
cd /mnt/shared/projects/rbgk/projects/rschley_repeatexplorer/Results_RE2_10pct_Craig_Barrett/All_results_prepped_CraigBarrett/
cat *CTonly.csv > All_clusters_RE2_CraigBarrett_FINAL.txt

cd ../../Results_RE2_10pct_XueJunGe/All_results_prepped_XueJunGe/
cat *CTonly.csv > All_clusters_RE2_XueJunGe_FINAL.txt


##### Note: Now we have to manually:
##### 1)Check all RE2 output concatenated in the last stage and put the results in the 'final annotation' column #####
##### 2)Create two new columns on the end with n reads and genome size using text to columns in excel (plus label species column as 'Species' #####
																						####	MAKE SURE TO KEEP NREADS AND GENOME SIZES IN NAME ^^^
##### 3)Remove TAREAN annotation column #####
##### 4)Add 'NA' to columns with no final annotation #####


### Prepare input and run analysis in my R script: ###
#Use Analysing_repeatexplorer_output_final_analysis.R
```
