## Supporting Information 3 for "The Ecology of Palm Genomes: Repeat-associated genome size expansion is constrained by aridity": 2_Analysing_repeatexplorer_output_final_analysis.R

```
#Analysing repeat profiles from RepeatExplorer2 for final analysis (genome size and environment)
#Or 'using community ecology metrics and analyses to gain insight on repeat evolution'
#Rowan Schley 03/12/2020

#Set working directory
setwd("C:/Users/rowst/OneDrive - The Royal Botanic Gardens, Kew/PROJECTS/PALM GENOME SIZES/Data analysis/RepeatExplorer/Rowans_analyses/ANALYSING_RE2_OUTPUT/Clustering/Analyses/All_palms_FINAL/")


### REMEMBER TO CHANGE THE SCRIPT DEPENDING ON WHICH SPECIES YOURE ANALYSIS (e.g. all species, just Phoenix) ###


#Read in the vegan package, a great package for community ecology analyses
library(vegan)
library(stringr)
library(tidyverse)
library(reshape2)
library(ggplot2)
library(textshape)
library(ggfortify)
library(ggpubr)
library(gridExtra)
library(caper)
library(phytools)
library(geiger)
library(corrplot)
library(factoextra)
library(RColorBrewer)
library(gridExtra)
library(ggstance)
library(car)
library(ggtree)
library(grid)


####################################################################################### 
		   #################### DATA PROCESSING #####################				  
####################################################################################### 

#Read in data, combine them and remove organellar reads
AP_CB<-read.table("../../Data/All_palms/All_clusters_RE2_CraigBarrett_FINAL.txt", header=T)
AP_CB[,length(AP_CB)+1]<-"Craig_Barrett"
colnames(AP_CB)[length(AP_CB)]<-"Data_source"
head(AP_CB)         
         
AP_XJG<-read.table("../../Data/All_palms/All_clusters_RE2_XueJunGe_FINAL.txt", header=T)
AP_XJG[,length(AP_XJG)+1]<-"Xue_Jun_Ge"
colnames(AP_XJG)[length(AP_XJG)]<-"Data_source"
head(AP_XJG)         

All_palms_GS<-rbind(AP_CB, AP_XJG)
head(All_palms_GS)


#Rename synonyms and mislabelled species (from ../../Data/All_palms/Data_palm_accepted_name_correction.csv)
All_palms_GS$Species<-str_replace(All_palms_GS$Species, "Acrocomia_intrumescens","Acrocomia_intumescens")
All_palms_GS$Species<-str_replace(All_palms_GS$Species, "Aeterospathe_elata","Heterospathe_elata")
All_palms_GS$Species<-str_replace(All_palms_GS$Species, "Aiphanes_caryotifolia","Aiphanes_horrida")
All_palms_GS$Species<-str_replace(All_palms_GS$Species, "Alloschmidia_glabrata","Basselinia_glabrata")
All_palms_GS$Species<-str_replace(All_palms_GS$Species, "Brahea_berlandieri","Brahea_dulcis")
All_palms_GS$Species<-str_replace(All_palms_GS$Species, "Brahea_brandeggeei","Brahea_brandegeei")
All_palms_GS$Species<-str_replace(All_palms_GS$Species, "Brahea_clara","Brahea_armata")
All_palms_GS$Species<-str_replace(All_palms_GS$Species, "Brahea_elegans","Brahea_armata")
All_palms_GS$Species<-str_replace(All_palms_GS$Species, "Brahea_scarukhunji","Brahea_sarukhanii")
All_palms_GS$Species<-str_replace(All_palms_GS$Species, "Chamaedorea_cataractatum","Chamaedorea_cataractarum")
All_palms_GS$Species<-str_replace(All_palms_GS$Species, "Copernicia_baileyna","Copernicia_baileyana")
All_palms_GS$Species<-str_replace(All_palms_GS$Species, "Daemonorops_jenkinsiana","Calamus_jenkinsianus")
All_palms_GS$Species<-str_replace(All_palms_GS$Species, "Drymophloeus_subdistichus","Veitchia_subdisticha")
All_palms_GS$Species<-str_replace(All_palms_GS$Species, "Gronophyllum_microcarpum","Hydriastele_microcarpa")
All_palms_GS$Species<-str_replace(All_palms_GS$Species, "Hydriastele_microspadix","Hydriastele_wendlandiana")
All_palms_GS$Species<-str_replace(All_palms_GS$Species, "Lanonia_hainamensis","Lanonia_hainanensis")
All_palms_GS$Species<-str_replace(All_palms_GS$Species, "Lytocaryum_weddellianum","Syagrus_weddelliana")
All_palms_GS$Species<-str_replace(All_palms_GS$Species, "Nephrosperma_van-houtteanum","Nephrosperma_vanhoutteanum")
All_palms_GS$Species<-str_replace(All_palms_GS$Species, "Normanbya_normanby","Normanbya_normanbyi")
All_palms_GS$Species<-str_replace(All_palms_GS$Species, "Normanbya_normanbyii","Normanbya_normanbyi")
All_palms_GS$Species<-str_replace(All_palms_GS$Species, "Phoenix_ana_var_formosana","Phoenix_loureiroi")
All_palms_GS$Species<-str_replace(All_palms_GS$Species, "Phoenix_humilis","Phoenix_loureiroi")
All_palms_GS$Species<-str_replace(All_palms_GS$Species, "Pinanga_sinii","Pinanga_baviensis")
All_palms_GS$Species<-str_replace(All_palms_GS$Species, "Roscheria_melanochaetes-qerminated","Roscheria_melanochaetes")
All_palms_GS$Species<-str_replace(All_palms_GS$Species, "Sabal_blackburniana","Sabal_palmetto")
All_palms_GS$Species<-str_replace(All_palms_GS$Species, "Sabal_deolbata","Sabal_dealbata")
All_palms_GS$Species<-str_replace(All_palms_GS$Species, "Sabal_jamaicaensis","Sabal_maritima")
All_palms_GS$Species<-str_replace(All_palms_GS$Species, "Sabal_louisiana","Sabal_lougheediana")
All_palms_GS$Species<-str_replace(All_palms_GS$Species, "Sabal_princeps","Sabal_bermudana")
All_palms_GS$Species<-str_replace(All_palms_GS$Species, "Trachycarpus_excelsus","Rhapis_excelsa")
All_palms_GS$Species<-str_replace(All_palms_GS$Species, "Trachycarpus_nana","Trachycarpus_nanus")
All_palms_GS$Species<-str_replace(All_palms_GS$Species, "Victyosperma_album","Dictyosperma_album")
All_palms_GS$Species<-str_replace(All_palms_GS$Species, "Wallichia_chinensis","Wallichia_gracilis")
All_palms_GS$Species<-str_replace(All_palms_GS$Species, "Wallichia_densiflora","Wallichia_oblongifolia")
All_palms_GS$Species<-str_replace(All_palms_GS$Species, "Wallichia_mooreana","Wallichia_caryotoides")
All_palms_GS$Species<-str_replace(All_palms_GS$Species, "Wallichia_siamensis","Wallichia_caryotoides")
All_palms_GS$Species<-str_replace(All_palms_GS$Species, "wodyetia_bifurcata","Wodyetia_bifurcata")


head(All_palms_GS)
All_palms_GS$Species


All_palms_GS<-All_palms_GS[-which(str_detect(All_palms_GS$Final_annotation,"organelle")),]
head(All_palms_GS)
summary(All_palms_GS$Final_annotation) # yep, no more organellar clusters


#Aggregate counts of each repeat classification for each species/treatment (e.g., collapse all Angela element numbers for each spp.)
All_palms_GS_ag<-aggregate(Size~Species+Final_annotation, data=All_palms_GS, FUN=sum) 
head(All_palms_GS_ag)


#Create a suitable table for input to vegan, with annotations of repeats as column headings and species as rownames
All_palms_GS_vegan<-dcast(All_palms_GS_ag, Species ~ Final_annotation)
rownames(All_palms_GS_vegan)<-All_palms_GS_vegan$Species
All_palms_GS_vegan<-All_palms_GS_vegan[,-1]
head(All_palms_GS_vegan)


#Now, let's replace NAs with 0
All_palms_GS_vegan[is.na(All_palms_GS_vegan)] <- 0
head(All_palms_GS_vegan)


#Let's plot number of reads for each species and repeat type
All_palms_GS_vegan %>% 
  as.data.frame() %>%
  rownames_to_column("Species") %>%
  pivot_longer(-c(Species), names_to = "Repeat_type", values_to = "Reads") %>%
  ggplot(aes(x=Repeat_type, y=Species, fill=Reads)) + 
  geom_raster() + 
  theme(axis.text.x = element_text(angle = 90)) + 
  scale_fill_viridis_c()


#Let's try plotting a stacked bar chart for each species and the abundances of different repeats
#Repeat % of all repeat types
ggplot(All_palms_GS_ag, aes(fill=Final_annotation, y=Size, x=Species)) + 
  theme(axis.text.x = element_text(angle = 90, size= 10)) + 
  theme(legend.text = element_text(size=6)) + 
  ylab("Percent of all repeats") + 
  geom_bar(position="fill", stat="identity")


#Absolute repeat number
ggplot(All_palms_GS_ag, aes(fill=Final_annotation, y=Size, x=Species)) + 
  theme(axis.text.x = element_text(angle = 90, size= 10)) + 
  theme(legend.text = element_text(size=6)) + 
  ylab("Number of repeat reads") + 
  geom_bar(stat="identity")


#Repeat % of whole genome
All_palms_GS_ag_pct<-All_palms_GS_ag
All_palms_GS_ag_pct[,4]<-str_split_fixed(All_palms_GS_ag[,1], "__", 2)[,2]
All_palms_GS_ag_pct[,c(4,5)]<-str_split_fixed(All_palms_GS_ag_pct[,4], "_", 2)[,c(1,2)]
head(All_palms_GS_ag_pct)
colnames(All_palms_GS_ag_pct)[c(4,5)]<-c("N_reads_analyzed","Genome_size")
head(All_palms_GS_ag_pct)

All_palms_GS_ag_pct[,6]<- All_palms_GS_ag_pct[,3] / as.numeric(unlist(All_palms_GS_ag_pct[,4])) * 100
colnames(All_palms_GS_ag_pct)[6]<-"Repeat_pct_whole_genome"
head(All_palms_GS_ag_pct)

ggplot(All_palms_GS_ag_pct, aes(fill=Final_annotation, y=Repeat_pct_whole_genome, x=Species)) + 
  theme(axis.text.x = element_text(angle = 90, size= 10)) + 
  theme(legend.text = element_text(size=6)) + 
  ylab("Percentage occupied within the genome") + 
  ylim(0,100) + 
  geom_bar(position="stack", stat="identity")


####################################################################################### 
     #################### CALCULATING DIVERSITY INDICES #####################				  
####################################################################################### 

## Richness (Mehinick's Index) - number of species (n)/sqrt(total number of individuals (N)) ##
#Important to take into account sample size because, of course, the more individuals we survey the more species there will be (until a point)
#Closer to 1 = higher richness
n <- apply(All_palms_GS_vegan>0,1,sum) # number of repeat types
rnames<-rownames(All_palms_GS_vegan)
All_palms_GS_vegan<-apply(All_palms_GS_vegan[,c(1:28)], 2, as.numeric)
rownames(All_palms_GS_vegan)<-rnames
N <- apply(All_palms_GS_vegan,1,sum) # total number of reads

#Since we aimed to get the same genome proportion across species, the correlation between mehinicks index and genome size is valid, as the 'sampling effort'
#(in other words, genome size or how many reads we sampled proportionally) is likely not causing lower richness in bigger genomes?
#That is, assuming you sample less, if the richness is the same it should be so even despite lower numbers of 'individuals' (hits) in each 'species' (repeat class).


MIdf<-n/sqrt(N)
MIdf<-as.data.frame(MIdf)
MIdf[,ncol(MIdf)+1]<-gsub(" ", "",paste(str_split_fixed(rownames(MIdf), "_", 3)[,1], "_", str_split_fixed(rownames(MIdf), "_", 3)[,2]))
colnames(MIdf)[ncol(MIdf)-1]<-"Mehinicks_index"
colnames(MIdf)[ncol(MIdf)]<-"Tree_name"
MIdf[,ncol(MIdf)+1]<-rownames(MIdf)
colnames(MIdf)[ncol(MIdf)]<-"Species"
head(MIdf)


################################################################## ##
## Nucleotide similarity - % shared bases between two repeats of the same type between genomes
#See Hs/Ho scripts


##################################################################
## Occupancy - % of the genome occupied by each repeat type ##
#Split columns to give genome size and number of reads examined
Ocdf <- as.data.frame(All_palms_GS_vegan)
Ocdf[,ncol(Ocdf)+1] <- str_split_fixed(rownames(All_palms_GS_vegan), "__", 2)[,2]
Ocdf[,c(ncol(Ocdf)+1,ncol(Ocdf)+2)] <- str_split_fixed(Ocdf[,as.numeric(length(Ocdf))], "_", 2)[,c(1,2)]
head(Ocdf)
colnames(Ocdf)[c((length(Ocdf)-1),length(Ocdf))] <- c("N_reads_analyzed", "Genome_size")
head(Ocdf)
Ocdf<-Ocdf[,-(ncol(Ocdf)-2)]
head(Ocdf)

#Make a numeric data frame for the conversion
Ocdf_numeric<- transform(Ocdf, as.numeric())
sapply(Ocdf_numeric, class)#Let's see if it's actually numeric now...
head(Ocdf_numeric)

#Create data frame with number of reads for each repeat type divided by number of reads analyzed * 100 
#(i.e., genome occupancy of each repeat type in a sample)
Repeat_occupancy<- Ocdf_numeric[,c(1:(length(Ocdf_numeric)-2))] / as.numeric(unlist(Ocdf_numeric[,length(Ocdf_numeric)-1])) * 100
head(Repeat_occupancy)
head(Ocdf) # to compare calculated percentages to actual dataframe

#Let's make a column with total genome occupancy per species, and make a species column
Repeat_occupancy2<-Repeat_occupancy
Repeat_occupancy2[,ncol(Repeat_occupancy2)+1]<-rowSums(Repeat_occupancy)
head(Repeat_occupancy2)
colnames(Repeat_occupancy2)[ncol(Repeat_occupancy2)]<-"Genome_occupancy"
Repeat_occupancy2[,ncol(Repeat_occupancy2)+1]<-rownames(Repeat_occupancy2)
colnames(Repeat_occupancy2)[ncol(Repeat_occupancy2)]<-"Species"
head(Repeat_occupancy2)
Genome_occupancy<-Repeat_occupancy2[,c(ncol(Repeat_occupancy2), ncol(Repeat_occupancy2)-1)]
head(Genome_occupancy)


##################################################################
## Diversity - Shannon's diversity index ##
#Let's calculate Shannon's diversity index for our vegan dataframe 
#(i.e., likelihood random individual belongs to a certain species)
Shannon_All_palms<-diversity(All_palms_GS_vegan, index = "shannon")
Shannon_All_palms


#Let's compare Shannon indices
SPdf<-as.data.frame(Shannon_All_palms)
SPdf[,2]<-rownames(SPdf)
colnames(SPdf)[c(1,2)]<-c("Shannon_Index","Species")
head(SPdf)


#Create column with Tree_name
SPdf[,ncol(SPdf)+1]<-gsub(" ", "",paste(str_split_fixed(rownames(SPdf), "_", 3)[,1], "_", str_split_fixed(rownames(SPdf), "_", 3)[,2]))
colnames(SPdf)[ncol(SPdf)]<-"Tree_name"
head(SPdf)


####################################################################################### 
  #################### MODELLING REPEAT PROFILES AND ENVIRONMENT #####################				  
####################################################################################### 

#Read in tree, environment and GS data, combine with current data by making 'tree names' column to enable matching
#Add 'Tree_names' (i.e., species) column to Palm_GS_vegan to allow matching of all datasets (and remove spaces with gsub)
All_palms_GS_vegan<-as.data.frame(All_palms_GS_vegan)
All_palms_GS_vegan[,ncol(All_palms_GS_vegan)+1]<-gsub(" ", "",paste(str_split_fixed(rownames(All_palms_GS_vegan), "_", 3)[,1], "_", str_split_fixed(rownames(All_palms_GS_vegan), "_", 3)[,2]))
colnames(All_palms_GS_vegan)[ncol(All_palms_GS_vegan)]<-"Tree_name"
head(All_palms_GS_vegan)


#Get genome size and number of reads analysed from species names
All_palms_GS_vegan_pct<-All_palms_GS_vegan
All_palms_GS_vegan_pct[,ncol(All_palms_GS_vegan_pct)+1]<-str_split_fixed(rownames(All_palms_GS_vegan_pct), "__", 2)[,2]
All_palms_GS_vegan_pct[,c(ncol(All_palms_GS_vegan_pct)+1, ncol(All_palms_GS_vegan_pct)+2)]<-str_split_fixed(All_palms_GS_vegan_pct[,ncol(All_palms_GS_vegan_pct)], "_", 2)[,c(1,2)]
head(All_palms_GS_vegan_pct)
colnames(All_palms_GS_vegan_pct)[c(ncol(All_palms_GS_vegan_pct)-1, ncol(All_palms_GS_vegan_pct))]<-c("N_reads_analyzed","Genome_size")
All_palms_GS_vegan_pct<-All_palms_GS_vegan_pct[,-(ncol(All_palms_GS_vegan_pct)-2)]
All_palms_GS_vegan_pct[,ncol(All_palms_GS_vegan_pct)+1]<-rownames(All_palms_GS_vegan_pct)
colnames(All_palms_GS_vegan_pct)[ncol(All_palms_GS_vegan_pct)]<-"Species"
head(All_palms_GS_vegan_pct)


##### CRITICALLY IMPORTANT - TURN ALL READ NUMBERS INTO GENOME PERCENTAGES FOR ALL SPECIES FOR EACH REPEAT CLASS ####
#To do this, divide number of reads in each repeat class by number of reads analysed in total
#All_palms_GS_vegan_pct[,c(1:(ncol(All_palms_GS_vegan_pct)-4))]<- All_palms_GS_vegan_pct[,c(1:(ncol(All_palms_GS_vegan_pct)-4))] / as.numeric(unlist(All_palms_GS_vegan_pct$N_reads_analyzed)) * 100
#head(All_palms_GS_vegan_pct)
#dim(All_palms_GS_vegan_pct)


#Read in tree
phy<-read.tree("C:/Users/rowst/OneDrive - The Royal Botanic Gardens, Kew/PROJECTS/PALM GENOME SIZES/Data collection/Trees/Phylogeny_NoCon_Henderson_Tree_1_.new")
tip2drop<-which(duplicated(phy$tip.label)) #ID duplicated tips
phyclean<-drop.tip(phy= phy, tip = tip2drop) #Remove duplicated tips


#Worldclim
climdat<- read.csv("C:/Users/rowst/OneDrive - The Royal Botanic Gardens, Kew/PROJECTS/PALM GENOME SIZES/Data collection/Trait data/Worldclim/Per_sp_all_worldclim.csv", header = T)
colnames(climdat)[1]<-"Tree_name"
head(climdat)

#Let's see which species we don't have in the worldclim dataset 
which(!All_palms_GS_vegan_pct$Tree_name%in%climdat$Tree_name)


#Combine everything into one big per-species dataframe
####IMPORTANT - before doing this ensure we have data for all species and there aren't spelling errors or synonyms
#Shannon index data and main repeat profile vegan dataset
ALL_DATA_palms_GS_vegan_pct<-merge(All_palms_GS_vegan_pct, SPdf, by="Species")
head(ALL_DATA_palms_GS_vegan_pct)
ALL_DATA_palms_GS_vegan_pct<-ALL_DATA_palms_GS_vegan_pct[,-ncol(ALL_DATA_palms_GS_vegan_pct)]
colnames(ALL_DATA_palms_GS_vegan_pct)[(ncol(ALL_DATA_palms_GS_vegan_pct)-3)]<-"Tree_name"
head(ALL_DATA_palms_GS_vegan_pct)

#Merge with Mehinicks index (richness) dataset
ALL_DATA_palms_GS_vegan_pct<-merge(ALL_DATA_palms_GS_vegan_pct, MIdf, by="Species")
head(ALL_DATA_palms_GS_vegan_pct)
ALL_DATA_palms_GS_vegan_pct<-ALL_DATA_palms_GS_vegan_pct[,-ncol(ALL_DATA_palms_GS_vegan_pct)]
colnames(ALL_DATA_palms_GS_vegan_pct)[(ncol(ALL_DATA_palms_GS_vegan_pct)-4)]<-"Tree_name"
head(ALL_DATA_palms_GS_vegan_pct)
dim(ALL_DATA_palms_GS_vegan_pct)

#Now merge with genome occupancy data
ALL_DATA_palms_GS_vegan_pct<-merge(ALL_DATA_palms_GS_vegan_pct, Genome_occupancy, by="Species")
head(ALL_DATA_palms_GS_vegan_pct)
dim(ALL_DATA_palms_GS_vegan_pct)

#Finally, merge all of the above with worldclim data
ALL_DATA_palms_GS_vegan_pct<-merge(ALL_DATA_palms_GS_vegan_pct, climdat, by="Tree_name")
head(ALL_DATA_palms_GS_vegan_pct)
dim(ALL_DATA_palms_GS_vegan_pct)
ALL_DATA_palms_GS_vegan_pct$Genome_size_gbp<-as.numeric(ALL_DATA_palms_GS_vegan_pct$Genome_size)*0.978
head(ALL_DATA_palms_GS_vegan_pct)


write.csv(file="ALL_DATA_palms_GS_vegan.csv", ALL_DATA_palms_GS_vegan_pct)


##~~~~~~~~~~~~~~~~~~~~Exploratory plotting~~~~~~~~~~~~~~~~~~~~## 

#How does richness change with genome size?
plot(ALL_DATA_palms_GS_vegan_pct$Mehinicks_index~ALL_DATA_palms_GS_vegan_pct$Genome_size_gbp, ylab = "Repeat type richness (Mehinicks Index)", xlab="Genome size (Gbp)")

ggplot(ALL_DATA_palms_GS_vegan_pct, aes(x=as.numeric(Genome_size_gbp), y= Mehinicks_index)) + 
  geom_point() +
  geom_smooth(method="loess", span=1) +
  labs(y="Repeat type richness (Mehinicks Index)", x = "Genome size (Gbp)") + 
  theme(text=element_text(size=14),axis.text = element_text(size = 12))


#Let's see if % of the genome occupied by repeats changes with genome size as in Novak et al (2020) - nature plants
plot(ALL_DATA_palms_GS_vegan_pct$Genome_occupancy~ALL_DATA_palms_GS_vegan_pct$Genome_size_gbp, ylab = "% of the Genome occupied by repeats", xlab="Genome size (Gbp)")

#Let's plot it with smoothing in ggplot, which uses the means to plot a line which helps the eye see trends
#(note, not a model, just a visualisation method)
ggplot(ALL_DATA_palms_GS_vegan_pct, aes(x=as.numeric(Genome_size_gbp), y= Genome_occupancy)) + 
  geom_point() +
  geom_smooth(method="loess", span=1) +
  labs(y="% of the genome occupied by repeats", x = "Genome size (Gbp)") + 
  theme(text=element_text(size=14),axis.text = element_text(size = 12))


#How does diversity change with genome size?
plot(ALL_DATA_palms_GS_vegan_pct$Shannon_Index~ALL_DATA_palms_GS_vegan_pct$Genome_size_gbp, ylab = "Repeat diversity (Shannon Index)", xlab="Genome size (Gbp)")

ggplot(ALL_DATA_palms_GS_vegan_pct, aes(x=as.numeric(Genome_size_gbp), y= Shannon_Index)) + 
  geom_point() +
  geom_smooth(method="loess", span=1) +
  labs(y="Repeat type diversity (Shannon Index)", x = "Genome size (Gbp)") + 
  theme(text=element_text(size=14),axis.text = element_text(size = 12))


#What about how richness, occupancy, and diversity change with precip driest month?
ggplot(ALL_DATA_palms_GS_vegan_pct, aes(x=as.numeric(Precipitation_of_Driest_Month), y= Mehinicks_index)) + 
  geom_point() +
  geom_smooth(method="loess", span=1) +
  labs(y="Repeat type diversity (Mehinicks Index)", x = "Precipitation of the driest month") + 
  theme(text=element_text(size=14),axis.text = element_text(size = 12))

ggplot(ALL_DATA_palms_GS_vegan_pct, aes(x=as.numeric(Precipitation_of_Driest_Month), y= Genome_occupancy)) + 
  geom_point() +
  geom_smooth(method="loess", span=1) +
  labs(y="% of the genome occupied by repeats", x = "Precipitation of the driest month") + 
  theme(text=element_text(size=14),axis.text = element_text(size = 12))

ggplot(ALL_DATA_palms_GS_vegan_pct, aes(x=as.numeric(Precipitation_of_Driest_Month), y= Shannon_Index)) + 
  geom_point() +
  geom_smooth(method="loess", span=1) +
  labs(y="Repeat type diversity (Shannon Index)", x = "Precipitation of the driest month") + 
  theme(text=element_text(size=14),axis.text = element_text(size = 12))

ggplot(ALL_DATA_palms_GS_vegan_pct, aes(x=Genome_occupancy, y= Mehinicks_index)) + 
  geom_point() +
  geom_smooth(method="loess", span=1) +
  labs(y="Repeat type richness (Mehinick's index)", x = "% of the genome occupied by repeats") + 
  theme(text=element_text(size=14),axis.text = element_text(size = 12))


##~~~~~~~~~~~~~~~~~~~~PHYLANOVA~~~~~~~~~~~~~~~~~~~~## 
###### Let's see if there are signficiant differences in Richness, Occupancy and Diversity between precipitation classes ######

### Richness model 
#Put data into precipitation groups
colnames(ALL_DATA_palms_GS_vegan_pct)
anova_input<-ALL_DATA_palms_GS_vegan_pct[,c(1,2, 3:35, 49,55)]
head(anova_input)
dim(anova_input)
  
sum_precip<-summary(anova_input$Precipitation_of_Driest_Mon)

q1_precip<-which(anova_input$Precipitation_of_Driest_Month <= sum_precip[[2]])
q2_precip<-which(anova_input$Precipitation_of_Driest_Month >= sum_precip[[2]] & anova_input$Precipitation_of_Driest_Month <= sum_precip[[3]])
q3_precip<-which(anova_input$Precipitation_of_Driest_Month >= sum_precip[[3]] & anova_input$Precipitation_of_Driest_Month <= sum_precip[[5]])
q4_precip<-which(anova_input$Precipitation_of_Driest_Month >= sum_precip[[5]]& anova_input$Precipitation_of_Driest_Month <= sum_precip[[6]])

anova_input[q1_precip,(ncol(anova_input)+1)]<-"Dry"
anova_input[q2_precip,(ncol(anova_input))]<-"Semidry"
anova_input[q3_precip,(ncol(anova_input))]<-"Semiwet"
anova_input[q4_precip,(ncol(anova_input))]<-"Wet"
colnames(anova_input)[ncol(anova_input)]<-"Precip_class"
head(anova_input)


#First, a preliminary ANOVA
richness_anova<-aov(Mehinicks_index~Precip_class, data=anova_input)
summary(richness_anova)
boxplot(Mehinicks_index~Precip_class, data=anova_input)


#Now a phylogenetically controlled ANOVA (phylanova) from phytools
MI_PA<-anova_input[which(anova_input$Tree_name%in%phyclean$tip.label),c(1,34)]
MI_PA<-MI_PA[-which(duplicated(MI_PA$Tree_name)),]
dim(MI_PA)
head(MI_PA)
MI_PA<-column_to_rownames(MI_PA, loc=1)
MI_PA<-as.matrix(MI_PA)[,1]  #### this it is KEY for phytools!
head(MI_PA)

precip_PA<-anova_input[which(anova_input$Tree_name%in%phyclean$tip.label),c(1,37)]
precip_PA<-precip_PA[-which(duplicated(precip_PA$Tree_name)),]
dim(precip_PA)
precip_PA<-column_to_rownames(precip_PA, loc=1)
precip_PA<-as.matrix(precip_PA)[,1]  #### this it is KEY for phytools!
head(precip_PA)

phyclean_anova<-keep.tip(phyclean, rownames(as.matrix(MI_PA)))
phyclean_anova$tip.label

phylANOVA(phyclean_anova, precip_PA, MI_PA)


#Let's try a pgls
mod_input_MI<-ALL_DATA_palms_GS_vegan_pct[,c(1,32,34,49,55)]
colnames(mod_input_MI)
mod_input_MI_unique<-mod_input_MI[-which(duplicated(mod_input_MI$Tree_name)),]
colnames(mod_input_MI_unique)
dim(mod_input_MI_unique)
pgls_dat_MI<-comparative.data(data=mod_input_MI_unique, phy=phyclean, names.col="Tree_name",vcv=TRUE)
print(pgls_dat_MI)

hist(mod_input_MI_unique$Mehinicks_index)
hist(log(mod_input_MI_unique$Mehinicks_index))
shapiro.test(mod_input_MI_unique$Mehinicks_index)
shapiro.test(log(mod_input_MI_unique$Mehinicks_index))
#Logging helps Mehinicks Index


mod_MI_lambda<-pgls(log(Mehinicks_index)~ as.numeric(Genome_size)*Precipitation_of_Driest_Month, data=pgls_dat_MI, lambda = "ML", bounds = list(lambda = c(0.05, 1))) #lambda branch transformation
summary(mod_MI_lambda)
mod_MI_lambda_2<-update(mod_MI_lambda,~.- as.numeric(Genome_size):Precipitation_of_Driest_Month, data=pgls_dat_MI, lambda = "ML", bounds = list(lambda = c(0.05, 1))) #lambda branch transformation
summary(mod_MI_lambda_2)
mod_MI_lambda_3<-update(mod_MI_lambda_2,~.- Precipitation_of_Driest_Month, data=pgls_dat_MI, lambda = "ML", bounds = list(lambda = c(0.5, 1))) #lambda branch transformation
summary(mod_MI_lambda_3)
#Genome size best explains variation (as we know from plotting)


#### Occupancy (H) model -
#First, a preliminary ANOVA
occupancy_anova<-aov(Genome_occupancy~Precip_class, data=anova_input)
summary(occupancy_anova)
boxplot(Genome_occupancy~Precip_class, data=anova_input)


#Now a phylogeneticically controlled ANOVA (phylanova) from phytools
occ_PA<-anova_input[which(anova_input$Tree_name%in%phyclean$tip.label),c(1,35)]
occ_PA<-occ_PA[-which(duplicated(occ_PA$Tree_name)),]
dim(occ_PA)
head(occ_PA)
occ_PA<-column_to_rownames(occ_PA, loc=1)
occ_PA<-as.matrix(occ_PA)[,1]  #### this it is KEY for phytools!
head(occ_PA)

phylANOVA(phyclean_anova, precip_PA, occ_PA)


#Let's try pgls
mod_input_occ<-ALL_DATA_palms_GS_vegan_pct[,c(1,32,35,49,55)]
colnames(mod_input_occ)
mod_input_occ_unique<-mod_input_occ[-which(duplicated(mod_input_occ$Tree_name)),]
colnames(mod_input_occ_unique)
dim(mod_input_occ_unique)
pgls_dat_occ<-comparative.data(data=mod_input_occ_unique, phy=phyclean, names.col="Tree_name",vcv=TRUE)
print(pgls_dat_occ)

hist(mod_input_occ_unique$Genome_occupancy)
shapiro.test(mod_input_occ_unique$Genome_occupancy)
#Fine without transformation


pmod_occ_lambda<-pgls(Genome_occupancy~ as.numeric(Genome_size)*Precipitation_of_Driest_Month, data=pgls_dat_occ, lambda = "ML", bounds = list(lambda = c(0.005, 1))) #lambda branch transformation
summary(pmod_occ_lambda)
pmod_occ_lambda_2<-update(mod_occ_lambda,~.- as.numeric(Genome_size):Precipitation_of_Driest_Month, data=pgls_dat_occ, lambda = "ML", bounds = list(lambda = c(0.005, 1))) #lambda branch transformation
summary(pmod_occ_lambda_2)
pmod_occ_lambda_3<-update(mod_occ_lambda_2,~.- Precipitation_of_Driest_Month, data=pgls_dat_occ, lambda = "ML", bounds = list(lambda = c(0.005, 1))) #lambda branch transformation
summary(pmod_occ_lambda_3)

AIC(pmodd_occ_lambda, pmod_occ_lambda_2, pmod_occ_lambda_3)
#Genome size best explains variation (as we already know from plotting)


#### Diversity (H) model -
#First, a preliminary ANOVA
diversity_anova<-aov(Shannon_Index~Precip_class, data=anova_input)
summary(diversity_anova)
boxplot(Shannon_Index~Precip_class, data=anova_input)


#Now a phylogeneticically controlled ANOVA (phylanova) from phytools
SI_PA<-anova_input[which(anova_input$Tree_name%in%phyclean$tip.label),c(1,33)]
SI_PA<-SI_PA[-which(duplicated(SI_PA$Tree_name)),]
dim(SI_PA)
head(SI_PA)
SI_PA<-column_to_rownames(SI_PA, loc=1)
SI_PA<-as.matrix(SI_PA)[,1]  #### this it is KEY for phytools!
head(SI_PA)

phylANOVA(phyclean_anova, precip_PA, SI_PA)


#Let's try pgls
mod_input_shan<-ALL_DATA_palms_GS_vegan_pct[,c(1,32,33,49,55)]
colnames(mod_input_shan)
mod_input_shan_unique<-mod_input_shan[-which(duplicated(mod_input_shan$Tree_name)),]
colnames(mod_input_shan_unique)
dim(mod_input_shan_unique)
pgls_dat_shan<-comparative.data(data=mod_input_shan_unique, phy=phyclean, names.col="Tree_name",vcv=TRUE)
print(pgls_dat_shan)

hist(mod_input_shan_unique$Shannon_Index)
shapiro.test(mod_input_shan_unique$Shannon_Index)
#Fine without transformation


mod_shan_lambda<-pgls(Shannon_Index~ Genome_size_gbp*Precipitation_of_Driest_Month, data=pgls_dat_shan, lambda = "ML", bounds = list(lambda = c(0.005, 1))) #lambda branch transformation
summary(mod_shan_lambda)
mod_shan_lambda_2<-update(mod_shan_lambda,~.- Genome_size_gbp:Precipitation_of_Driest_Month, data=pgls_dat_shan, lambda = "ML", bounds = list(lambda = c(0.005, 1))) #lambda branch transformation
summary(mod_shan_lambda_2)
mod_shan_lambda_3<-update(mod_shan_lambda_2,~.- Genome_size_gbp, data=pgls_dat_shan, lambda = "ML", bounds = list(lambda = c(0.005, 1))) #lambda branch transformation
summary(mod_shan_lambda_3)
#Nothing significant (as we know from plotting)


## And Genome size, mehinicks index and genome occupancy?
mod_input_MI_occ_unique<-merge(mod_input_MI_unique, mod_input_occ_unique, by="Tree_name")
mod_input_MI_occ_unique<-mod_input_MI_occ_unique[,-c(4,5)]
head(mod_input_MI_occ_unique)
dim(mod_input_MI_occ_unique)
pgls_dat_MI_occ<-comparative.data(data=mod_input_MI_occ_unique, phy=phyclean, names.col="Tree_name",vcv=TRUE)
print(pgls_dat_MI_occ)

mod_MI_occ_lambda<-pgls(log(Mehinicks_index)~ Genome_occupancy*as.numeric(Genome_size_gbp.y)*Precipitation_of_Driest_Month.y, data=pgls_dat_MI_occ, lambda = "ML", bounds = list(lambda = c(0.005, 1))) #lambda branch transformation
summary(mod_MI_occ_lambda)
mod_MI_occ_lambda_2<-update(mod_MI_occ_lambda,~.- Genome_occupancy:as.numeric(Genome_size_gbp.y):Precipitation_of_Driest_Month.y, data=pgls_dat_MI_occ, lambda = "ML", bounds = list(lambda = c(0.005, 1))) #lambda branch transformation
summary(mod_MI_occ_lambda_2)
mod_MI_occ_lambda_3<-update(mod_MI_occ_lambda_2,~.- as.numeric(Genome_size_gbp.y):Precipitation_of_Driest_Month.y, data=pgls_dat_MI_occ, lambda = "ML", bounds = list(lambda = c(0.005, 1))) #lambda branch transformation
summary(mod_MI_occ_lambda_3)
mod_MI_occ_lambda_4<-update(mod_MI_occ_lambda_3,~.- Genome_occupancy:Precipitation_of_Driest_Month.y, data=pgls_dat_MI_occ, lambda = "ML", bounds = list(lambda = c(0.005, 1))) #lambda branch transformation
summary(mod_MI_occ_lambda_4)
mod_MI_occ_lambda_5<-update(mod_MI_occ_lambda_4,~.- Precipitation_of_Driest_Month.y, data=pgls_dat_MI_occ, lambda = "ML", bounds = list(lambda = c(0.005, 1))) #lambda branch transformation
summary(mod_MI_occ_lambda_5)


###PCA
#Let's plot a PCA to see how different species are in terms of their repeat classifications, and colour it by precipitation class
df <- anova_input[,c(3:30,37)]
pca_res <- prcomp(df[,c(1:28)], scale. = TRUE)
autoplot(pca_res, data=df, colour="Precip_class")


##~~~~~~~~~~~~~~~~~~~~PGLS - REPEAT CLASSES AND GENOME SIZE ~~~~~~~~~~~~~~~~~~~~## 
###### Let's try to see whether different repeat types significantly impact genome size (as in Macas et al 2015) #######
#We will only use original data without exons, introns etc removed (type1)
########################################################################################################################

#let's write a simple linear model to see if genome size correlates with different repeat types in all variables
#Test with simple lm
#Log to normalise genome size response
colnames(ALL_DATA_palms_GS_vegan_pct)
mod_input_GS<-ALL_DATA_palms_GS_vegan_pct[,c(1:30,32,55)]
colnames(mod_input_GS)
#Convert to gigabases
mod_input_GS[,c(3:30)]<-mod_input_GS[,c(3:30)]/10000000


mod_input_GS_fin<-mod_input_GS[,-c(1,2)]
colnames(mod_input_GS_fin)
dim(mod_input_GS_fin)
mod_1_GS<-lm(log(as.numeric(Genome_size))~., data=mod_input_GS_fin)
summary(mod_1_GS)

vif(mod_1_GS) # yay! none are correlated

mod_2_GS<-step(mod_1_GS)
summary(mod_2_GS) #Cool, seems interesting.


#Let's try in a phylogenetically-corrected fashion
#PGLS of genome size vs repeat class
mod_input_GS_unique<-mod_input_GS[-which(duplicated(mod_input_GS$Tree_name)),-2]
colnames(mod_input_GS_unique)
dim(mod_input_GS_unique)
pgls_dat_GS<-comparative.data(data=mod_input_GS_unique, phy=phyclean, names.col="Tree_name",vcv=TRUE)
print(pgls_dat_GS)

mod_GS_lambda<-pgls(log(as.numeric(Genome_size_gbp))~ `All/repeat/mobile_element/Class_I/LINE`+`All/repeat/mobile_element/Class_I/LTR/Ty1_copia/Ale`+`All/repeat/mobile_element/Class_I/LTR/Ty1_copia/Alesia`+`All/repeat/mobile_element/Class_I/LTR/Ty1_copia/Angela`+`All/repeat/mobile_element/Class_I/LTR/Ty1_copia/Bianca`+`All/repeat/mobile_element/Class_I/LTR/Ty1_copia/Ikeros`+`All/repeat/mobile_element/Class_I/LTR/Ty1_copia/Ivana`+`All/repeat/mobile_element/Class_I/LTR/Ty1_copia/SIRE`+`All/repeat/mobile_element/Class_I/LTR/Ty1_copia/TAR`+`All/repeat/mobile_element/Class_I/LTR/Ty1_copia/Tork`+`All/repeat/mobile_element/Class_I/LTR/Ty3_gypsy/chromovirus/CRM`+`All/repeat/mobile_element/Class_I/LTR/Ty3_gypsy/chromovirus/Galadriel`+`All/repeat/mobile_element/Class_I/LTR/Ty3_gypsy/chromovirus/Reina`+`All/repeat/mobile_element/Class_I/LTR/Ty3_gypsy/chromovirus/Tekay`+`All/repeat/mobile_element/Class_I/LTR/Ty3_gypsy/non-chromovirus/OTA`+`All/repeat/mobile_element/Class_I/LTR/Ty3_gypsy/non-chromovirus/OTA/Athila`+`All/repeat/mobile_element/Class_I/LTR/Ty3_gypsy/non-chromovirus/OTA/Tat/Ogre`+`All/repeat/mobile_element/Class_I/LTR/Ty3_gypsy/non-chromovirus/OTA/Tat/Retand`+`All/repeat/mobile_element/Class_I/pararetrovirus`+`All/repeat/mobile_element/Class_II/Subclass_1/TIR/EnSpm_CACTA`+`All/repeat/mobile_element/Class_II/Subclass_1/TIR/hAT`+`All/repeat/mobile_element/Class_II/Subclass_1/TIR/MuDR_Mutator`+`All/repeat/mobile_element/Class_II/Subclass_1/TIR/PIF_Harbinger`+`All/repeat/rDNA/45S_rDNA/18S_rDNA`+`All/repeat/rDNA/45S_rDNA/25S_rDNA`+`All/repeat/rDNA/5S_rDNA`+`All/repeat/satellite`, data=pgls_dat_GS, lambda = "ML", bounds = list(lambda = c(0.005, 1))) #lambda branch transformation
summary(mod_GS_lambda)

mod_GS_lambda_2<-update(mod_GS_lambda,~. -`All/repeat/mobile_element/Class_I/LTR/Ty3_gypsy/chromovirus/Reina`, lambda = 'ML', bounds = list(lambda = c(0.05, 1))) #lambda branch transformation
summary(mod_GS_lambda_2)
mod_GS_lambda_3<-update(mod_GS_lambda_2,~. -`All/repeat/mobile_element/Class_I/LINE`, lambda = 'ML', bounds = list(lambda = c(0.05, 1))) #lambda branch transformation
summary(mod_GS_lambda_3)
mod_GS_lambda_4<-update(mod_GS_lambda_3,~. -`All/repeat/mobile_element/Class_I/LTR/Ty3_gypsy/chromovirus/Galadriel`, lambda = 'ML', bounds = list(lambda = c(0.05, 1))) #lambda branch transformation
summary(mod_GS_lambda_4)
mod_GS_lambda_5<-update(mod_GS_lambda_4,~. -`All/repeat/mobile_element/Class_I/LTR/Ty3_gypsy/non-chromovirus/OTA/Athila`, lambda = 'ML', bounds = list(lambda = c(0.05, 1))) #lambda branch transformation
summary(mod_GS_lambda_5)
mod_GS_lambda_6<-update(mod_GS_lambda_5,~. -`All/repeat/mobile_element/Class_I/LTR/Ty3_gypsy/non-chromovirus/OTA/Tat/Ogre`, lambda = 'ML', bounds = list(lambda = c(0.05, 1))) #lambda branch transformation
summary(mod_GS_lambda_6)
mod_GS_lambda_7<-update(mod_GS_lambda_6,~. -`All/repeat/rDNA/45S_rDNA/18S_rDNA`, lambda = 'ML', bounds = list(lambda = c(0.05, 1))) #lambda branch transformation
summary(mod_GS_lambda_7)
mod_GS_lambda_8<-update(mod_GS_lambda_7,~. -`All/repeat/mobile_element/Class_I/LTR/Ty1_copia/Tork`, lambda = 'ML', bounds = list(lambda = c(0.05, 1))) #lambda branch transformation
summary(mod_GS_lambda_8)
mod_GS_lambda_9<-update(mod_GS_lambda_8,~. -`All/repeat/mobile_element/Class_I/LTR/Ty1_copia/Ivana`, lambda = 'ML', bounds = list(lambda = c(0.09, 1))) #lambda branch transformation
summary(mod_GS_lambda_9)
mod_GS_lambda_10<-update(mod_GS_lambda_9,~. -`All/repeat/satellite` , lambda = 'ML', bounds = list(lambda = c(0.05, 1))) #lambda branch transformation
summary(mod_GS_lambda_10)
mod_GS_lambda_11<-update(mod_GS_lambda_10,~. -`All/repeat/rDNA/45S_rDNA/25S_rDNA`, lambda = 'ML', bounds = list(lambda = c(0.05, 1))) #lambda branch transformation
summary(mod_GS_lambda_11)
mod_GS_lambda_12<-update(mod_GS_lambda_11,~. -`All/repeat/mobile_element/Class_II/Subclass_1/TIR/PIF_Harbinger`, lambda = 'ML', bounds = list(lambda = c(0.05, 1))) #lambda branch transformation
summary(mod_GS_lambda_12)
mod_GS_lambda_13<-update(mod_GS_lambda_12,~. -`All/repeat/mobile_element/Class_I/LTR/Ty1_copia/SIRE`, lambda = 'ML', bounds = list(lambda = c(0.09, 1))) #lambda branch transformation
summary(mod_GS_lambda_13)
mod_GS_lambda_14<-update(mod_GS_lambda_13,~. -`All/repeat/rDNA/5S_rDNA`, lambda = 'ML', bounds = list(lambda = c(0.05, 1))) #lambda branch transformation
summary(mod_GS_lambda_14)
mod_GS_lambda_15<-update(mod_GS_lambda_14,~. -`All/repeat/mobile_element/Class_I/LTR/Ty1_copia/Ale`, lambda = 'ML', bounds = list(lambda = c(0.05, 1))) #lambda branch transformation
summary(mod_GS_lambda_15)
mod_GS_lambda_16<-update(mod_GS_lambda_15,~. -`All/repeat/mobile_element/Class_I/LTR/Ty3_gypsy/non-chromovirus/OTA`, lambda = 'ML', bounds = list(lambda = c(0.05, 1))) #lambda branch transformation
summary(mod_GS_lambda_16)
mod_GS_lambda_17<-update(mod_GS_lambda_16,~. -`All/repeat/mobile_element/Class_I/LTR/Ty1_copia/Ikeros`, lambda = 'ML', bounds = list(lambda = c(0.05, 1))) #lambda branch transformation
summary(mod_GS_lambda_17)
mod_GS_lambda_18<-update(mod_GS_lambda_17,~. -`All/repeat/mobile_element/Class_I/LTR/Ty1_copia/Bianca`, lambda = 'ML', bounds = list(lambda = c(0.05, 1))) #lambda branch transformation
summary(mod_GS_lambda_18)
mod_GS_lambda_19<-update(mod_GS_lambda_18,~. -`All/repeat/mobile_element/Class_II/Subclass_1/TIR/hAT`, lambda = 'ML', bounds = list(lambda = c(0.05, 1))) #lambda branch transformation
summary(mod_GS_lambda_19)
mod_GS_lambda_20<-update(mod_GS_lambda_19,~. -`All/repeat/mobile_element/Class_I/LTR/Ty1_copia/Alesia`, lambda = 'ML', bounds = list(lambda = c(0.05, 1))) #lambda branch transformation
summary(mod_GS_lambda_20)
mod_GS_lambda_21<-update(mod_GS_lambda_20,~. -`All/repeat/mobile_element/Class_I/pararetrovirus`, lambda = 'ML', bounds = list(lambda = c(0.05, 1))) #lambda branch transformation
summary(mod_GS_lambda_21)
mod_GS_lambda_22<-update(mod_GS_lambda_21,~. -`All/repeat/mobile_element/Class_II/Subclass_1/TIR/MuDR_Mutator`, lambda = 'ML', bounds = list(lambda = c(0.05, 1))) #lambda branch transformation
summary(mod_GS_lambda_22)

AIC(mod_GS_lambda_22,mod_GS_lambda_21,mod_GS_lambda_20,mod_GS_lambda_19,mod_GS_lambda_18,mod_GS_lambda_17,mod_GS_lambda_16,mod_GS_lambda_15,mod_GS_lambda_14,mod_GS_lambda_13,mod_GS_lambda_12,mod_GS_lambda_11,mod_GS_lambda_10,mod_GS_lambda_9,mod_GS_lambda_8,mod_GS_lambda_7,mod_GS_lambda_6,mod_GS_lambda_5,mod_GS_lambda_4,mod_GS_lambda_3,mod_GS_lambda_2,mod_GS_lambda)
summary(mod_GS_lambda_18)


#Let's try with ABSOLUTE repeat values
ALL_DATA_palms_GS_vegan_abs<-read.csv("ALL_DATA_palms_GS_vegan_absol_vals.csv", header=T,row.names = 1)
head(ALL_DATA_palms_GS_vegan_abs)
mod_input_GS_abs<-ALL_DATA_palms_GS_vegan_abs[,c(1:30,32)]
mod_input_GS_abs_fin<-mod_input_GS_abs[,-c(1,2)]
colnames(mod_input_GS_abs_fin)
dim(mod_input_GS_abs_fin)


#Let's try in a phylogenetically-corrected fashion
#PGLS of genome size vs repeat class
mod_input_GS_abs_unique<-mod_input_GS_abs[-which(duplicated(mod_input_GS_abs$Tree_name)),-2]
colnames(mod_input_GS_abs_unique)
dim(mod_input_GS_abs_unique)
pgls_dat_GS_abs<-comparative.data(data=mod_input_GS_abs_unique, phy=phyclean, names.col="Tree_name",vcv=TRUE)
print(pgls_dat_GS_abs)

mod_GS_abs_lambda<-pgls(log(as.numeric(Genome_size))~ All.repeat.mobile_element.Class_I.LINE+All.repeat.mobile_element.Class_I.LTR.Ty1_copia.Ale+All.repeat.mobile_element.Class_I.LTR.Ty1_copia.Alesia+All.repeat.mobile_element.Class_I.LTR.Ty1_copia.Angela+All.repeat.mobile_element.Class_I.LTR.Ty1_copia.Bianca+All.repeat.mobile_element.Class_I.LTR.Ty1_copia.Ikeros+All.repeat.mobile_element.Class_I.LTR.Ty1_copia.Ivana+All.repeat.mobile_element.Class_I.LTR.Ty1_copia.SIRE+All.repeat.mobile_element.Class_I.LTR.Ty1_copia.TAR+All.repeat.mobile_element.Class_I.LTR.Ty1_copia.Tork+All.repeat.mobile_element.Class_I.LTR.Ty3_gypsy.chromovirus.CRM+All.repeat.mobile_element.Class_I.LTR.Ty3_gypsy.chromovirus.Galadriel+All.repeat.mobile_element.Class_I.LTR.Ty3_gypsy.chromovirus.Reina+All.repeat.mobile_element.Class_I.LTR.Ty3_gypsy.chromovirus.Tekay+All.repeat.mobile_element.Class_I.LTR.Ty3_gypsy.non_chromovirus.OTA+All.repeat.mobile_element.Class_I.LTR.Ty3_gypsy.non_chromovirus.OTA.Athila+All.repeat.mobile_element.Class_I.LTR.Ty3_gypsy.non_chromovirus.OTA.Tat.Ogre+All.repeat.mobile_element.Class_I.LTR.Ty3_gypsy.non_chromovirus.OTA.Tat.Retand+All.repeat.mobile_element.Class_I.pararetrovirus+All.repeat.mobile_element.Class_II.Subclass_1.TIR.EnSpm_CACTA+All.repeat.mobile_element.Class_II.Subclass_1.TIR.hAT+All.repeat.mobile_element.Class_II.Subclass_1.TIR.MuDR_Mutator+All.repeat.mobile_element.Class_II.Subclass_1.TIR.PIF_Harbinger+All.repeat.rDNA.45S_rDNA.18S_rDNA+All.repeat.rDNA.45S_rDNA.25S_rDNA+All.repeat.rDNA.5S_rDNA+All.repeat.satellite, data=pgls_dat_GS_abs, lambda = "ML", bounds = list(lambda = c(0.005, 1))) #lambda branch transformation
summary(mod_GS_abs_lambda)

mod_GS_abs_lambda_2<-update(mod_GS_abs_lambda,~. -All.repeat.mobile_element.Class_I.LTR.Ty3_gypsy.chromovirus.Reina, lambda = 'ML', bounds = list(lambda = c(0.005, 1))) #lambda branch transformation
summary(mod_GS_abs_lambda_2)
mod_GS_abs_lambda_3<-update(mod_GS_abs_lambda_2,~. -All.repeat.mobile_element.Class_I.LINE, lambda = 'ML', bounds = list(lambda = c(0.005, 1))) #lambda branch transformation
summary(mod_GS_abs_lambda_3)
mod_GS_abs_lambda_4<-update(mod_GS_abs_lambda_3,~. -All.repeat.mobile_element.Class_I.LTR.Ty3_gypsy.chromovirus.Galadriel, lambda = 'ML', bounds = list(lambda = c(0.005, 1))) #lambda branch transformation
summary(mod_GS_abs_lambda_4)
mod_GS_abs_lambda_5<-update(mod_GS_abs_lambda_4,~. -All.repeat.mobile_element.Class_I.LTR.Ty3_gypsy.non_chromovirus.OTA.Athila, lambda = 'ML', bounds = list(lambda = c(0.005, 1))) #lambda branch transformation
summary(mod_GS_abs_lambda_5)
mod_GS_abs_lambda_6<-update(mod_GS_abs_lambda_5,~. -All.repeat.mobile_element.Class_I.LTR.Ty3_gypsy.non_chromovirus.OTA.Tat.Ogre, lambda = 'ML', bounds = list(lambda = c(0.005, 1))) #lambda branch transformation
summary(mod_GS_abs_lambda_6)
mod_GS_abs_lambda_7<-update(mod_GS_abs_lambda_6,~. -All.repeat.rDNA.45S_rDNA.18S_rDNA, lambda = 'ML', bounds = list(lambda = c(0.005, 1))) #lambda branch transformation
summary(mod_GS_abs_lambda_7)
mod_GS_abs_lambda_8<-update(mod_GS_abs_lambda_7,~. -All.repeat.mobile_element.Class_I.LTR.Ty1_copia.Tork, lambda = 'ML', bounds = list(lambda = c(0.005, 1))) #lambda branch transformation
summary(mod_GS_abs_lambda_8)
mod_GS_abs_lambda_9<-update(mod_GS_abs_lambda_8,~. -All.repeat.mobile_element.Class_I.LTR.Ty1_copia.Ivana, lambda = 'ML', bounds = list(lambda = c(0.09, 1))) #lambda branch transformation
summary(mod_GS_abs_lambda_9)
mod_GS_abs_lambda_10<-update(mod_GS_abs_lambda_9,~. -All.repeat.satellite, lambda = 'ML', bounds = list(lambda = c(0.005, 1))) #lambda branch transformation
summary(mod_GS_abs_lambda_10)
mod_GS_abs_lambda_11<-update(mod_GS_abs_lambda_10,~. -All.repeat.rDNA.45S_rDNA.25S_rDNA, lambda = 'ML', bounds = list(lambda = c(0.005, 1))) #lambda branch transformation
summary(mod_GS_abs_lambda_11)
mod_GS_abs_lambda_12<-update(mod_GS_abs_lambda_11,~. -All.repeat.mobile_element.Class_II.Subclass_1.TIR.PIF_Harbinger, lambda = 'ML', bounds = list(lambda = c(0.005, 1))) #lambda branch transformation
summary(mod_GS_abs_lambda_12)
mod_GS_abs_lambda_13<-update(mod_GS_abs_lambda_12,~. -All.repeat.mobile_element.Class_I.LTR.Ty1_copia.SIRE, lambda = 'ML', bounds = list(lambda = c(0.09, 1))) #lambda branch transformation
summary(mod_GS_abs_lambda_13)
mod_GS_abs_lambda_14<-update(mod_GS_abs_lambda_13,~. -All.repeat.rDNA.5S_rDNA, lambda = 'ML', bounds = list(lambda = c(0.005, 1))) #lambda branch transformation
summary(mod_GS_abs_lambda_14)
mod_GS_abs_lambda_15<-update(mod_GS_abs_lambda_14,~. -All.repeat.mobile_element.Class_I.LTR.Ty1_copia.Ale, lambda = 'ML', bounds = list(lambda = c(0.005, 1))) #lambda branch transformation
summary(mod_GS_abs_lambda_15)
mod_GS_abs_lambda_16<-update(mod_GS_abs_lambda_15,~. -All.repeat.mobile_element.Class_I.LTR.Ty3_gypsy.non_chromovirus.OTA, lambda = 'ML', bounds = list(lambda = c(0.005, 1))) #lambda branch transformation
summary(mod_GS_abs_lambda_16)
mod_GS_abs_lambda_17<-update(mod_GS_abs_lambda_16,~. -All.repeat.mobile_element.Class_I.LTR.Ty1_copia.Ikeros, lambda = 'ML', bounds = list(lambda = c(0.005, 1))) #lambda branch transformation
summary(mod_GS_abs_lambda_17)
mod_GS_abs_lambda_18<-update(mod_GS_abs_lambda_17,~. -All.repeat.mobile_element.Class_I.LTR.Ty1_copia.Bianca, lambda = 'ML', bounds = list(lambda = c(0.005, 1))) #lambda branch transformation
summary(mod_GS_abs_lambda_18)
mod_GS_abs_lambda_19<-update(mod_GS_abs_lambda_18,~. -All.repeat.mobile_element.Class_II.Subclass_1.TIR.hAT, lambda = 'ML', bounds = list(lambda = c(0.005, 1))) #lambda branch transformation
summary(mod_GS_abs_lambda_19)
mod_GS_abs_lambda_20<-update(mod_GS_abs_lambda_19,~. -All.repeat.mobile_element.Class_I.LTR.Ty1_copia.Alesia, lambda = 'ML', bounds = list(lambda = c(0.005, 1))) #lambda branch transformation
summary(mod_GS_abs_lambda_20)
mod_GS_abs_lambda_21<-update(mod_GS_abs_lambda_20,~. -All.repeat.mobile_element.Class_I.pararetrovirus, lambda = 'ML', bounds = list(lambda = c(0.005, 1))) #lambda branch transformation
summary(mod_GS_abs_lambda_21)
mod_GS_abs_lambda_22<-update(mod_GS_abs_lambda_21,~. -All.repeat.mobile_element.Class_II.Subclass_1.TIR.MuDR_Mutator, lambda = 'ML', bounds = list(lambda = c(0.005, 1))) #lambda branch transformation
summary(mod_GS_abs_lambda_22)

AIC(mod_GS_abs_lambda_22,mod_GS_abs_lambda_21,mod_GS_abs_lambda_20,mod_GS_abs_lambda_19,mod_GS_abs_lambda_18,mod_GS_abs_lambda_17,mod_GS_abs_lambda_16,mod_GS_abs_lambda_15,mod_GS_abs_lambda_14,mod_GS_abs_lambda_13,mod_GS_abs_lambda_12,mod_GS_abs_lambda_11,mod_GS_abs_lambda_10,mod_GS_abs_lambda_9,mod_GS_abs_lambda_8,mod_GS_abs_lambda_7,mod_GS_abs_lambda_6,mod_GS_abs_lambda_5,mod_GS_abs_lambda_4,mod_GS_abs_lambda_3,mod_GS_abs_lambda_2,mod_GS_abs_lambda)
summary(mod_GS_abs_lambda_18)


##~~~~~~~~~~~~~~~~~~~~PGLS - REPEAT CLASSES AND PRECIPITATION DRIEST MONTH ~~~~~~~~~~~~~~~~~~~~## 
###### Now, let's see if any repeat classifications correlate with precipitation driest month ######
#Test with simple lm
#Square root to normalise precipitation of the driest month response
ALL_DATA_palms_GS_vegan_pct$Genome_occupancy<-as.numeric(ALL_DATA_palms_GS_vegan_pct$Genome_occupancy)
mod_input_precip<-ALL_DATA_palms_GS_vegan_pct[,c(1:30,32,49)]
mod_input_precip[,c(3:30)]<-mod_input_precip[,c(3:30)]/10000000
mod_input_precip_fin<-mod_input_precip[,-c(1,2)]
mod_input_precip_fin<-as.data.frame(apply(mod_input_precip_fin, 2, as.numeric))
colnames(mod_input_precip_fin)
dim(mod_input_precip_fin)
mod_1_precip<-lm(sqrt(as.numeric(Precipitation_of_Driest_Month))~., data=mod_input_precip_fin)
summary(mod_1_precip)

mod_2_precip<-step(mod_1_precip)
summary(mod_2_precip) # looks interesting


#Let's try in a phylogenetically-corrected fashion, with genome size in two bins as an interaction term
#(to get different slopes for different genome sizes, since repeat abundance correlates with genome size)
#This is taking an ANCOVA approach, with at least one categorical and at least one continuous predictor variable
#PGLS of precipitation driest month vs repeat class*genome_size

mod_input_precip_unique<-mod_input_precip[-which(duplicated(mod_input_precip$Tree_name)),-2]
mod_input_precip_unique[,-1]<-as.data.frame(apply(mod_input_precip_unique[,-1], 2, as.numeric))
head(mod_input_precip_unique)

#Let's create genome size bins
sum_GS<-summary(mod_input_precip_unique$Genome_size)

q1_GS<-which(mod_input_precip_unique$Genome_size <= sum_GS[[3]])
q2_GS<-which(mod_input_precip_unique$Genome_size > sum_GS[[3]])

mod_input_precip_unique[q1_GS,(ncol(mod_input_precip_unique)+1)]<-"Small"
mod_input_precip_unique[q2_GS,(ncol(mod_input_precip_unique))]<-"Large"
colnames(mod_input_precip_unique)[ncol(mod_input_precip_unique)]<-"Genome_size_class"
head(mod_input_precip_unique)

colnames(mod_input_precip_unique)
dim(mod_input_precip_unique)
pgls_dat_precip<-comparative.data(data=mod_input_precip_unique, phy=phyclean, names.col=Tree_name,vcv=TRUE)
print(pgls_dat_precip)

#remove variables with zero variance, and remember that R takes interactions alphabetically, so the first variables with no interactions are for 'large' genome species
mod_precip_lambda<-pgls(sqrt(as.numeric(Precipitation_of_Driest_Month))~`All/repeat/mobile_element/Class_I/LINE`+`All/repeat/mobile_element/Class_I/LTR/Ty1_copia/Ale`+`All/repeat/mobile_element/Class_I/LTR/Ty1_copia/Alesia`+`All/repeat/mobile_element/Class_I/LTR/Ty1_copia/Angela`+`All/repeat/mobile_element/Class_I/LTR/Ty1_copia/Bianca`+`All/repeat/mobile_element/Class_I/LTR/Ty1_copia/Ikeros`+`All/repeat/mobile_element/Class_I/LTR/Ty1_copia/Ivana`+`All/repeat/mobile_element/Class_I/LTR/Ty1_copia/SIRE`+`All/repeat/mobile_element/Class_I/LTR/Ty1_copia/TAR`+`All/repeat/mobile_element/Class_I/LTR/Ty1_copia/Tork`+`All/repeat/mobile_element/Class_I/LTR/Ty3_gypsy/chromovirus/CRM`+`All/repeat/mobile_element/Class_I/LTR/Ty3_gypsy/chromovirus/Galadriel`+`All/repeat/mobile_element/Class_I/LTR/Ty3_gypsy/chromovirus/Reina`+`All/repeat/mobile_element/Class_I/LTR/Ty3_gypsy/chromovirus/Tekay`+`All/repeat/mobile_element/Class_I/LTR/Ty3_gypsy/non-chromovirus/OTA/Athila`+`All/repeat/mobile_element/Class_I/LTR/Ty3_gypsy/non-chromovirus/OTA/Tat/Ogre`+`All/repeat/mobile_element/Class_I/LTR/Ty3_gypsy/non-chromovirus/OTA/Tat/Retand`+`All/repeat/mobile_element/Class_I/pararetrovirus`+`All/repeat/mobile_element/Class_II/Subclass_1/TIR/EnSpm_CACTA`+`All/repeat/mobile_element/Class_II/Subclass_1/TIR/hAT`+`All/repeat/mobile_element/Class_II/Subclass_1/TIR/MuDR_Mutator`+`All/repeat/mobile_element/Class_II/Subclass_1/TIR/PIF_Harbinger`+`All/repeat/rDNA/45S_rDNA/18S_rDNA`+`All/repeat/rDNA/45S_rDNA/25S_rDNA`+`All/repeat/rDNA/5S_rDNA`+`All/repeat/satellite`+as.factor(Genome_size_class):(`All/repeat/mobile_element/Class_I/LINE`+`All/repeat/mobile_element/Class_I/LTR/Ty1_copia/Ale`+`All/repeat/mobile_element/Class_I/LTR/Ty1_copia/Alesia`+`All/repeat/mobile_element/Class_I/LTR/Ty1_copia/Angela`+`All/repeat/mobile_element/Class_I/LTR/Ty1_copia/Bianca`+`All/repeat/mobile_element/Class_I/LTR/Ty1_copia/Ikeros`+`All/repeat/mobile_element/Class_I/LTR/Ty1_copia/Ivana`+`All/repeat/mobile_element/Class_I/LTR/Ty1_copia/SIRE`+`All/repeat/mobile_element/Class_I/LTR/Ty1_copia/TAR`+`All/repeat/mobile_element/Class_I/LTR/Ty1_copia/Tork`+`All/repeat/mobile_element/Class_I/LTR/Ty3_gypsy/chromovirus/CRM`+`All/repeat/mobile_element/Class_I/LTR/Ty3_gypsy/chromovirus/Galadriel`+`All/repeat/mobile_element/Class_I/LTR/Ty3_gypsy/chromovirus/Reina`+`All/repeat/mobile_element/Class_I/LTR/Ty3_gypsy/chromovirus/Tekay`+`All/repeat/mobile_element/Class_I/LTR/Ty3_gypsy/non-chromovirus/OTA/Athila`+`All/repeat/mobile_element/Class_I/LTR/Ty3_gypsy/non-chromovirus/OTA/Tat/Ogre`+`All/repeat/mobile_element/Class_I/LTR/Ty3_gypsy/non-chromovirus/OTA/Tat/Retand`+`All/repeat/mobile_element/Class_I/pararetrovirus`+`All/repeat/mobile_element/Class_II/Subclass_1/TIR/EnSpm_CACTA`+`All/repeat/mobile_element/Class_II/Subclass_1/TIR/hAT`+`All/repeat/mobile_element/Class_II/Subclass_1/TIR/MuDR_Mutator`+`All/repeat/mobile_element/Class_II/Subclass_1/TIR/PIF_Harbinger`+`All/repeat/rDNA/45S_rDNA/18S_rDNA`+`All/repeat/rDNA/45S_rDNA/25S_rDNA`+`All/repeat/rDNA/5S_rDNA`+`All/repeat/satellite`), data=pgls_dat_precip, lambda = 'ML', bounds = list(lambda = c(0.0005, 1))) #lambda branch transformation
summary(mod_precip_lambda)

mod_precip_lambda_2<-update(mod_precip_lambda,~. -`All/repeat/mobile_element/Class_I/LTR/Ty1_copia/Bianca`:as.factor(Genome_size_class), lambda = 'ML', bounds = list(lambda = c(0.0005, 1))) #lambda branch transformation
summary(mod_precip_lambda_2)
mod_precip_lambda_3<-update(mod_precip_lambda_2,~. -`All/repeat/satellite`:as.factor(Genome_size_class), lambda = 'ML', bounds = list(lambda = c(0.0005, 1))) #lambda branch transformation
summary(mod_precip_lambda_3)
mod_precip_lambda_4<-update(mod_precip_lambda_3,~. -`All/repeat/mobile_element/Class_I/LTR/Ty3_gypsy/chromovirus/CRM`:as.factor(Genome_size_class), lambda = 'ML', bounds = list(lambda = c(0.0005, 1))) #lambda branch transformation
summary(mod_precip_lambda_4)
mod_precip_lambda_5<-update(mod_precip_lambda_4,~. -`All/repeat/mobile_element/Class_I/LTR/Ty1_copia/TAR`:as.factor(Genome_size_class), lambda = 'ML', bounds = list(lambda = c(0.0005, 1))) #lambda branch transformation
summary(mod_precip_lambda_5)
mod_precip_lambda_6<-update(mod_precip_lambda_5,~. -`All/repeat/mobile_element/Class_II/Subclass_1/TIR/MuDR_Mutator`:as.factor(Genome_size_class), lambda = 'ML', bounds = list(lambda = c(0.0005, 1))) #lambda branch transformation
summary(mod_precip_lambda_6)
mod_precip_lambda_7<-update(mod_precip_lambda_6,~. -`All/repeat/mobile_element/Class_I/LTR/Ty1_copia/Alesia`:as.factor(Genome_size_class), lambda = 'ML', bounds = list(lambda = c(0.0005, 1))) #lambda branch transformation
summary(mod_precip_lambda_7)
mod_precip_lambda_8<-update(mod_precip_lambda_7,~. -`All/repeat/rDNA/45S_rDNA/25S_rDNA`:as.factor(Genome_size_class), lambda = 'ML', bounds = list(lambda = c(0.0005, 1))) #lambda branch transformation
summary(mod_precip_lambda_8)
mod_precip_lambda_9<-update(mod_precip_lambda_8,~. -`All/repeat/mobile_element/Class_I/LTR/Ty3_gypsy/non-chromovirus/OTA/Tat/Retand`:as.factor(Genome_size_class), lambda = 'ML', bounds = list(lambda = c(0.0005, 1))) #lambda branch transformation
summary(mod_precip_lambda_9)
mod_precip_lambda_10<-update(mod_precip_lambda_9,~. -`All/repeat/mobile_element/Class_I/LTR/Ty3_gypsy/chromovirus/Reina`:as.factor(Genome_size_class), lambda = 'ML', bounds = list(lambda = c(0.0005, 1))) #lambda branch transformation
summary(mod_precip_lambda_10)
mod_precip_lambda_11<-update(mod_precip_lambda_10,~. -`All/repeat/mobile_element/Class_I/LTR/Ty3_gypsy/non-chromovirus/OTA/Athila`:as.factor(Genome_size_class), lambda = 'ML', bounds = list(lambda = c(0.0005, 1))) #lambda branch transformation
summary(mod_precip_lambda_11)
mod_precip_lambda_12<-update(mod_precip_lambda_11,~. -`All/repeat/mobile_element/Class_II/Subclass_1/TIR/PIF_Harbinger`:as.factor(Genome_size_class), lambda = 'ML', bounds = list(lambda = c(0.0005, 1))) #lambda branch transformation
summary(mod_precip_lambda_12)
mod_precip_lambda_13<-update(mod_precip_lambda_12,~. -`All/repeat/mobile_element/Class_II/Subclass_1/TIR/hAT`:as.factor(Genome_size_class), lambda = 'ML', bounds = list(lambda = c(0.0005, 1))) #lambda branch transformation
summary(mod_precip_lambda_13)
mod_precip_lambda_14<-update(mod_precip_lambda_13,~. -`All/repeat/mobile_element/Class_I/LTR/Ty1_copia/Ale`:as.factor(Genome_size_class), lambda = 'ML', bounds = list(lambda = c(0.0005, 1))) #lambda branch transformation
summary(mod_precip_lambda_14)
mod_precip_lambda_15<-update(mod_precip_lambda_14,~. -`All/repeat/mobile_element/Class_I/LTR/Ty1_copia/Ivana`:as.factor(Genome_size_class), lambda = 'ML', bounds = list(lambda = c(0.0005, 1))) #lambda branch transformation
summary(mod_precip_lambda_15)
mod_precip_lambda_16<-update(mod_precip_lambda_15,~. -`All/repeat/mobile_element/Class_I/pararetrovirus`:as.factor(Genome_size_class), lambda = 'ML', bounds = list(lambda = c(0.0005, 1))) #lambda branch transformation
summary(mod_precip_lambda_16)
mod_precip_lambda_17<-update(mod_precip_lambda_16,~. -`All/repeat/mobile_element/Class_I/LTR/Ty1_copia/Tork`:as.factor(Genome_size_class), lambda = 'ML', bounds = list(lambda = c(0.0005, 1))) #lambda branch transformation
summary(mod_precip_lambda_17)
mod_precip_lambda_18<-update(mod_precip_lambda_17,~. -`All/repeat/mobile_element/Class_I/LTR/Ty3_gypsy/non-chromovirus/OTA/Tat/Ogre`:as.factor(Genome_size_class), lambda = 'ML', bounds = list(lambda = c(0.0005, 1))) #lambda branch transformation
summary(mod_precip_lambda_18)
mod_precip_lambda_19<-update(mod_precip_lambda_18,~. -`All/repeat/mobile_element/Class_I/LTR/Ty3_gypsy/chromovirus/Tekay`:as.factor(Genome_size_class), lambda = 'ML', bounds = list(lambda = c(0.0005, 1))) #lambda branch transformation
summary(mod_precip_lambda_19)
mod_precip_lambda_20<-update(mod_precip_lambda_19,~. -`All/repeat/rDNA/45S_rDNA/18S_rDNA`:as.factor(Genome_size_class), lambda = 'ML', bounds = list(lambda = c(0.0005, 1))) #lambda branch transformation
summary(mod_precip_lambda_20)
mod_precip_lambda_21<-update(mod_precip_lambda_20,~. -`All/repeat/mobile_element/Class_I/LTR/Ty1_copia/Ikeros`:as.factor(Genome_size_class), lambda = 'ML', bounds = list(lambda = c(0.0005, 1))) #lambda branch transformation
summary(mod_precip_lambda_21)
mod_precip_lambda_22<-update(mod_precip_lambda_21,~. -`All/repeat/mobile_element/Class_I/LINE`:as.factor(Genome_size_class), lambda = 'ML', bounds = list(lambda = c(0.0005, 1))) #lambda branch transformation
summary(mod_precip_lambda_22)
mod_precip_lambda_23<-update(mod_precip_lambda_22,~. -`All/repeat/mobile_element/Class_I/LTR/Ty1_copia/SIRE`:as.factor(Genome_size_class), lambda = 'ML', bounds = list(lambda = c(0.0005, 1))) #lambda branch transformation
summary(mod_precip_lambda_23)
mod_precip_lambda_24<-update(mod_precip_lambda_23,~. -`All/repeat/mobile_element/Class_II/Subclass_1/TIR/EnSpm_CACTA`:as.factor(Genome_size_class), lambda = 'ML', bounds = list(lambda = c(0.0005, 1))) #lambda branch transformation
summary(mod_precip_lambda_24)
mod_precip_lambda_25<-update(mod_precip_lambda_24,~. -`All/repeat/mobile_element/Class_I/LTR/Ty3_gypsy/chromovirus/Galadriel`:as.factor(Genome_size_class), lambda = 'ML', bounds = list(lambda = c(0.0005, 1))) #lambda branch transformation
summary(mod_precip_lambda_25)
mod_precip_lambda_26<-update(mod_precip_lambda_25,~. -`All/repeat/rDNA/5S_rDNA`:as.factor(Genome_size_class), lambda = 'ML', bounds = list(lambda = c(0.0005, 1))) #lambda branch transformation
summary(mod_precip_lambda_26)
mod_precip_lambda_27<-update(mod_precip_lambda_26,~. -`All/repeat/mobile_element/Class_II/Subclass_1/TIR/MuDR_Mutator`, lambda = 'ML', bounds = list(lambda = c(0.0005, 1))) #lambda branch transformation
summary(mod_precip_lambda_27)
mod_precip_lambda_28<-update(mod_precip_lambda_27,~. -`All/repeat/mobile_element/Class_I/LTR/Ty1_copia/Alesia`, lambda = 'ML', bounds = list(lambda = c(0.0005, 1))) #lambda branch transformation
summary(mod_precip_lambda_28)
mod_precip_lambda_29<-update(mod_precip_lambda_28,~. -`All/repeat/mobile_element/Class_I/LTR/Ty1_copia/SIRE`, lambda = 'ML', bounds = list(lambda = c(0.0005, 1))) #lambda branch transformation
summary(mod_precip_lambda_29)
mod_precip_lambda_30<-update(mod_precip_lambda_29,~. -`All/repeat/mobile_element/Class_I/LTR/Ty3_gypsy/chromovirus/Galadriel`, lambda = 'ML', bounds = list(lambda = c(0.0005, 1))) #lambda branch transformation
summary(mod_precip_lambda_30)
mod_precip_lambda_31<-update(mod_precip_lambda_30,~. -`All/repeat/mobile_element/Class_I/LTR/Ty3_gypsy/chromovirus/Reina`, lambda = 'ML', bounds = list(lambda = c(0.0005, 1))) #lambda branch transformation
summary(mod_precip_lambda_31)
mod_precip_lambda_32<-update(mod_precip_lambda_31,~. -`All/repeat/mobile_element/Class_I/LTR/Ty1_copia/Ale`, lambda = 'ML', bounds = list(lambda = c(0.0005, 1))) #lambda branch transformation
summary(mod_precip_lambda_32)
mod_precip_lambda_33<-update(mod_precip_lambda_32,~. -`All/repeat/mobile_element/Class_I/pararetrovirus`, lambda = 'ML', bounds = list(lambda = c(0.0005, 1))) #lambda branch transformation
summary(mod_precip_lambda_33)
mod_precip_lambda_34<-update(mod_precip_lambda_33,~. -`All/repeat/mobile_element/Class_I/LTR/Ty1_copia/Angela`, lambda = 'ML', bounds = list(lambda = c(0.0005, 1))) #lambda branch transformation
summary(mod_precip_lambda_34)
mod_precip_lambda_35<-update(mod_precip_lambda_34,~. -`All/repeat/mobile_element/Class_I/LTR/Ty1_copia/Ikeros`, lambda = 'ML', bounds = list(lambda = c(0.0005, 1))) #lambda branch transformation
summary(mod_precip_lambda_35)
mod_precip_lambda_36<-update(mod_precip_lambda_35,~. -`All/repeat/mobile_element/Class_I/LTR/Ty1_copia/Bianca`, lambda = 'ML', bounds = list(lambda = c(0.0005, 1))) #lambda branch transformation
summary(mod_precip_lambda_36)
mod_precip_lambda_37<-update(mod_precip_lambda_36,~. -`All/repeat/mobile_element/Class_II/Subclass_1/TIR/hAT`, lambda = 'ML', bounds = list(lambda = c(0.0005, 1))) #lambda branch transformation
summary(mod_precip_lambda_37)
mod_precip_lambda_38<-update(mod_precip_lambda_37,~. -`All/repeat/mobile_element/Class_I/LTR/Ty3_gypsy/chromovirus/Tekay`, lambda = 'ML', bounds = list(lambda = c(0.0005, 1))) #lambda branch transformation
summary(mod_precip_lambda_38)
mod_precip_lambda_39<-update(mod_precip_lambda_38,~. -`All/repeat/mobile_element/Class_I/LTR/Ty3_gypsy/non-chromovirus/OTA/Tat/Ogre`, lambda = 'ML', bounds = list(lambda = c(0.0005, 1))) #lambda branch transformation
summary(mod_precip_lambda_39)
mod_precip_lambda_40<-update(mod_precip_lambda_39,~. -`All/repeat/mobile_element/Class_I/LTR/Ty3_gypsy/non-chromovirus/OTA/Tat/Retand`, lambda = 'ML', bounds = list(lambda = c(0.0005, 1))) #lambda branch transformation
summary(mod_precip_lambda_40)
mod_precip_lambda_41<-update(mod_precip_lambda_40,~. -`All/repeat/rDNA/45S_rDNA/18S_rDNA`, lambda = 'ML', bounds = list(lambda = c(0.0005, 1))) #lambda branch transformation
summary(mod_precip_lambda_41)
mod_precip_lambda_42<-update(mod_precip_lambda_41,~. -`All/repeat/mobile_element/Class_I/LINE`, lambda = 'ML', bounds = list(lambda = c(0.0005, 1))) #lambda branch transformation
summary(mod_precip_lambda_42)
mod_precip_lambda_43<-update(mod_precip_lambda_42,~. -`All/repeat/mobile_element/Class_I/LTR/Ty3_gypsy/chromovirus/CRM`, lambda = 'ML', bounds = list(lambda = c(0.0005, 1))) #lambda branch transformation
summary(mod_precip_lambda_43)
mod_precip_lambda_44<-update(mod_precip_lambda_43,~. -`All/repeat/mobile_element/Class_II/Subclass_1/TIR/PIF_Harbinger`, lambda = 'ML', bounds = list(lambda = c(0.0005, 1))) #lambda branch transformation
summary(mod_precip_lambda_44)
mod_precip_lambda_45<-update(mod_precip_lambda_44,~. -`All/repeat/satellite`, lambda = 'ML', bounds = list(lambda = c(0.0005, 1))) #lambda branch transformation
summary(mod_precip_lambda_45)
mod_precip_lambda_46<-update(mod_precip_lambda_45,~. -`All/repeat/mobile_element/Class_I/LTR/Ty1_copia/TAR`, lambda = 'ML', bounds = list(lambda = c(0.0005, 1))) #lambda branch transformation
summary(mod_precip_lambda_46)
mod_precip_lambda_47<-update(mod_precip_lambda_46,~. -`All/repeat/mobile_element/Class_I/LTR/Ty3_gypsy/non-chromovirus/OTA/Athila`, lambda = 'ML', bounds = list(lambda = c(0.0005, 1))) #lambda branch transformation
summary(mod_precip_lambda_47)
mod_precip_lambda_48<-update(mod_precip_lambda_47,~. -`All/repeat/mobile_element/Class_I/LTR/Ty1_copia/Ivana`, lambda = 'ML', bounds = list(lambda = c(0.0005, 1))) #lambda branch transformation
summary(mod_precip_lambda_48)
mod_precip_lambda_49<-update(mod_precip_lambda_48,~. -`All/repeat/rDNA/5S_rDNA`, lambda = 'ML', bounds = list(lambda = c(0.0005, 1))) #lambda branch transformation
summary(mod_precip_lambda_49)
mod_precip_lambda_50<-update(mod_precip_lambda_49,~. -`All/repeat/rDNA/45S_rDNA/25S_rDNA`, lambda = 'ML', bounds = list(lambda = c(0.0005, 1))) #lambda branch transformation
summary(mod_precip_lambda_50)
mod_precip_lambda_51<-update(mod_precip_lambda_50,~. -`All/repeat/mobile_element/Class_I/LTR/Ty1_copia/Angela`:as.factor(Genome_size_class), lambda = 'ML', bounds = list(lambda = c(0.0005, 1))) #lambda branch transformation
summary(mod_precip_lambda_51)


AIC(mod_precip_lambda,mod_precip_lambda_2,mod_precip_lambda_3,mod_precip_lambda_4,mod_precip_lambda_5,mod_precip_lambda_6,mod_precip_lambda_7,mod_precip_lambda_8,mod_precip_lambda_9,mod_precip_lambda_10,mod_precip_lambda_11,mod_precip_lambda_12,mod_precip_lambda_13,mod_precip_lambda_14,mod_precip_lambda_15,mod_precip_lambda_16,mod_precip_lambda_17,mod_precip_lambda_18,mod_precip_lambda_19,mod_precip_lambda_20,mod_precip_lambda_21,mod_precip_lambda_22,mod_precip_lambda_23,mod_precip_lambda_24,mod_precip_lambda_25,mod_precip_lambda_26,mod_precip_lambda_27,mod_precip_lambda_28,mod_precip_lambda_29,mod_precip_lambda_30,mod_precip_lambda_31,mod_precip_lambda_32,mod_precip_lambda_33,mod_precip_lambda_34,mod_precip_lambda_35,mod_precip_lambda_36,mod_precip_lambda_37,mod_precip_lambda_38,mod_precip_lambda_39,mod_precip_lambda_40,mod_precip_lambda_41,mod_precip_lambda_42,mod_precip_lambda_43,mod_precip_lambda_44,mod_precip_lambda_45,mod_precip_lambda_46,mod_precip_lambda_47,mod_precip_lambda_48,mod_precip_lambda_49,mod_precip_lambda_50,mod_precip_lambda_51)
summary(mod_precip_lambda_46)


#~~~~~Let's try the same with ABSOLUTE REPEAT NUMBERS/HITS~~~~~
ALL_DATA_palms_GS_vegan_abs<-read.csv("ALL_DATA_palms_GS_vegan_absol_vals.csv", header=T,row.names = 1)
head(ALL_DATA_palms_GS_vegan_abs)
ALL_DATA_palms_GS_vegan_abs$Genome_occupancy<-as.numeric(ALL_DATA_palms_GS_vegan_abs$Genome_occupancy)
mod_input_precip_abs<-ALL_DATA_palms_GS_vegan_abs[,c(1:30,32,49)]
colnames(mod_input_precip_abs)
mod_input_precip_abs_fin<-mod_input_precip_abs[,-c(1,2)]
mod_input_precip_abs_fin<-as.data.frame(apply(mod_input_precip_abs_fin, 2, as.numeric))
colnames(mod_input_precip_abs_fin)
dim(mod_input_precip_abs_fin)

mod_input_precip_abs_unique<-mod_input_precip_abs[-which(duplicated(mod_input_precip_abs$Tree_name)),-2]
mod_input_precip_abs_unique[,-1]<-as.data.frame(apply(mod_input_precip_abs_unique[,-1], 2, as.numeric))
head(mod_input_precip_abs_unique)

#Let's create genome size bins
sum_GS<-summary(mod_input_precip_abs_unique$Genome_size)

q1_GS<-which(mod_input_precip_abs_unique$Genome_size <= sum_GS[[3]])
q2_GS<-which(mod_input_precip_abs_unique$Genome_size > sum_GS[[3]])

mod_input_precip_abs_unique[q1_GS,(ncol(mod_input_precip_abs_unique)+1)]<-"Small"
mod_input_precip_abs_unique[q2_GS,(ncol(mod_input_precip_abs_unique))]<-"Large"
colnames(mod_input_precip_abs_unique)[ncol(mod_input_precip_abs_unique)]<-"Genome_size_class"
head(mod_input_precip_abs_unique)

colnames(mod_input_precip_abs_unique)
dim(mod_input_precip_abs_unique)
pgls_dat_precip_abs<-comparative.data(data=mod_input_precip_abs_unique, phy=phyclean, names.col=Tree_name,vcv=TRUE)
print(pgls_dat_precip_abs)

#remove variables with zero variance, and remember that R takes interactions alphabetically, so the first variables with no interactions are for 'large' genome species
mod_precip_lambda_abs<-pgls(sqrt(as.numeric(Precipitation_of_Driest_Month))~All.repeat.mobile_element.Class_I.LINE+All.repeat.mobile_element.Class_I.LTR.Ty1_copia.Ale+All.repeat.mobile_element.Class_I.LTR.Ty1_copia.Alesia+All.repeat.mobile_element.Class_I.LTR.Ty1_copia.Angela+All.repeat.mobile_element.Class_I.LTR.Ty1_copia.Bianca+All.repeat.mobile_element.Class_I.LTR.Ty1_copia.Ikeros+All.repeat.mobile_element.Class_I.LTR.Ty1_copia.Ivana+All.repeat.mobile_element.Class_I.LTR.Ty1_copia.SIRE+All.repeat.mobile_element.Class_I.LTR.Ty1_copia.TAR+All.repeat.mobile_element.Class_I.LTR.Ty1_copia.Tork+All.repeat.mobile_element.Class_I.LTR.Ty3_gypsy.chromovirus.CRM+All.repeat.mobile_element.Class_I.LTR.Ty3_gypsy.chromovirus.Galadriel+All.repeat.mobile_element.Class_I.LTR.Ty3_gypsy.chromovirus.Reina+All.repeat.mobile_element.Class_I.LTR.Ty3_gypsy.chromovirus.Tekay+All.repeat.mobile_element.Class_I.LTR.Ty3_gypsy.non_chromovirus.OTA.Athila+All.repeat.mobile_element.Class_I.LTR.Ty3_gypsy.non_chromovirus.OTA.Tat.Ogre+All.repeat.mobile_element.Class_I.LTR.Ty3_gypsy.non_chromovirus.OTA.Tat.Retand+All.repeat.mobile_element.Class_I.pararetrovirus+All.repeat.mobile_element.Class_II.Subclass_1.TIR.EnSpm_CACTA+All.repeat.mobile_element.Class_II.Subclass_1.TIR.hAT+All.repeat.mobile_element.Class_II.Subclass_1.TIR.MuDR_Mutator+All.repeat.mobile_element.Class_II.Subclass_1.TIR.PIF_Harbinger+All.repeat.rDNA.45S_rDNA.18S_rDNA+All.repeat.rDNA.45S_rDNA.25S_rDNA+All.repeat.rDNA.5S_rDNA+All.repeat.satellite+as.factor(Genome_size_class):(All.repeat.mobile_element.Class_I.LINE+All.repeat.mobile_element.Class_I.LTR.Ty1_copia.Ale+All.repeat.mobile_element.Class_I.LTR.Ty1_copia.Alesia+All.repeat.mobile_element.Class_I.LTR.Ty1_copia.Angela+All.repeat.mobile_element.Class_I.LTR.Ty1_copia.Bianca+All.repeat.mobile_element.Class_I.LTR.Ty1_copia.Ikeros+All.repeat.mobile_element.Class_I.LTR.Ty1_copia.Ivana+All.repeat.mobile_element.Class_I.LTR.Ty1_copia.SIRE+All.repeat.mobile_element.Class_I.LTR.Ty1_copia.TAR+All.repeat.mobile_element.Class_I.LTR.Ty1_copia.Tork+All.repeat.mobile_element.Class_I.LTR.Ty3_gypsy.chromovirus.CRM+All.repeat.mobile_element.Class_I.LTR.Ty3_gypsy.chromovirus.Galadriel+All.repeat.mobile_element.Class_I.LTR.Ty3_gypsy.chromovirus.Reina+All.repeat.mobile_element.Class_I.LTR.Ty3_gypsy.chromovirus.Tekay+All.repeat.mobile_element.Class_I.LTR.Ty3_gypsy.non_chromovirus.OTA.Athila+All.repeat.mobile_element.Class_I.LTR.Ty3_gypsy.non_chromovirus.OTA.Tat.Ogre+All.repeat.mobile_element.Class_I.LTR.Ty3_gypsy.non_chromovirus.OTA.Tat.Retand+All.repeat.mobile_element.Class_I.pararetrovirus+All.repeat.mobile_element.Class_II.Subclass_1.TIR.EnSpm_CACTA+All.repeat.mobile_element.Class_II.Subclass_1.TIR.hAT+All.repeat.mobile_element.Class_II.Subclass_1.TIR.MuDR_Mutator+All.repeat.mobile_element.Class_II.Subclass_1.TIR.PIF_Harbinger+All.repeat.rDNA.45S_rDNA.18S_rDNA+All.repeat.rDNA.45S_rDNA.25S_rDNA+All.repeat.rDNA.5S_rDNA+All.repeat.satellite), data=pgls_dat_precip_abs, lambda = 'ML', bounds = list(lambda = c(0.0005, 1))) #lambda branch transformation
summary(mod_precip_lambda_abs)

mod_precip_lambda_abs_2<-update(mod_precip_lambda_abs,~. -All.repeat.mobile_element.Class_I.LTR.Ty1_copia.Bianca:as.factor(Genome_size_class), lambda = 'ML', bounds = list(lambda = c(0.0005, 1))) #lambda branch transformation
summary(mod_precip_lambda_abs_2)
mod_precip_lambda_abs_3<-update(mod_precip_lambda_abs_2,~. -All.repeat.satellite:as.factor(Genome_size_class), lambda = 'ML', bounds = list(lambda = c(0.0005, 1))) #lambda branch transformation
summary(mod_precip_lambda_abs_3)
mod_precip_lambda_abs_4<-update(mod_precip_lambda_abs_3,~. -All.repeat.mobile_element.Class_I.LTR.Ty3_gypsy.chromovirus.CRM:as.factor(Genome_size_class), lambda = 'ML', bounds = list(lambda = c(0.0005, 1))) #lambda branch transformation
summary(mod_precip_lambda_abs_4)
mod_precip_lambda_abs_5<-update(mod_precip_lambda_abs_4,~. -All.repeat.mobile_element.Class_I.LTR.Ty1_copia.TAR:as.factor(Genome_size_class), lambda = 'ML', bounds = list(lambda = c(0.0005, 1))) #lambda branch transformation
summary(mod_precip_lambda_abs_5)
mod_precip_lambda_abs_6<-update(mod_precip_lambda_abs_5,~. -All.repeat.mobile_element.Class_II.Subclass_1.TIR.MuDR_Mutator:as.factor(Genome_size_class), lambda = 'ML', bounds = list(lambda = c(0.0005, 1))) #lambda branch transformation
summary(mod_precip_lambda_abs_6)
mod_precip_lambda_abs_7<-update(mod_precip_lambda_abs_6,~. -All.repeat.mobile_element.Class_I.LTR.Ty1_copia.Alesia:as.factor(Genome_size_class), lambda = 'ML', bounds = list(lambda = c(0.0005, 1))) #lambda branch transformation
summary(mod_precip_lambda_abs_7)
mod_precip_lambda_abs_8<-update(mod_precip_lambda_abs_7,~. -All.repeat.rDNA.45S_rDNA.25S_rDNA:as.factor(Genome_size_class), lambda = 'ML', bounds = list(lambda = c(0.0005, 1))) #lambda branch transformation
summary(mod_precip_lambda_abs_8)
mod_precip_lambda_abs_9<-update(mod_precip_lambda_abs_8,~. -All.repeat.mobile_element.Class_I.LTR.Ty3_gypsy.non_chromovirus.OTA.Tat.Retand:as.factor(Genome_size_class), lambda = 'ML', bounds = list(lambda = c(0.0005, 1))) #lambda branch transformation
summary(mod_precip_lambda_abs_9)
mod_precip_lambda_abs_10<-update(mod_precip_lambda_abs_9,~. -All.repeat.mobile_element.Class_I.LTR.Ty3_gypsy.chromovirus.Reina:as.factor(Genome_size_class), lambda = 'ML', bounds = list(lambda = c(0.0005, 1))) #lambda branch transformation
summary(mod_precip_lambda_abs_10)
mod_precip_lambda_abs_11<-update(mod_precip_lambda_abs_10,~. -All.repeat.mobile_element.Class_I.LTR.Ty3_gypsy.non_chromovirus.OTA.Athila:as.factor(Genome_size_class), lambda = 'ML', bounds = list(lambda = c(0.0005, 1))) #lambda branch transformation
summary(mod_precip_lambda_abs_11)
mod_precip_lambda_abs_12<-update(mod_precip_lambda_abs_11,~. -All.repeat.mobile_element.Class_II.Subclass_1.TIR.PIF_Harbinger:as.factor(Genome_size_class), lambda = 'ML', bounds = list(lambda = c(0.0005, 1))) #lambda branch transformation
summary(mod_precip_lambda_abs_12)
mod_precip_lambda_abs_13<-update(mod_precip_lambda_abs_12,~. -All.repeat.mobile_element.Class_II.Subclass_1.TIR.hAT:as.factor(Genome_size_class), lambda = 'ML', bounds = list(lambda = c(0.0005, 1))) #lambda branch transformation
summary(mod_precip_lambda_abs_13)
mod_precip_lambda_abs_14<-update(mod_precip_lambda_abs_13,~. -All.repeat.mobile_element.Class_I.LTR.Ty1_copia.Ale:as.factor(Genome_size_class), lambda = 'ML', bounds = list(lambda = c(0.0005, 1))) #lambda branch transformation
summary(mod_precip_lambda_abs_14)
mod_precip_lambda_abs_15<-update(mod_precip_lambda_abs_14,~. -All.repeat.mobile_element.Class_I.LTR.Ty1_copia.Ivana:as.factor(Genome_size_class), lambda = 'ML', bounds = list(lambda = c(0.0005, 1))) #lambda branch transformation
summary(mod_precip_lambda_abs_15)
mod_precip_lambda_abs_16<-update(mod_precip_lambda_abs_15,~. -All.repeat.mobile_element.Class_I.pararetrovirus:as.factor(Genome_size_class), lambda = 'ML', bounds = list(lambda = c(0.0005, 1))) #lambda branch transformation
summary(mod_precip_lambda_abs_16)
mod_precip_lambda_abs_17<-update(mod_precip_lambda_abs_16,~. -All.repeat.mobile_element.Class_I.LTR.Ty1_copia.Tork:as.factor(Genome_size_class), lambda = 'ML', bounds = list(lambda = c(0.0005, 1))) #lambda branch transformation
summary(mod_precip_lambda_abs_17)
mod_precip_lambda_abs_18<-update(mod_precip_lambda_abs_17,~. -All.repeat.mobile_element.Class_I.LTR.Ty3_gypsy.non_chromovirus.OTA.Tat.Ogre:as.factor(Genome_size_class), lambda = 'ML', bounds = list(lambda = c(0.0005, 1))) #lambda branch transformation
summary(mod_precip_lambda_abs_18)
mod_precip_lambda_abs_19<-update(mod_precip_lambda_abs_18,~. -All.repeat.mobile_element.Class_I.LTR.Ty3_gypsy.chromovirus.Tekay:as.factor(Genome_size_class), lambda = 'ML', bounds = list(lambda = c(0.0005, 1))) #lambda branch transformation
summary(mod_precip_lambda_abs_19)
mod_precip_lambda_abs_20<-update(mod_precip_lambda_abs_19,~. -All.repeat.mobile_element.Class_I.LINE:as.factor(Genome_size_class), lambda = 'ML', bounds = list(lambda = c(0.0005, 1))) #lambda branch transformation
summary(mod_precip_lambda_abs_20)
mod_precip_lambda_abs_21<-update(mod_precip_lambda_abs_20,~. -All.repeat.mobile_element.Class_I.LTR.Ty1_copia.SIRE:as.factor(Genome_size_class), lambda = 'ML', bounds = list(lambda = c(0.0005, 1))) #lambda branch transformation
summary(mod_precip_lambda_abs_21)
mod_precip_lambda_abs_22<-update(mod_precip_lambda_abs_21,~. -All.repeat.rDNA.45S_rDNA.18S_rDNA:as.factor(Genome_size_class), lambda = 'ML', bounds = list(lambda = c(0.0005, 1))) #lambda branch transformation
summary(mod_precip_lambda_abs_22)
mod_precip_lambda_abs_23<-update(mod_precip_lambda_abs_22,~. -All.repeat.mobile_element.Class_I.LTR.Ty1_copia.Ikeros:as.factor(Genome_size_class), lambda = 'ML', bounds = list(lambda = c(0.0005, 1))) #lambda branch transformation
summary(mod_precip_lambda_abs_23)
mod_precip_lambda_abs_24<-update(mod_precip_lambda_abs_23,~. -All.repeat.mobile_element.Class_II.Subclass_1.TIR.EnSpm_CACTA:as.factor(Genome_size_class), lambda = 'ML', bounds = list(lambda = c(0.0005, 1))) #lambda branch transformation
summary(mod_precip_lambda_abs_24)
mod_precip_lambda_abs_25<-update(mod_precip_lambda_abs_24,~. -All.repeat.mobile_element.Class_I.LTR.Ty3_gypsy.chromovirus.Galadriel:as.factor(Genome_size_class), lambda = 'ML', bounds = list(lambda = c(0.0005, 1))) #lambda branch transformation
summary(mod_precip_lambda_abs_25)
mod_precip_lambda_abs_26<-update(mod_precip_lambda_abs_25,~. -All.repeat.rDNA.5S_rDNA:as.factor(Genome_size_class), lambda = 'ML', bounds = list(lambda = c(0.0005, 1))) #lambda branch transformation
summary(mod_precip_lambda_abs_26)
mod_precip_lambda_abs_27<-update(mod_precip_lambda_abs_26,~. -All.repeat.mobile_element.Class_II.Subclass_1.TIR.MuDR_Mutator, lambda = 'ML', bounds = list(lambda = c(0.0005, 1))) #lambda branch transformation
summary(mod_precip_lambda_abs_27)
mod_precip_lambda_abs_28<-update(mod_precip_lambda_abs_27,~. -All.repeat.mobile_element.Class_I.LTR.Ty1_copia.Alesia, lambda = 'ML', bounds = list(lambda = c(0.0005, 1))) #lambda branch transformation
summary(mod_precip_lambda_abs_28)
mod_precip_lambda_abs_29<-update(mod_precip_lambda_abs_28,~. -All.repeat.mobile_element.Class_I.LTR.Ty3_gypsy.chromovirus.Galadriel, lambda = 'ML', bounds = list(lambda = c(0.0005, 1))) #lambda branch transformation
summary(mod_precip_lambda_abs_29)
mod_precip_lambda_abs_30<-update(mod_precip_lambda_abs_29,~. -All.repeat.mobile_element.Class_I.LTR.Ty3_gypsy.chromovirus.Reina, lambda = 'ML', bounds = list(lambda = c(0.0005, 1))) #lambda branch transformation
summary(mod_precip_lambda_abs_30)
mod_precip_lambda_abs_31<-update(mod_precip_lambda_abs_30,~. -All.repeat.mobile_element.Class_I.LTR.Ty1_copia.SIRE, lambda = 'ML', bounds = list(lambda = c(0.0005, 1))) #lambda branch transformation
summary(mod_precip_lambda_abs_31)
mod_precip_lambda_abs_32<-update(mod_precip_lambda_abs_31,~. -All.repeat.mobile_element.Class_I.LTR.Ty1_copia.Ale, lambda = 'ML', bounds = list(lambda = c(0.0005, 1))) #lambda branch transformation
summary(mod_precip_lambda_abs_32)
mod_precip_lambda_abs_33<-update(mod_precip_lambda_abs_32,~. -All.repeat.mobile_element.Class_I.pararetrovirus, lambda = 'ML', bounds = list(lambda = c(0.0005, 1))) #lambda branch transformation
summary(mod_precip_lambda_abs_33)
mod_precip_lambda_abs_34<-update(mod_precip_lambda_abs_33,~. -All.repeat.mobile_element.Class_I.LTR.Ty1_copia.Ikeros , lambda = 'ML', bounds = list(lambda = c(0.0005, 1))) #lambda branch transformation
summary(mod_precip_lambda_abs_34)
mod_precip_lambda_abs_35<-update(mod_precip_lambda_abs_34,~. -All.repeat.mobile_element.Class_I.LTR.Ty1_copia.Angela, lambda = 'ML', bounds = list(lambda = c(0.0005, 1))) #lambda branch transformation
summary(mod_precip_lambda_abs_35)
mod_precip_lambda_abs_36<-update(mod_precip_lambda_abs_35,~. -All.repeat.mobile_element.Class_I.LTR.Ty1_copia.Bianca, lambda = 'ML', bounds = list(lambda = c(0.0005, 1))) #lambda branch transformation
summary(mod_precip_lambda_abs_36)
mod_precip_lambda_abs_37<-update(mod_precip_lambda_abs_36,~. -All.repeat.mobile_element.Class_II.Subclass_1.TIR.hAT, lambda = 'ML', bounds = list(lambda = c(0.0005, 1))) #lambda branch transformation
summary(mod_precip_lambda_abs_37)
mod_precip_lambda_abs_38<-update(mod_precip_lambda_abs_37,~. -All.repeat.mobile_element.Class_I.LTR.Ty3_gypsy.chromovirus.Tekay, lambda = 'ML', bounds = list(lambda = c(0.0005, 1))) #lambda branch transformation
summary(mod_precip_lambda_abs_38)
mod_precip_lambda_abs_39<-update(mod_precip_lambda_abs_38,~. -All.repeat.mobile_element.Class_I.LTR.Ty3_gypsy.non_chromovirus.OTA.Tat.Ogre, lambda = 'ML', bounds = list(lambda = c(0.0005, 1))) #lambda branch transformation
summary(mod_precip_lambda_abs_39)
mod_precip_lambda_abs_40<-update(mod_precip_lambda_abs_39,~. -All.repeat.mobile_element.Class_I.LTR.Ty3_gypsy.non_chromovirus.OTA.Tat.Retand, lambda = 'ML', bounds = list(lambda = c(0.0005, 1))) #lambda branch transformation
summary(mod_precip_lambda_abs_40)
mod_precip_lambda_abs_41<-update(mod_precip_lambda_abs_40,~. -All.repeat.rDNA.45S_rDNA.18S_rDNA, lambda = 'ML', bounds = list(lambda = c(0.0005, 1))) #lambda branch transformation
summary(mod_precip_lambda_abs_41)
mod_precip_lambda_abs_42<-update(mod_precip_lambda_abs_41,~. -All.repeat.mobile_element.Class_I.LINE, lambda = 'ML', bounds = list(lambda = c(0.0005, 1))) #lambda branch transformation
summary(mod_precip_lambda_abs_42)
mod_precip_lambda_abs_43<-update(mod_precip_lambda_abs_42,~. -All.repeat.mobile_element.Class_I.LTR.Ty3_gypsy.chromovirus.CRM, lambda = 'ML', bounds = list(lambda = c(0.0005, 1))) #lambda branch transformation
summary(mod_precip_lambda_abs_43)
mod_precip_lambda_abs_44<-update(mod_precip_lambda_abs_43,~. -All.repeat.mobile_element.Class_II.Subclass_1.TIR.PIF_Harbinger, lambda = 'ML', bounds = list(lambda = c(0.0005, 1))) #lambda branch transformation
summary(mod_precip_lambda_abs_44)
mod_precip_lambda_abs_45<-update(mod_precip_lambda_abs_44,~. -All.repeat.mobile_element.Class_I.LTR.Ty1_copia.TAR, lambda = 'ML', bounds = list(lambda = c(0.0005, 1))) #lambda branch transformation
summary(mod_precip_lambda_abs_45)
mod_precip_lambda_abs_46<-update(mod_precip_lambda_abs_45,~. -All.repeat.mobile_element.Class_I.LTR.Ty3_gypsy.non_chromovirus.OTA.Athila, lambda = 'ML', bounds = list(lambda = c(0.0005, 1))) #lambda branch transformation
summary(mod_precip_lambda_abs_46)
mod_precip_lambda_abs_47<-update(mod_precip_lambda_abs_46,~. -All.repeat.satellite, lambda = 'ML', bounds = list(lambda = c(0.0005, 1))) #lambda branch transformation
summary(mod_precip_lambda_abs_47)
mod_precip_lambda_abs_48<-update(mod_precip_lambda_abs_47,~. -All.repeat.mobile_element.Class_I.LTR.Ty1_copia.Ivana, lambda = 'ML', bounds = list(lambda = c(0.0005, 1))) #lambda branch transformation
summary(mod_precip_lambda_abs_48)
mod_precip_lambda_abs_49<-update(mod_precip_lambda_abs_48,~. -All.repeat.rDNA.5S_rDNA, lambda = 'ML', bounds = list(lambda = c(0.0005, 1))) #lambda branch transformation
summary(mod_precip_lambda_abs_49)
mod_precip_lambda_abs_50<-update(mod_precip_lambda_abs_49,~. -All.repeat.rDNA.45S_rDNA.25S_rDNA, lambda = 'ML', bounds = list(lambda = c(0.0005, 1))) #lambda branch transformation
summary(mod_precip_lambda_abs_50)


AIC(mod_precip_lambda_abs,mod_precip_lambda_abs_2,mod_precip_lambda_abs_3,mod_precip_lambda_abs_4,mod_precip_lambda_abs_5,mod_precip_lambda_abs_6,mod_precip_lambda_abs_7,mod_precip_lambda_abs_8,mod_precip_lambda_abs_9,mod_precip_lambda_abs_10,mod_precip_lambda_abs_11,mod_precip_lambda_abs_12,mod_precip_lambda_abs_13,mod_precip_lambda_abs_14,mod_precip_lambda_abs_15,mod_precip_lambda_abs_16,mod_precip_lambda_abs_17,mod_precip_lambda_abs_18,mod_precip_lambda_abs_19,mod_precip_lambda_abs_20,mod_precip_lambda_abs_21,mod_precip_lambda_abs_22,mod_precip_lambda_abs_23,mod_precip_lambda_abs_24,mod_precip_lambda_abs_25,mod_precip_lambda_abs_26,mod_precip_lambda_abs_27,mod_precip_lambda_abs_28,mod_precip_lambda_abs_29,mod_precip_lambda_abs_30,mod_precip_lambda_abs_31,mod_precip_lambda_abs_32,mod_precip_lambda_abs_33,mod_precip_lambda_abs_34,mod_precip_lambda_abs_35,mod_precip_lambda_abs_36,mod_precip_lambda_abs_37,mod_precip_lambda_abs_38,mod_precip_lambda_abs_39,mod_precip_lambda_abs_40,mod_precip_lambda_abs_41,mod_precip_lambda_abs_42,mod_precip_lambda_abs_43,mod_precip_lambda_abs_44,mod_precip_lambda_abs_45,mod_precip_lambda_abs_46,mod_precip_lambda_abs_47,mod_precip_lambda_abs_48,mod_precip_lambda_abs_49,mod_precip_lambda_abs_50)
summary(mod_precip_lambda_abs_45)
#Same result as percent repeat in each genome


### Let's plot a matrix of genome proportions for each repeat with a phylogenetic tree showing precip. driest month
#Get data to match tree and vice versa
fig_input_precip<-mod_input_precip
colnames(fig_input_precip)
fig_input_precip_fin<-fig_input_precip
colnames(fig_input_precip_fin)
dim(fig_input_precip_fin)

fig_tree_names<-fig_input_precip_fin[-which(duplicated(fig_input_precip_fin$Tree_name)),1]
length(fig_tree_names)

fig_tree_names_fin<-fig_tree_names[which(fig_tree_names%in%phyclean$tip.label)]
length(fig_tree_names_fin)

phyclean_fig<-keep.tip(phyclean, fig_tree_names_fin)
plot(phyclean_fig)


fig_input_precip_fin<-fig_input_precip_fin[which(fig_input_precip_fin$Tree_name%in%phyclean_fig$tip.label),]
fig_input_precip_fin<-fig_input_precip_fin[-which(duplicated(fig_input_precip_fin$Tree_name)),]
dim(fig_input_precip_fin)

precip_data<-fig_input_precip_fin[,c(1,32)]
precip_data<-column_to_rownames(precip_data, loc=1)
precip_data<-as.matrix(precip_data)[,1]  #### this it is KEY for phytools!
head(precip_data)
length(precip_data)


name.check(fig_input_precip_fin, phyclean_fig)


#Do and plot contmap of precip driest month
layout(matrix(1:2,1,2),widths=c(0.4,0.6))
par(cex=1)
precip_contmap<-contMap(phyclean_fig,precip_data,sig=1, fsize= c(0.4, 1))
precip_contmap

#Do and plot corrplot of repeat profiles
repeat_profiles<-fig_input_precip_fin[,c(1,3:30)]
rn<-repeat_profiles[,1]
repeat_profiles<-repeat_profiles[,-1]
repeat_profiles<-apply(repeat_profiles, 2, as.numeric)
rownames(repeat_profiles)<-rn
head(repeat_profiles)
repeat_profiles2<-repeat_profiles[match(phyclean_fig$tip.label, rownames(repeat_profiles)),]
head(repeat_profiles2)
head(repeat_profiles2)
dim(repeat_profiles2)

phylo.heatmap(phyclean_fig, repeat_profiles2, method= , fsize= c(0.4,0.4, 1))
### now use inkscape to stitch the former two plots together


###Let's plot repeat types as stacked bars on a tree in ggtree
p <- ggtree(phyclean_fig) + geom_tiplab(size=5)
p
rp2m<-melt(repeat_profiles2)

p2 <- facet_plot(p, panel='Repeat proportion of genome', data=rp2m, 
                 geom = geom_barh, mapping = aes(x = value, fill = as.factor(Var2)), stat='identity', show.legend=T)

p2+theme_tree2()


###Let's plot repeat richness, occupancy and diversity as bars on a tree in ggtree
#Make data frame
ecol_indices<-ALL_DATA_palms_GS_vegan_pct[,c(1,32:35)]
en<-ecol_indices[,1]
ecol_indices<-ecol_indices[,-1]
ecol_indices<-apply(ecol_indices, 2, as.numeric)
rownames(ecol_indices)<-en
head(ecol_indices)
ecol_indices2<-ecol_indices[match(phyclean_fig$tip.label, rownames(ecol_indices)),]
head(repeat_profiles2)
dim(ecol_indices)
dim(ecol_indices2)


#Do the plot
p <- ggtree(phyclean_fig) + geom_tiplab(size=1)
p
eifin<-melt(ecol_indices2)

genome_size<-eifin[which(eifin$Var2=="Genome_size"),]
richness<-eifin[which(eifin$Var2=="Mehinicks_index"),]
occupancy<-eifin[which(eifin$Var2=="Genome_occupancy"),]
diversity<-eifin[which(eifin$Var2=="Shannon_Index"),]

p2 <- facet_plot(p, panel="Genome size", data=genome_size, 
                 geom = geom_barh, mapping = aes(x = value, fill="4"), stat='identity', show.legend=F)

p3 <- facet_plot(p2, panel="Repeat richness (Mehinicks's index)", data=richness, 
                 geom = geom_barh, mapping = aes(x = value, fill="1"), stat='identity', show.legend=F)

p4 <- facet_plot(p3, panel='% of the genome occupied by repeats', data=occupancy, 
                 geom = geom_barh, mapping = aes(x = value, fill = "2"), stat='identity', show.legend=F)

p5 <- facet_plot(p4, panel='Repeat diversity (Shannon Index)', data=diversity, 
                 geom = geom_barh, mapping = aes(x = value, fill="3"), stat='identity', show.legend=F)

p5+theme_tree2()


 ##~~~~~~~~~~~~~~~~~~~~PGLS - WORLDCLIM VARIABLES AND ECOLOGICAL METRICS~~~~~~~~~~~~~~~~~~~~## 
######### LET's SEE WHICH ENVIRONMENTAL VARIABLES AFFECT REPEAT RICHNESS, GENOME OCCUPANCY AND DIVERSITY ########

##### REPEAT TYPE RICHNESS (Mehinicks Index)######
#Test with simple lm
mod_input_MI<-ALL_DATA_palms_GS_vegan_pct[,c(1,2,34,36:54)]
mod_input_MI_fin<-mod_input_MI[,-c(1,2)]
colnames(mod_input_MI_fin)
dim(mod_input_MI_fin)

#How to normalise richness (response variable)
shapiro.test(mod_input_MI_fin$Mehinicks_index)
hist(mod_input_MI_fin$Mehinicks_index)
shapiro.test(log(mod_input_MI_fin$Mehinicks_index))

mod_1_MI<-lm(log(as.numeric(Mehinicks_index))~., data=mod_input_MI_fin)
summary(mod_1_MI)


#Which variables should we keep?
alias(mod_1_MI) #Max_Temperature_of_Warmest_Month is correlated

mod_1_MI_upd<-update(mod_1_MI,~.-Max_Temperature_of_Warmest_Month, data=mod_input_MI_fin)
summary(mod_1_MI_upd)
alias(mod_1_MI_upd)
vif(mod_1_MI_upd)

#Worth removing:
#Annual_Mean_Temperature	
#Temperature_Seasonality_standard_deviation_100
#Mean_Temperature_of_Warmest_Quarter	
#Mean_Temperature_of_Coldest_Quarter
#Precipitation_of_Wettest_Quarter	
#Precipitation_of_Driest_Quarter
#Precipitation_of_Warmest_Quarter	
#Precipitation_of_Coldest_Quarter


mod_2_MI_upd<-update(mod_1_MI_upd,~.-Annual_Mean_Temperature-Temperature_Seasonality_standard_deviation_100-Mean_Temperature_of_Warmest_Quarter-Mean_Temperature_of_Coldest_Quarter-Precipitation_of_Wettest_Quarter-Precipitation_of_Driest_Quarter-Precipitation_of_Warmest_Quarter-Precipitation_of_Coldest_Quarter, data=mod_input_MI_fin)
summary(mod_2_MI_upd)
alias(mod_2_MI_upd)
vif(mod_2_MI_upd)

#Worth removing:
#Annual_Precipitation
#Mean_Diurnal_Range_Mean_of_monthly_max_temp_min_temp
#Min_Temperature_of_Coldest_Month


mod_3_MI_upd<-update(mod_2_MI_upd,~.-Annual_Precipitation-Min_Temperature_of_Coldest_Month-Mean_Diurnal_Range_Mean_of_monthly_max_temp_min_temp, data=mod_input_MI_fin)
summary(mod_3_MI_upd)
alias(mod_3_MI_upd)
vif(mod_3_MI_upd)


mod_2_MI<-step(mod_3_MI_upd)
summary(mod_2_MI) #Cool, seems interesting.


#Let's try in a phylogenetically-corrected fashion
#PGLS of repeat richness vs worldclim variables, keeping only variables listed above
mod_input_MI_unique<-mod_input_MI[-which(duplicated(mod_input_MI$Tree_name)),c(1,3,10,11,12,16,17,18)]
colnames(mod_input_MI_unique)
dim(mod_input_MI_unique)

pairs(mod_input_MI_unique[,-1])

pgls_dat_MI<-comparative.data(data=mod_input_MI_unique, phy=phyclean, names.col="Tree_name",vcv=TRUE)
print(pgls_dat_MI)

mod_MI_lambda<-pgls(log(as.numeric(Mehinicks_index))~Temperature_Annual_Range_BIO5_BIO6+Mean_Temperature_of_Driest_Quarter+Precipitation_of_Driest_Month+Mean_Temperature_of_Wettest_Quarter+Precipitation_of_Wettest_Month+Precipitation_Seasonality_Coefficient_of_Variation, data=pgls_dat_MI, lambda = 'ML', bounds = list(lambda = c(0.0005, 1))) #lambda branch transformation
summary(mod_MI_lambda)

mod_MI_lambda_2<-update(mod_MI_lambda,~. -Precipitation_Seasonality_Coefficient_of_Variation, lambda = 'ML', bounds = list(lambda = c(0.0005, 1))) #lambda branch transformation
summary(mod_MI_lambda_2)
mod_MI_lambda_3<-update(mod_MI_lambda_2,~. -Temperature_Annual_Range_BIO5_BIO6, lambda = 'ML', bounds = list(lambda = c(0.0005, 1))) #lambda branch transformation
summary(mod_MI_lambda_3)
mod_MI_lambda_4<-update(mod_MI_lambda_3,~. -Precipitation_of_Driest_Month, lambda = 'ML', bounds = list(lambda = c(0.0005, 1))) #lambda branch transformation
summary(mod_MI_lambda_4)
mod_MI_lambda_5<-update(mod_MI_lambda_4,~. -Mean_Temperature_of_Driest_Quarter, lambda = 'ML', bounds = list(lambda = c(0.0005, 1))) #lambda branch transformation
summary(mod_MI_lambda_5)
mod_MI_lambda_6<-update(mod_MI_lambda_5,~. -Precipitation_of_Wettest_Month, lambda = 'ML', bounds = list(lambda = c(0.0005, 1))) #lambda branch transformation
summary(mod_MI_lambda_6)
#mod_MI_lambda_7<-update(mod_MI_lambda_6,~. -Mean_Temperature_of_Wettest_Quarter, lambda = 'ML', bounds = list(lambda = c(0.0005, 1))) #lambda branch transformation
#summary(mod_MI_lambda_7)

#Mean temp of wettest quarter is significant


##### REPEAT GENOME OCCUPANCY ######
#Test with simple lm
mod_input_occ<-ALL_DATA_palms_GS_vegan_pct[,c(1,2,35,36:54)]
mod_input_occ_fin<-mod_input_occ[,-c(1,2)]
colnames(mod_input_occ_fin)
dim(mod_input_occ_fin)


#How to normalise richness (response variable)
shapiro.test(mod_input_occ_fin$Genome_occupancy)
hist(mod_input_occ_fin$Genome_occupancy) # seems fine untransformed


mod_1_occ<-lm(Genome_occupancy~., data=mod_input_occ_fin)
summary(mod_1_occ)

mod_2_occ<-step(mod_1_occ)
summary(mod_2_occ) #Cool, seems interesting.


#Let's try in a phylogenetically-corrected fashion
#PGLS of genome size vs repeat class
mod_input_occ_unique<-mod_input_occ[-which(duplicated(mod_input_occ$Tree_name)),c(1,3,6,10,11,12,16,17,18)]
colnames(mod_input_occ_unique)
dim(mod_input_occ_unique)

pairs(mod_input_occ_unique[,-1])

pgls_dat_occ<-comparative.data(data=mod_input_occ_unique, phy=phyclean, names.col="Tree_name",vcv=TRUE)
print(pgls_dat_occ)

mod_occ_lambda<-pgls(Genome_occupancy~Temperature_Annual_Range_BIO5_BIO6+Mean_Temperature_of_Driest_Quarter+Precipitation_of_Driest_Month+Mean_Temperature_of_Wettest_Quarter+Precipitation_of_Wettest_Month+Precipitation_Seasonality_Coefficient_of_Variation, data=pgls_dat_occ, lambda = 'ML', bounds = list(lambda = c(0.0005, 1))) #lambda branch transformation
summary(mod_occ_lambda)

mod_occ_lambda_2<-update(mod_occ_lambda,~. -Precipitation_Seasonality_Coefficient_of_Variation, lambda = 'ML', bounds = list(lambda = c(0.0005, 1))) #lambda branch transformation
summary(mod_occ_lambda_2)
mod_occ_lambda_3<-update(mod_occ_lambda_2,~. -Mean_Temperature_of_Wettest_Quarter, lambda = 'ML', bounds = list(lambda = c(0.0005, 1))) #lambda branch transformation
summary(mod_occ_lambda_3)
mod_occ_lambda_4<-update(mod_occ_lambda_3,~. -Mean_Temperature_of_Driest_Quarter, lambda = 'ML', bounds = list(lambda = c(0.0005, 1))) #lambda branch transformation
summary(mod_occ_lambda_4)
mod_occ_lambda_5<-update(mod_occ_lambda_4,~. -Precipitation_of_Driest_Month, lambda = 'ML', bounds = list(lambda = c(0.0005, 1))) #lambda branch transformation
summary(mod_occ_lambda_5)
mod_occ_lambda_6<-update(mod_occ_lambda_5,~. -Precipitation_of_Wettest_Month, lambda = 'ML', bounds = list(lambda = c(0.009, 1))) #lambda branch transformation
summary(mod_occ_lambda_6)
#Temp annual range is significant 

AIC(mod_occ_lambda_6,mod_occ_lambda_5,mod_occ_lambda_4,mod_occ_lambda_3,mod_occ_lambda_2,mod_occ_lambda)
summary(mod_occ_lambda_5)


##### REPEAT DIVERSITY (Shannon Index) ######
#Test with simple lm
mod_input_SI<-ALL_DATA_palms_GS_vegan_pct[,c(1,2,32,33,35:54)]
mod_input_SI[,-c(1,2)]<-apply(mod_input_SI[,-c(1,2)],2,as.numeric)
mod_input_SI_fin<-mod_input_SI[,-c(1,2)]
colnames(mod_input_SI_fin)
dim(mod_input_SI_fin)


#How to normalise richness (response variable)
shapiro.test(mod_input_SI_fin$Shannon_Index)
hist(mod_input_SI_fin$Shannon_Index) # seems fine untransformed


mod_1_SI<-lm(Shannon_Index~., data=mod_input_SI_fin)
summary(mod_1_SI)

mod_2_SI<-step(mod_1_SI)
summary(mod_2_SI) #Cool, seems interesting.


#Let's try in a phylogenetically-corrected fashion
#PGLS of genome size vs repeat class
mod_input_SI_unique<-mod_input_SI[-which(duplicated(mod_input_SI$Tree_name)),c(1,3,4,5,8,12,13,14,18,19,20)]
colnames(mod_input_SI_unique)
dim(mod_input_SI_unique)

pairs(mod_input_SI_unique[,-1])

pgls_dat_SI<-comparative.data(data=mod_input_SI_unique, phy=phyclean, names.col="Tree_name",vcv=TRUE)
print(pgls_dat_SI)

mod_SI_lambda<-pgls(Shannon_Index~Temperature_Annual_Range_BIO5_BIO6+Mean_Temperature_of_Driest_Quarter+Precipitation_of_Driest_Month+Mean_Temperature_of_Wettest_Quarter+Precipitation_of_Wettest_Month+Precipitation_Seasonality_Coefficient_of_Variation+Genome_occupancy+Genome_size, data=pgls_dat_SI, lambda = 'ML', bounds = list(lambda = c(0.0005, 1))) #lambda branch transformation
summary(mod_SI_lambda)

mod_SI_lambda_2<-update(mod_SI_lambda,~. -Temperature_Annual_Range_BIO5_BIO6, lambda = 'ML', bounds = list(lambda = c(0.0005, 1))) #lambda branch transformation
summary(mod_SI_lambda_2)
mod_SI_lambda_3<-update(mod_SI_lambda_2,~. -Precipitation_of_Driest_Month, lambda = 'ML', bounds = list(lambda = c(0.005, 1))) #lambda branch transformation
summary(mod_SI_lambda_3)
mod_SI_lambda_4<-update(mod_SI_lambda_3,~. -Mean_Temperature_of_Wettest_Quarter, lambda = 'ML', bounds = list(lambda = c(0.0005, 1))) #lambda branch transformation
summary(mod_SI_lambda_4)
mod_SI_lambda_5<-update(mod_SI_lambda_4,~. -Precipitation_of_Wettest_Month, lambda = 'ML', bounds = list(lambda = c(0.0005, 1))) #lambda branch transformation
summary(mod_SI_lambda_5)
mod_SI_lambda_6<-update(mod_SI_lambda_5,~. -Precipitation_Seasonality_Coefficient_of_Variation, lambda = 'ML', bounds = list(lambda = c(0.0005, 1))) #lambda branch transformation
summary(mod_SI_lambda_6)


AIC(mod_SI_lambda_6,mod_SI_lambda_5,mod_SI_lambda_4,mod_SI_lambda_3,mod_SI_lambda_2,mod_SI_lambda)
summary(mod_SI_lambda_5)
```
