## Supporting Information 3 for "The Ecology of Palm Genomes: Repeat-associated genome size expansion is constrained by aridity": README_Supp_Info_3.txt

################# README for Supplementary Information 3 ################
# Repeatexplorer output analysis scripts from 'The Ecology of Palm Genomes: Repeat-associated genome size expansion is constrained by aridity'
# (R. J. Schley et al.) 


## NOTE, THESE SCRIPTS ARE NOT DESIGNED TO RUN 'OUT OF THE BOX', THEY REQUIRE SOME MODIFICATION ##


############################################################################
#### Analysing output from repeat explorer single (CLUSTERING) analysis ####
############################################################################

When you get your output files from the RepeatExplorer2 clustering analysis, you will need to 
parse them and, following that, you will need to run those parsed results through an R script 
in which all of the PGLS modelling and analyses are undertaken.


1) So, first run '1_RepeatExplorer2_output_analysis_wrapper.sh'. This script:
	> Renames output files, so you know which species each repeat profile comes from
	> Adds read numbers and genome sizes to each output file name from a list of those two metrics for each species that you prepare before
	> Prepares each species' CLUSTER_TABLE.csv (which contains repeat profiles) for analysis with R. 
	  Part of this involves concatenating all CLUSTER_TABLE.csv files for all species to make manual checking of repeat assignment easier.	


2) Next comes the R script for analysis of the parsed output files. This is 'Analysing_repeatexplorer_output_final_analysis.R'.
	This script:
	> Cleans output files (renames incorrect species names, removes organellar reads)
	> Aggregates output files and remoulds them into a format that is appropriate for the R package Vegan. 
	  In this case, rows will be species (i.e., palm genomes) and columns represent repeat families
	> Attaches other data to these aggregate repeat profiles (e.g., precipitation data, genome size)
	> Performs analyses (calculates ecological metrics e.g. richness, diversity, performs PGLS analaysis, visualises data).
	
And you're done!
